# Genome-wide insights into population structure and host specificity of *Campylobacter jejuni*

**DOI:** 10.1101/2021.02.18.431648

**Authors:** Lennard Epping, Birgit Walther, Rosario M. Piro, Marie-Theres Knüver, Charlotte Huber, Andrea Thürmer, Antje Flieger, Angelika Fruth, Nicol Janecko, Lothar H. Wieler, Kerstin Stingl, Torsten Semmler

## Abstract

The zoonotic pathogen *Campylobacter jejuni* is among the leading causes of foodborne diseases worldwide. While *C. jejuni* colonises many wild animals and livestock, persistence mechanisms enabling the bacterium to adapt to host species’ guts are not fully understood. In order to identify putative determinants influencing host preferences of distinct lineages, bootstrapping based on stratified random sampling combined with a *k-mer-*based genome-wide association was conducted on 490 genomes from diverse origins in Germany and Canada.

We show a strong association of both the core and the accessory genome characteristics with distinct host animal species, indicating multiple adaptive trajectories defining the evolution of *C. jejuni* lifestyle preferences in different ecosystems. Here, we demonstrate that adaptation towards a specific host niche ecology is most likely a long evolutionary and multifactorial process, expressed by gene absence or presence and allele variations of core genes. Several host-specific allelic variants from different phylogenetic backgrounds, including *dna*E, *rpo*B, *ftsX or pyc*B play important roles for genome maintenance and metabolic pathways. Thus, variants of genes important for *C. jejuni* to cope with specific ecological niches or hosts may be useful markers for both surveillance and future pathogen intervention strategies.

## Introduction

*Campylobacter jejuni* is a bacterium isolated from human patients suffering from acute gastroenteritis [1]. The species is regarded as a common resident among the gut microbiota of many wild and agriculture-associated animals [2], especially birds, poultry and cattle [3, 4]. Contamination of (chicken) meat, water, raw-milk and other food products along the food production chain is therefore the most attributable factor of diarrheal disease caused by *C. jejuni* in humans [1,4–6].

Previous research using multilocus sequence typing (MLST) of *C. jejuni* from different origins showed that specific sequence types (STs) were frequently associated with a particular host species [15]. While STs belonging to the clonal complexes (CC)-42 and CC-61 are common among *C. jejuni* of cattle and/or other ruminate origins, STs belonging to CC-257, CC-353 or CC-1034 are regarded as chicken-specific [16–18]. Isolates belonging to STs sharing a clonal complex such as CC-21, CC-45 or CC-48 commonly occur in samples of multiple host species, indicating the ability of these phylogenetic lineages to rapidly switch between different (intestinal) conditions, and, therefore, representing a typical host-generalist lifestyle [19]. Factors influencing adaptation of *C. jejuni* to certain host species, especially to poultry and cattle, were an important focus of *Campylobacter* research over the last decade [20–22]. In recent years, novel bioinformatic methods and tools such as genome-wide association studies (GWAS) proved their potential to identify genetic factors promoting host adaptation and/or pathogenicity in *C. jejuni* [21–25]. For instance, accessory genes encoding factors involved in the bacterial vitamin B5 biosynthesis pathway were found to be associated with cattle and its typical diet [21], while proteins enhancing iron acquisition abilities of the bacteria during infection were harboured by isolates from human clinical samples [24].

Most of the GWAS have been predominately focused on the variable set of genes commonly addressed as accessory genome. However, changes among (essential) core genes (i.e. basic cellular and regulatory functions) within the *C. jejuni* population may reflect adaptation towards a particular bacterial lifestyle as well.

Core genome alterations are thought to play an important role in overcoming specific host-associated intestinal stress conditions [26, 27], while other alterations may enable certain *Campylobacter* lineages to cope with colonisation inhibitors or even diets associated with gastrointestinal tracts of a much broader range of host species [28]. A recent GWAS study indicated that the worldwide intensified cattle farming for meat production was accompanied by a timeline of genomic events enhancing host adaptation of certain *C. jejuni* lineages to cattle [29].

The aim of this study was to generate in-depth insights into the current population structure of *C. jejuni* by using high resolution of whole genome sequencing and a stratified random sampling approach combined with GWAS considering all nucleotide substrings of length *k* (*k-mers*) to investigate host adaptation niche gene associations and outbreak potential associated with host-specific *C. jejuni* lineages.

## Material and Methods

### Strain selection and genome sequencing

A uniform stratified random collection comprising 324 *C. jejuni* isolates obtained from samples of four different species, including human (n=96), chicken (n=102), cattle (n=98) and pig (n=28). The original samples were collected in 16 different federal states in Germany, between 2010 and 2019. Isolates from healthy and diseased animals as well as human clinical isolates were included (Table S1). The animal-derived isolates were provided by the National Reference Laboratory for *Campylobacter* at the German Federal Institute for Risk Assessment (BFR) and the Institute of Microbiology and Epizootics (IMT) at Freie Universität Berlin, while the human-derived isolates were provided by the National Reference Centre for *Salmonella* and other Bacterial Enterics at the Robert Koch Institute (RKI). *C. jejuni* is rarely isolated from porcine, therefore porcine-derived isolates were limited. In order to limit spatial and temporal effects, the set of genomes investigated here was complemented by whole genome data of further 166 isolates from a Canadian study which included *C. jejuni* from cattle (n=39), chicken (n=12), human clinical cases (n=40), environmental (n=54) and other animal (n=21) origins [24]. The original purpose of the Canadian study was to identify diagnostic markers which can be used for rapid screening approaches to detect *C. jejuni* subtypes [24]. The complete list of all 490 genomes, including available metadata such as sample origin/source and baseline typing data such as ST is provided in Table S1. Detailed protocols used for whole genome sequencing (WGS) are provided as supplementary material. Illumina raw read data sequenced for this study is available at the National Center for Biotechnology Information (NCBI) under Bioproject ID PRJNA648048. Furthermore we included the strain BfR-CA-14430, available at NCBI under the accessory numbers CP043763.1 and CP043764.1, already published as a representative *C. jejuni* genome by the zoonosis monitoring program of Germany [30].

### Assembly and annotation

The Illumina paired-end reads were adapter-trimmed by Flexbar v.3.0.3 [31] and corrected using BayesHammer [32]. The *de novo* assembly was performed using SPAdes v3.11.1 [33] with default settings. All genomes were annotated by Prokka v1.13 [34] employing a customized database which consist of 137 complete annotated reference genomes provided by NCBI as described before [30].

### Multilocus sequence type (MLST) analysis

*In silico* MLST was carried out on seven housekeeping genes (*aspA, glnA, gltA, glyA, pgm, tkt, uncA*) as described by Dingle et al. [35]. This was done with the BLAST-based tool “mlst” (https://github.com/tseemann/mlst) based on the *Campylobacter jejuni* / *coli* database of pubmlst [36]. Obtained MLST profiles were then used to calculate a minimum spanning tree by MSTree V2 that was visualized with GrapeTree [37].

### Pan-genome and phylogenetic analyses

Open reading frames (ORFs) predicted by Prokka were subsequently used as input for Roary v3.12.0 [38] to calculate the pan-genome size and core genome alignment using default settings. The resulting alignment was used to calculate a maximum likelihood-based phylogeny with RAxML v.8.2.10 [39] with 100 bootstraps under the assumption of the gtr-gamma DNA substitution model [40]. ClonalFrameML v1.11 [41] was used to correct for recombination events and phylogenetic groups were identified with Bayesian Analysis of Population Structure (BAPS). Here, we used BAPS with hierarchical clustering that was implemented in the R packages RhierBAPS v1.0.1 [42]. Grouping of the accessory genome was further analysed by t-distributed stochastic neighbour embedding (t-SNE) [43].

### Recombination analysis

BratNextGen [44] was used to reconstruct putative recombination events based on the analysis of the core genome alignment of our selection comprising 490 *C. jejuni* genomes. Parameter estimation was performed based on 20 iterations and significant recombinations (p-value ≤0.05) were obtained using permutation testing with 100 permutations executed in parallel.

### Genome-wide association study (GWAS)

In order to perform an in-depth analysis of genomic alterations possibly associated with host specificity, pyseer v.1.1.2 [45] was used for GWAS based on variable-length *k-mer* composition (9 to 100 base pairs) for all 490 genomes. *K-mers* significantly representing distinct isolate origins (human, cattle, chicken or pig) were further mapped by bwa v0.7.17 [46] against selected reference genomes from this study set in order to identify putative origin-specific factors, genes and consecutive gene loci.

In order to reduce the false positive rate of the GWAS and account for highly unbalanced groups, we employed a bootstrapping approach. Further details can be found in the supplementary material.

The consequential set of genes was further analysed considering functional annotations and metabolic pathways using EggNog v.4.5.1 [47, 48].

### *C. jejuni* lifestyle classification

In order to facilitate statistical comparison, we adapted a definition from Shepard et al. [49] and defined a set of closely-related *C. jejuni* lineages as host-specific if ≥ 50% genomes building the respective BAPS cluster were associated with isolates from a specific animal origin (e.g. cattle, chicken) while each of the other isolate origins contributed less than 10% in the BAPS cluster. Potential host-generalist lineages were assumed when more than 25% of the clustering genomes represented in the corresponding BAPS cluster were from *C. jejuni* of human clinical cases while at least two further animal origins account for more than 10% of the remaining genomes, respectively.

## Results

### *C. jejuni* core and accessory genome analysis

Here we report on 490 genomes of *C. jejuni* isolated from samples of animal, human and environmental origins from two distinct continents. The average size of the *C. jejuni* genomes was 1 690 635 bp. We identified 1 111 core genes that covered 60% of the average *C. jejuni* genome size, while a set of additional 7 250 genes was identified in at least one of the genomes under consideration and therefore assigned to the accessory gene content.

### Core and accessory genome: phylogenetic structure and organisation of the *C. jejuni* population

The phylogenetic representation of the 490 core genomes showed 15 distinct phylogenetic branches (1-15) that have been confirmed by BAPS clustering (Figure 1). BAPS clusters identified here, which comprised of more than 15 *C. jejuni* genomes, were further evaluated according to their respective CCs, original sample source and lifestyle classification (Table S1).

**Figure 1:**
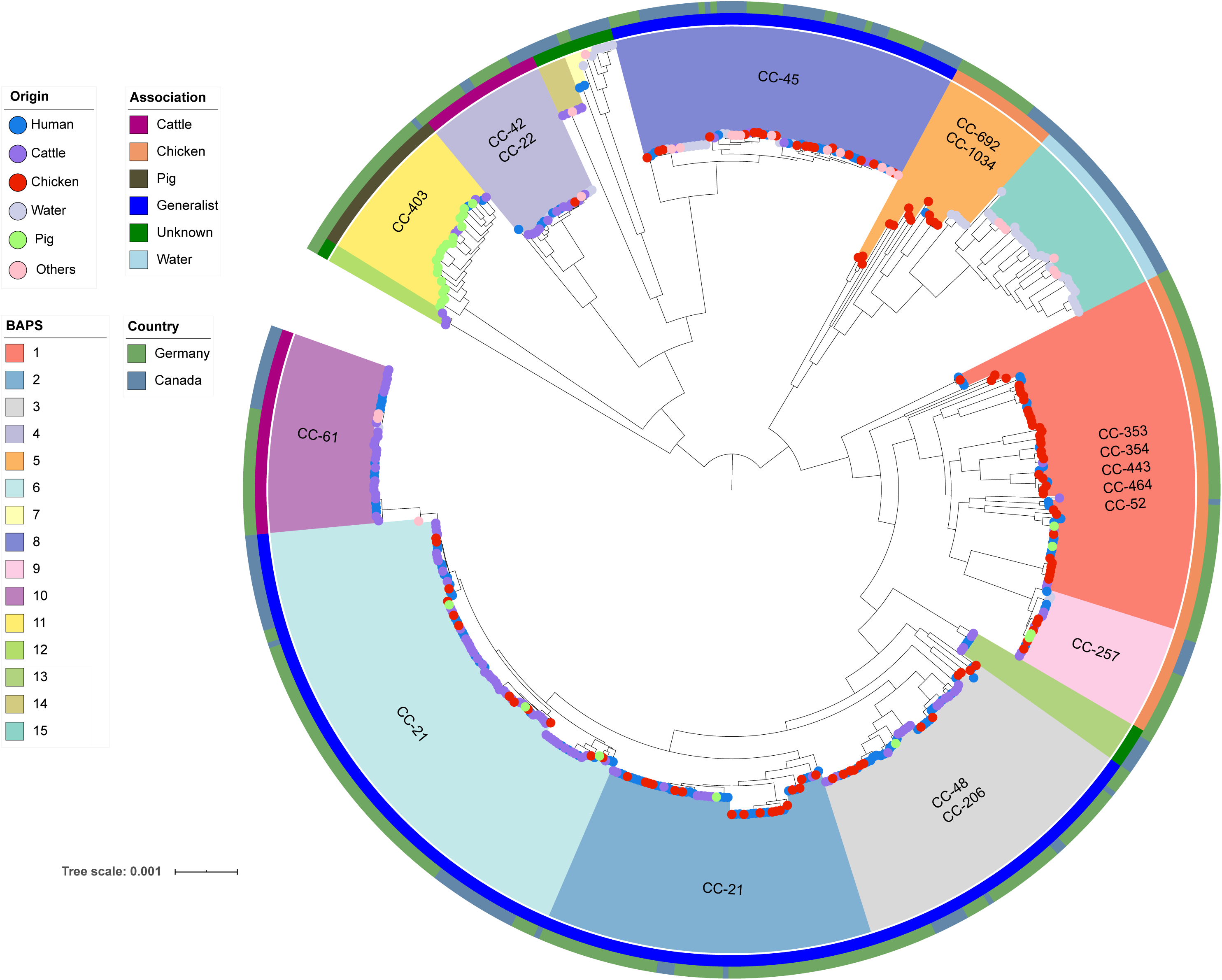
Population structure of *C. jejuni* based on the core genome alignment with BAPS clusters and clonal complexes colour-coded in the inner ring; Lifestyle preferences of the genomes coded in the second -ring; and country of genome origin described in the outer ring. The leaves are coloured by the origin of each sample.

**Table 1.**
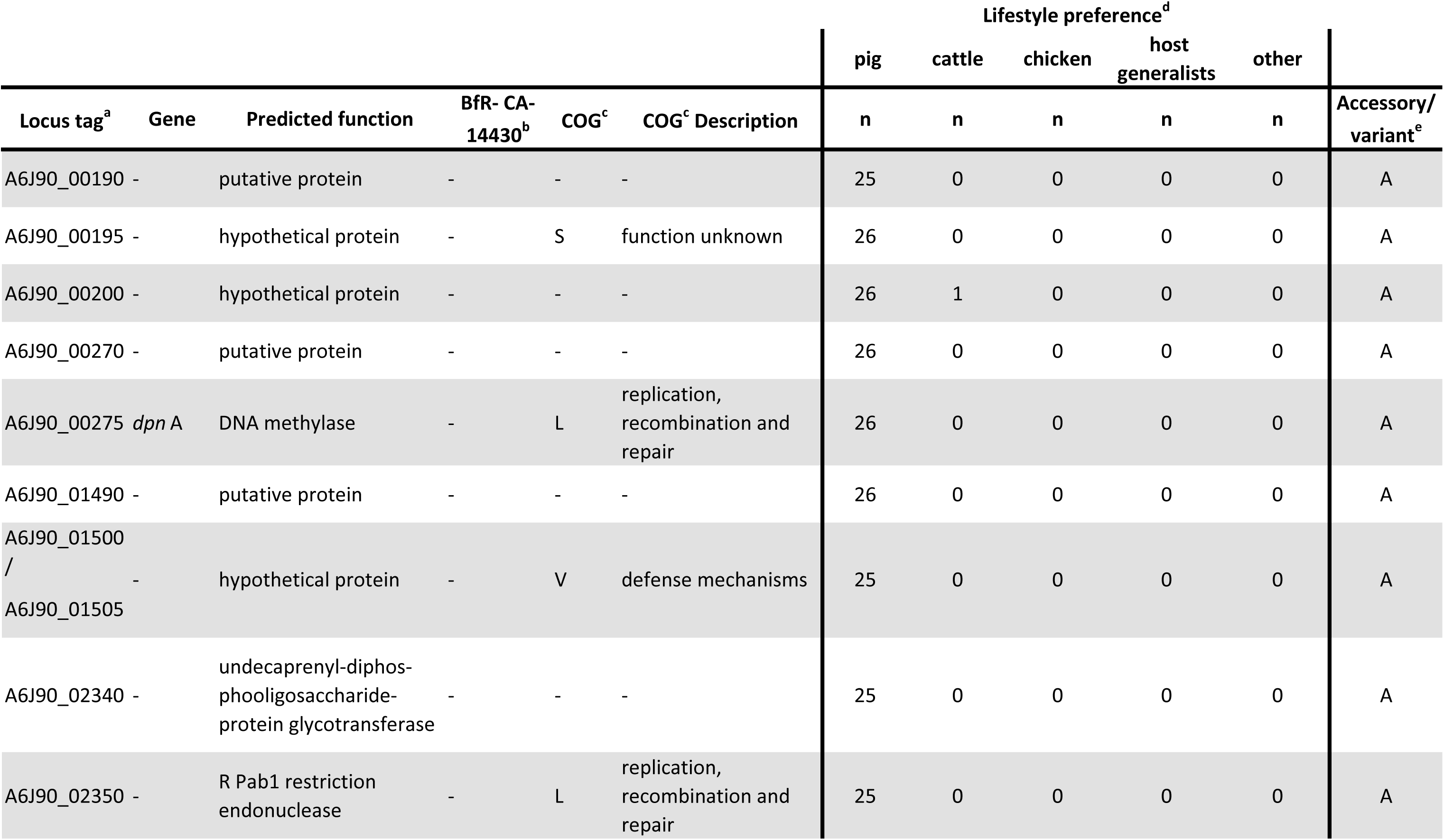
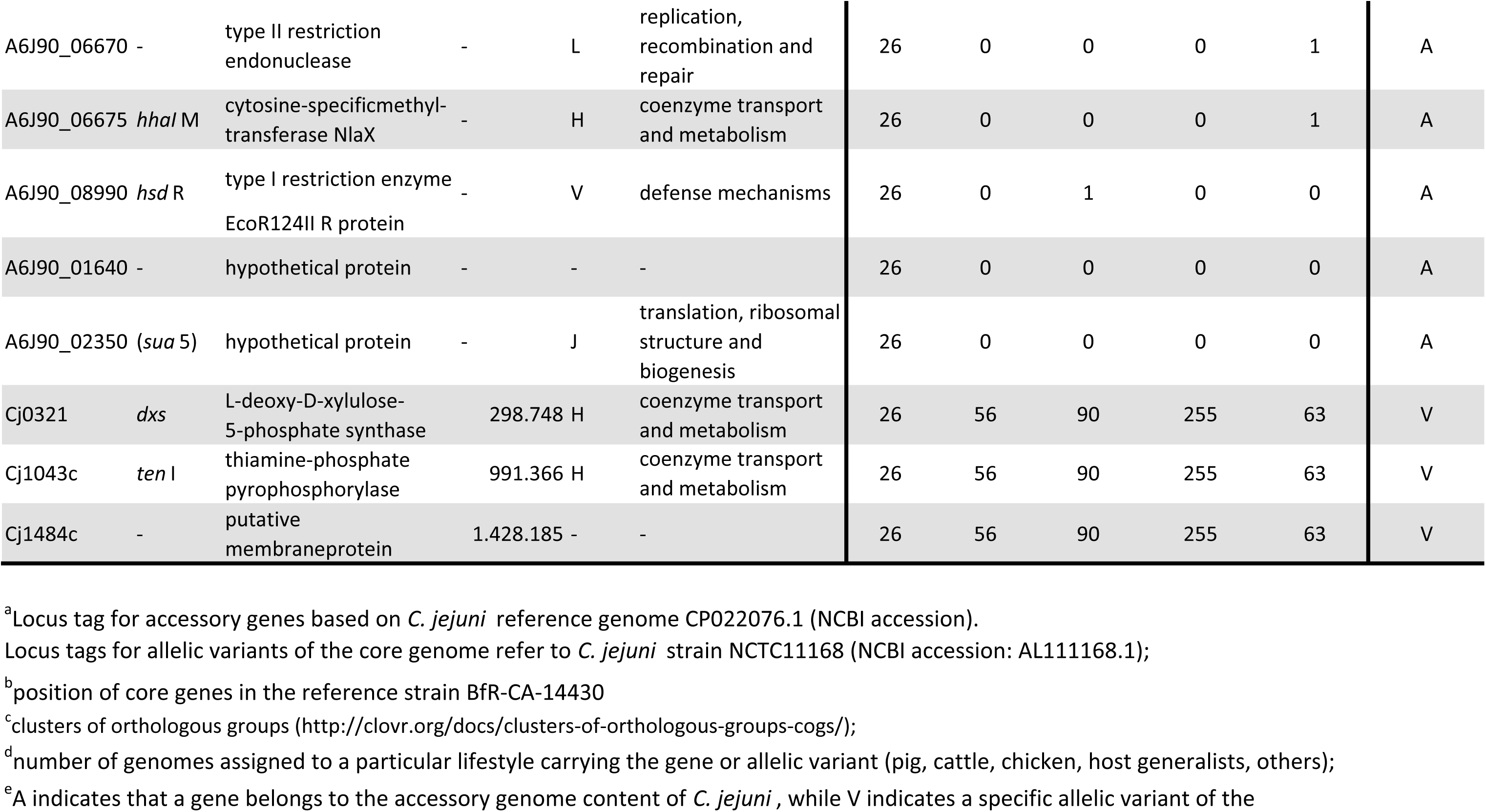
Selected pig-associated accessory genes and allelic variants of the *C. jejuni* core genome content.

For the original sample sources of the *C. jejuni* genomes investigated here, the relative proportion and absolute distribution for each of the BAPS clusters are visualised in Figure 2a and supplementary Figure S1a. We identified a close phylogenetic relationship between genomes of BAPS cluster 5 representing the origin chicken with those of BAPS cluster 15 representing waterborne environmental *C. jejuni* (Figure 1).

**Figure 2.**
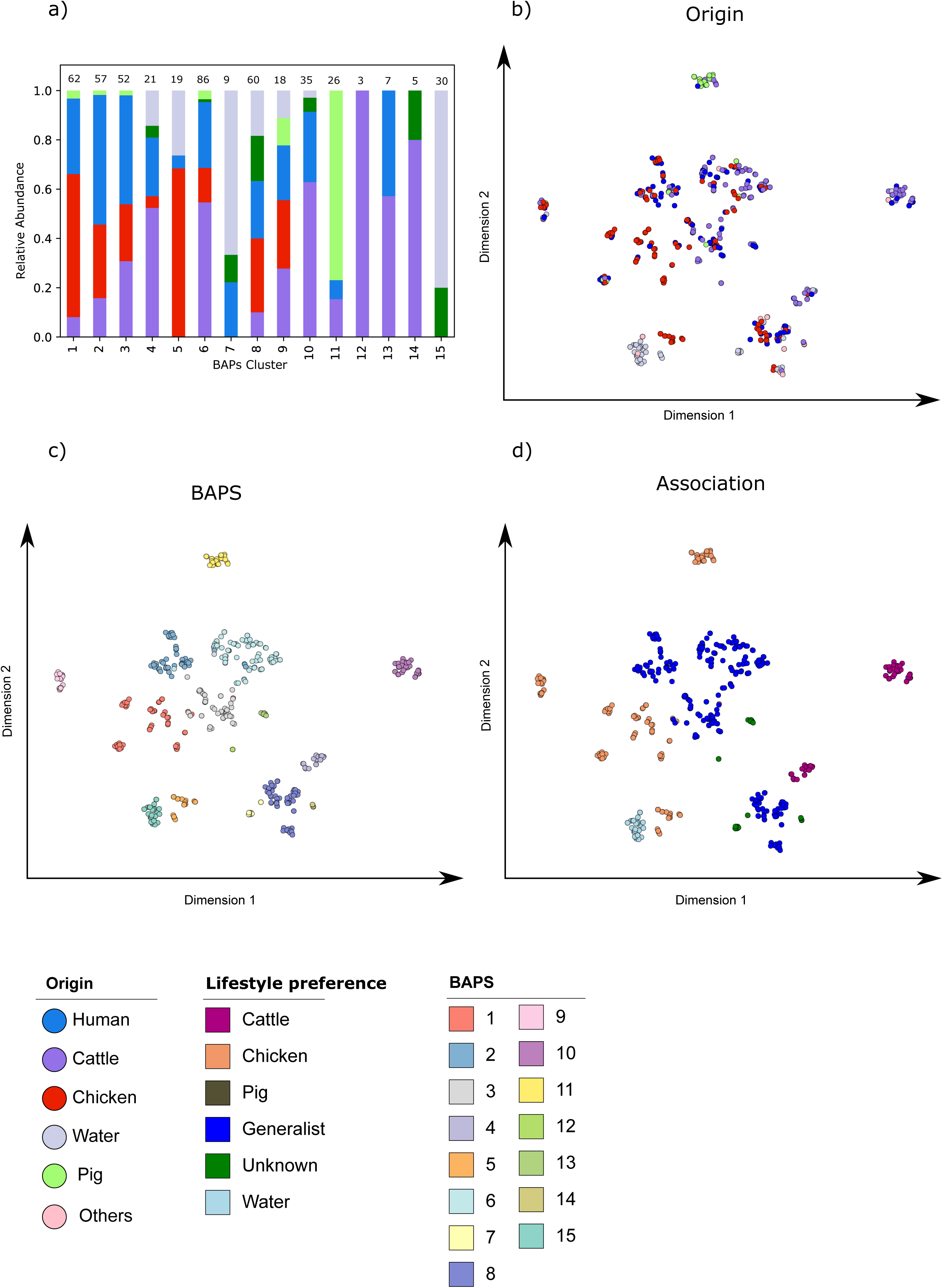
Relative distribution of sample origin among BAPS clusters and t-SNE plots of the accessory genome profile. a) shows the relative proportion of sample origins within the BAPS cluster that are later used for the stratified random sampling approach. b), c) and d) show t-SNE plots in the 2-dimensional space of the accessory genome profiles. The colours included in the legend represent the sampling source, the BAPS clusters and the lifestyle preference are included in the legend.

The genomes of BAPS cluster 15 and those of BAPS clusters with genomes from less than 15 isolates were not analysed with respect to their lifestyle preference and were therefore used as a control group in our study.

The lifestyle preference of each major BAPS cluster was determined and subjected to an internal assessment: As shown in Table S2, our assignments are generally concordant with lifestyle preferences reported for frequently occurring lineages such as CC-353, CC-354, CC-443, CC-464 and CC-52 (chicken), CC-42 and CC-61 (cattle) and CC-403 (pig). We also identified the probable lifestyle classification for the CC-22 lineage (cattle) and for isolates belonging to ST-2274 (chicken) (Table 1 and Table S2). Of note, the *C. jejuni* genomes associated with CC-21, CC-45 and CC-48 fulfilled the criteria for host-generalist lineages (Table S2).

Overall, the genomes assigned to individual BAPS clusters consisting of lineages considered as either host-specific for cattle (BAPS 4; including CC-42 and CC-22; BAPS 10, CC-61) or pigs (BAPS 11, CC-403) showed generally a less diverse population structure than those assigned to clusters associated with the host chicken (e.g. BAPS 5, including CC-1034 and CC-692). The distinct BAPS clusters comprising of host-generalist lineages (BAPS 8, including CC-45 and CC-283; BAPS 2, CC-21; BAPS 6, CC-21) showed a more diverse population structure (Figure 1).

Our core genome-based phylogenetic analysis further revealed that cattle-related BAPS cluster 4 lineages (including CC-42 and CC-22) were more closely related to host-generalist lineages of BAPS cluster 6 with CC-21 than to other cattle-related lineages, for instance those of BAPS cluster 10 (Figure 1). This also holds true for the chicken-related phylogenetic background (Figure 1): While chicken-related BAPS cluster 1 was found being more closely related to BAPS cluster 6 of host-generalist lineage, BAPS cluster 5 showed less phylogenetic distance to BAPS cluster 8 (host-generalist lineage). These findings clearly reject the hypothesis of a common evolutionary background for host-specific lineages with respect to the host species represented here.

Minimum spanning trees based on MLST utilising BAPS cluster classification and lifestyle preferences are shown in the supplementary material (Figure S1). Finally, the accessory genome profiles of all genomes were visualised by t-SNE plots in Figure 2 b - d including sample origin, BAPS cluster and lifestyle preference. As expected, the overall population structure derived from the core genome is mirrored in the accessory genome content. Each BAPS cluster carries its unique set of accessory genes (Figure 2c) confirming the population structure based on BAPS. Also, *C. jejuni* genomes belonging to different BAPS clusters while sharing a particular lifestyle preference differ with respect to their accessory gene content (Figure 2d). This observation is supported, for instance, by the accessory gene content identified for the cattle-specific BAPS clusters 4 and 10 (CC-42 and CC-61) and the chicken-specific BAPS clusters 1, 5 and 9 (CC-354, CC-692, CC-257, etc.) (Figures 2c, 2d). Overall, BAPS clusters with a host-generalist lifestyle preference appear to have a broader gene pool within the accessory genome content than strains identified as host-specific.

### Recombination events in *Campylobacter jejuni* lineages

Recombination events that show more differences between taxa than expected by mutation-driven evolutionary processes alone were illustrated in Figure 3. Overall, CCs assigned as cattle- or pig-associated as well as those belonging to the group of host-generalists showed recombination profiles most likely resulting from intra-lineage genomic events. The pig-associated lineages of BAPS cluster 11 and the cattle-associated lineages of BAPS cluster 4 shared limited recombination patterns with other lineages and yielded a low recombination rate compared with other clusters, indicating the possible presence of lineage-specific recombination barriers (Figure 3). The cattle-associated genomes forming BAPS cluster 10 showed several recombination events which were also indicated in the host-generalist lineages assigned to BAPS clusters 2, 3 and 6 (Figure 3). However, the cattle-associated BAPS clusters 4 and 10 shared a single recombination site only. The host-generalist BAPS clusters 2, 3, and 6 were found being associated with more recombination events and some of these were shared by host-specific lineages, i.e. BAPS cluster 10 (cattle) and BAPS clusters 1, 5 and 9 (chicken), indicating genomic exchanges between these lineages. In addition, the analysis revealed that chicken-associated lineages (BAPS clusters 1, 5 and 9) were prone to trade off genetic material with each other and with host-generalist lineages (Figure 3).

**Figure 3:**
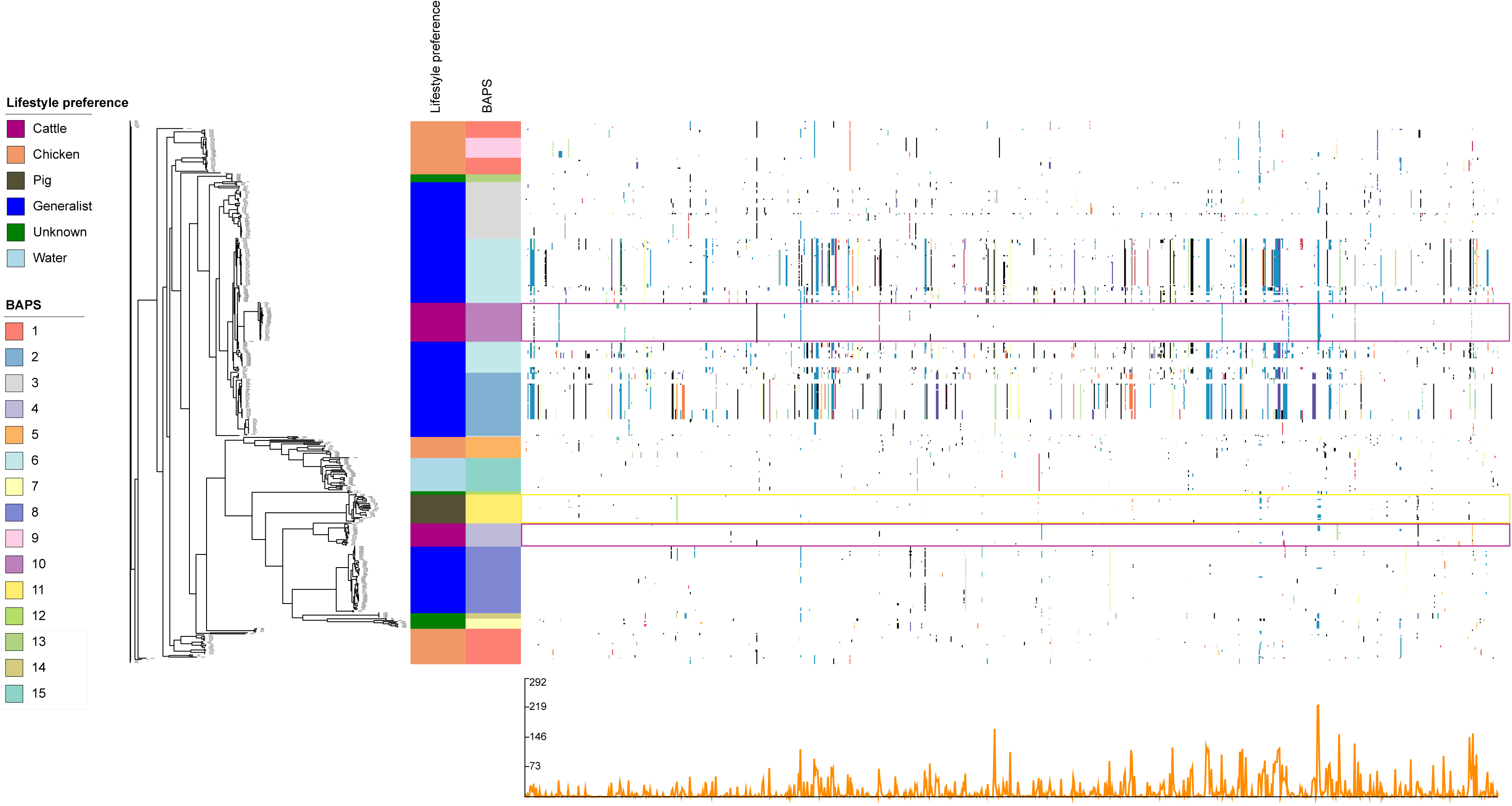
Recombination profile of the core genome alignment of 490 *C. jejuni* isolates calculated by BRATNextGen and visualized in Phandango. The left side shows the core genome phylogeny. The metadata provide information about lifestyle preferences (association) and BAPS clusters. Significant recombinations are marked by coloured dots and lines. Purple and yellow boxes highlight cattle- and pig-associated BAPS clusters, respectively. Presence of dot of the same colour across multiple isolates within a column represents acquisition of the same recombinant segment, otherwise colours are arbitrary. The line graph at the bottom presents recombination prevalence along the genome sequence.

### *In-Silico* identification of host-specific factors

After identifying significant *k-mers* using a consensus GWAS approach, the *k-mers* were mapped to an annotated reference genome in order to identify coding sequences of the genome known to promote a particular lifestyle preference of *C. jejuni* [45]. A visualization of the resulting genes with corresponding p-values and frequencies for the matching *k-mers* are provided in supplementary Figure S2.

Genes identified by *k-mers* in the genomes of *C. jejuni* isolates with lifestyle preferences in pig and cattle showed a denser distribution around the expected allele frequency than the results obtained for the genomes representing chicken- or host-generalist lineages (Figure S2).

The genes identified by our analysis included accessory genes present in a limited number of genomic backgrounds and allelic variants of the core genome content. We identified several variants of core genes supporting specific lifestyle preferences in *C. jejuni*. To further evaluate the putative host-specific importance of the allelic variants identified, genes under consideration have been checked for non-synonymous base changes by comparing their predicted amino acid (aa) sequences. Several of these predicted aa sequences can be linked to particular lifestyle preferences of *C. jejuni* isolates. Details for all loci and aa sequence variants identified are provided in the Tables S3 (cattle), S4 (chicken), S5 (pig) and S6 (host-generalists).

### Accessory genes and allelic variants of the core genome associated with *C. jejuni* lineages assigned as pig-specific

In the genomes belonging to BAPS cluster 11 (CC-403) we identified 21,681 *k-mers* which are significantly associated with the host pig. These *k-mers* mapped to 49 accessory genes and 78 allelic variants of the core genome (Table S5). Considering the accessory genes, 14 were exclusively found within *C. jejuni* genomes from pig hosts. (Table 1). Three accessory genes (A6J90_06670, A6J90_06675, A6J90_02350) belonged to transcription units encoding type II restriction modification systems (RM systems), while a further gene encodes the restriction subunit (R) of the host specificity determinant (*hsd*R; A6J90_08990) of a type I RM system. Additional 8/14 genes were annotated as hypothetical or putative proteins without any specific functional information available in NCBI GenBank (17.06.2020).

Considering the *k-mer* results for genes belonging to the core genome, nucleotide changes leading to actual effects with respect to host adaptation capabilities of certain lineages are difficult to pinpoint. Here, we noted alterations for the predicted aa sequences associated with the capability of *C. jejuni* to synthesize vitamins and enzyme co-factors such as TenI and Dxs (Figure 4a). In addition, the predicted aa sequence for Cj1484 was found to be altered (Figure 4a).

**Figure 4:**
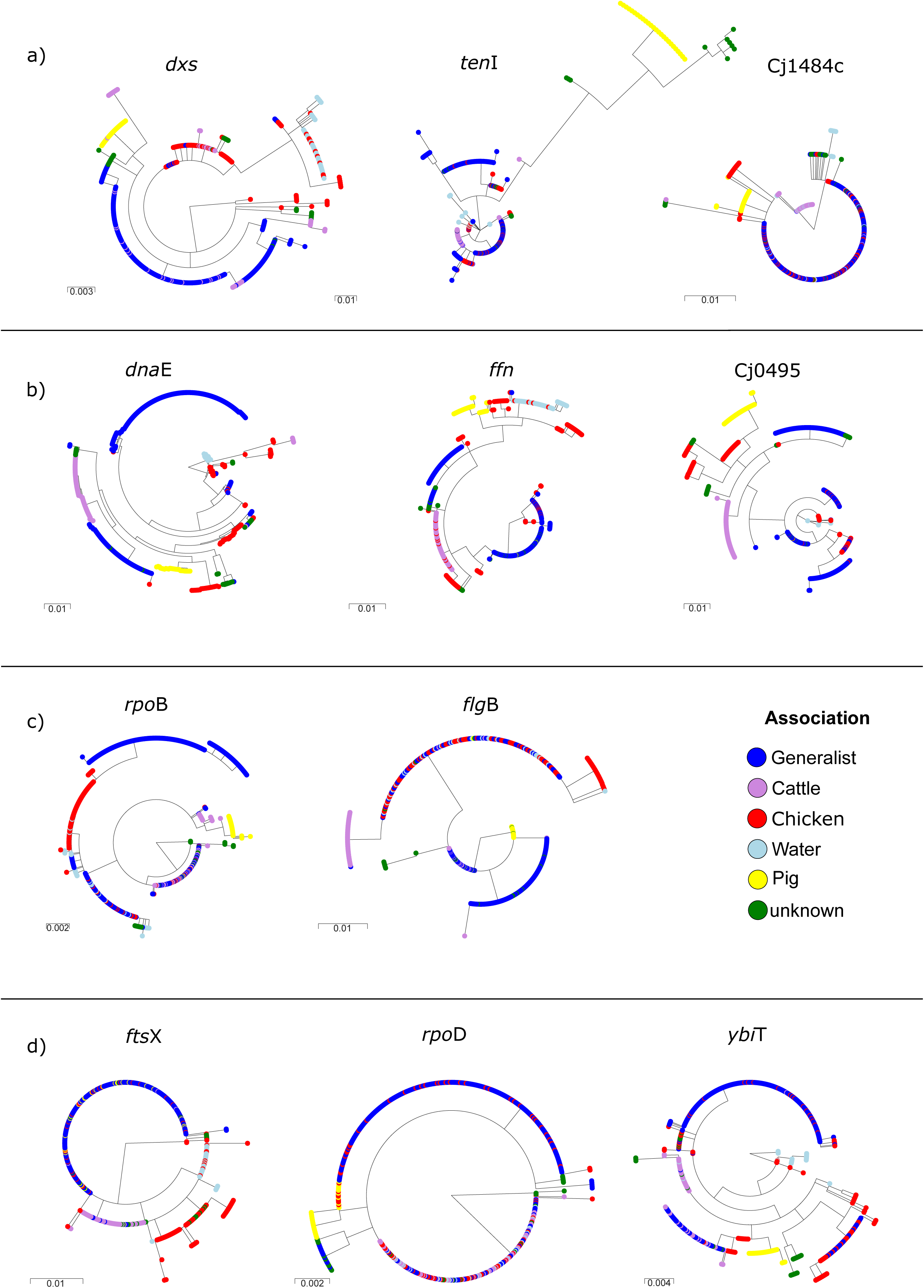
Phylogenetic tree of predicted amino acid sequence variants encoded by *dna*E, *ffh*, Cj0495, *rpo*B, *flg*B, *fts*X, *rpo*D, *ybi*T, *dxs*, *ten*I and Cj484c (selected from Tables 2-4) that show lifestyle associated variants (colour coded in legend) in different phylogenetic lineages originating from different genetic and geographic backgrounds (Figure 1).

**Table 2:**
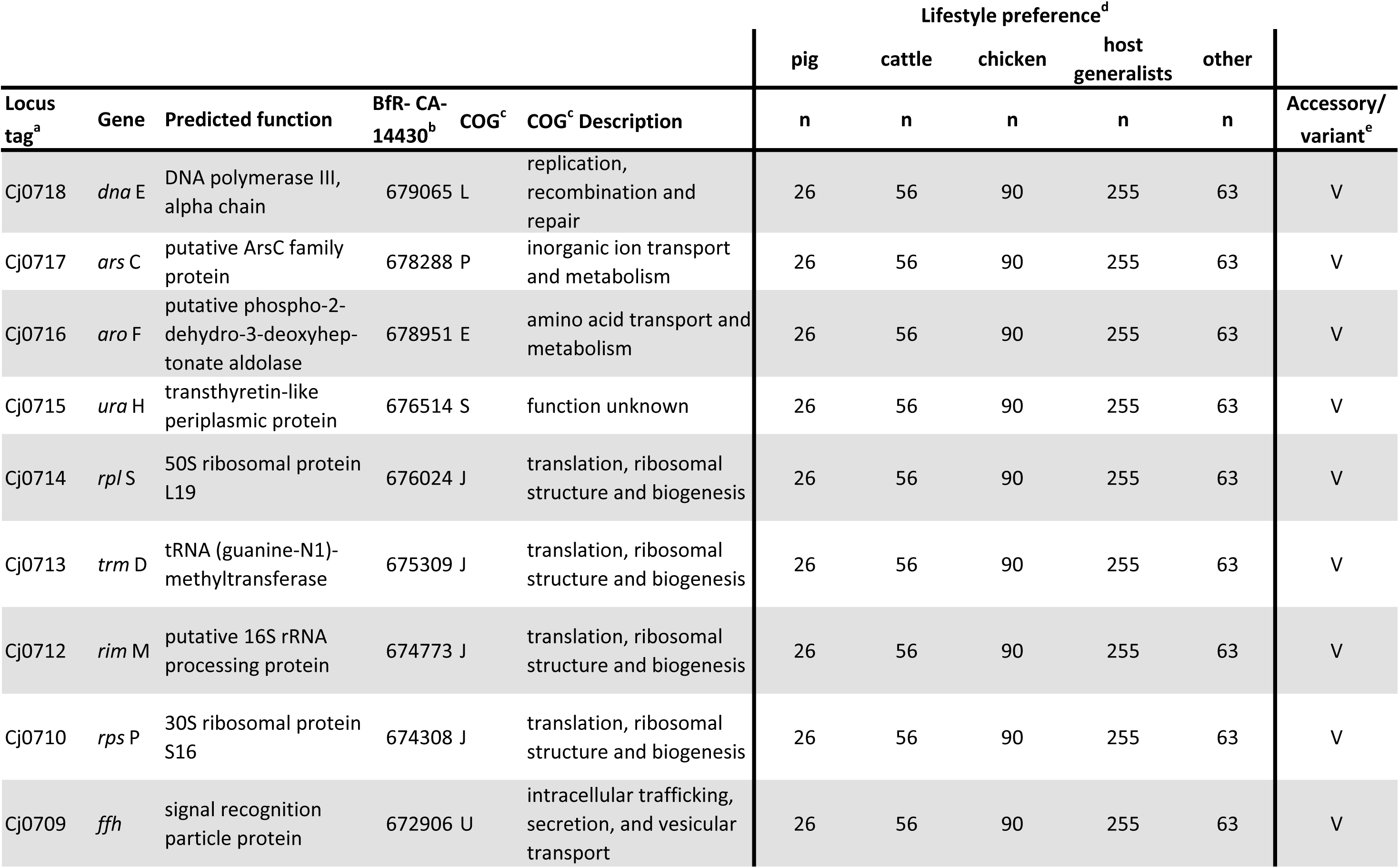
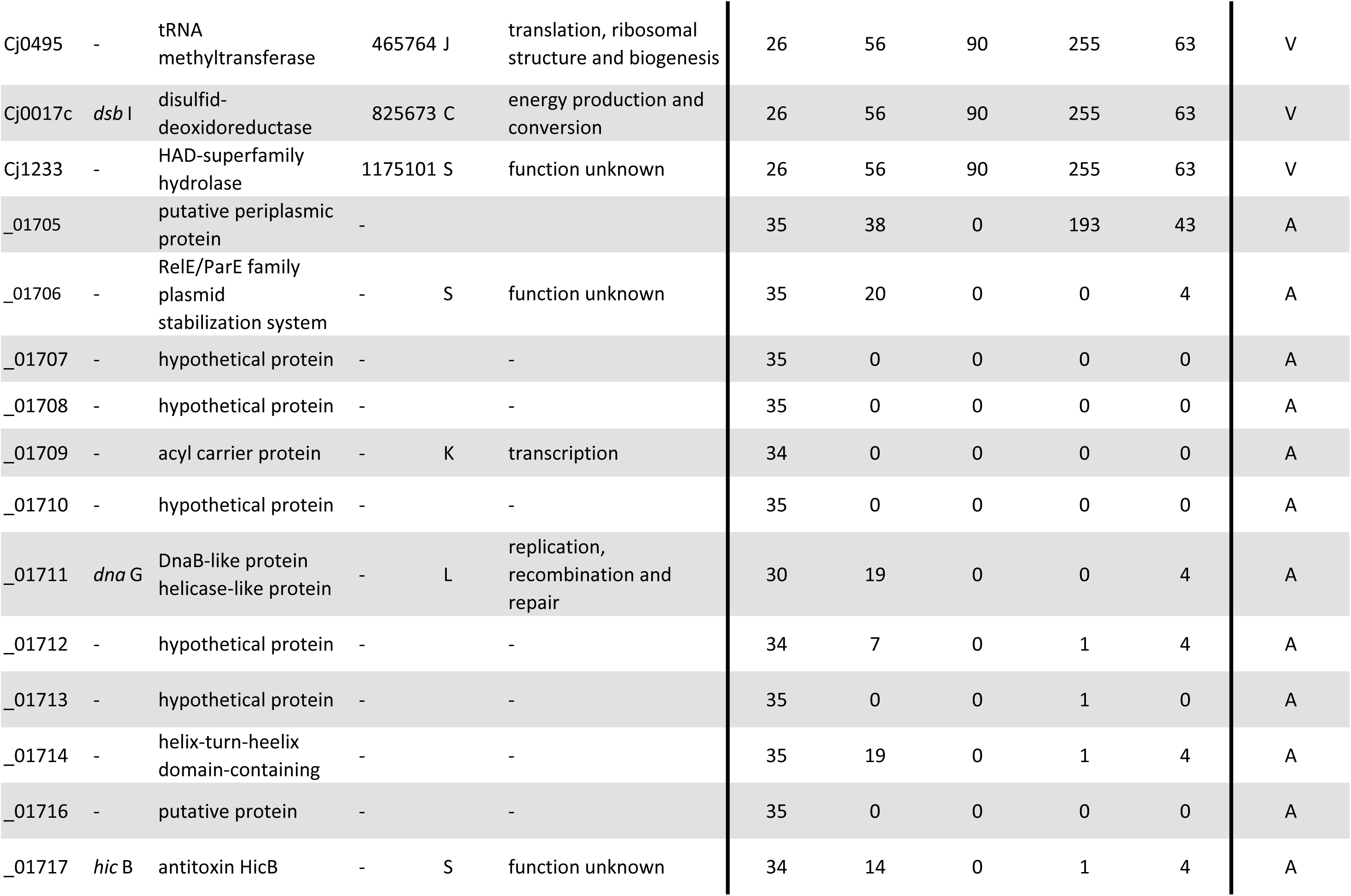
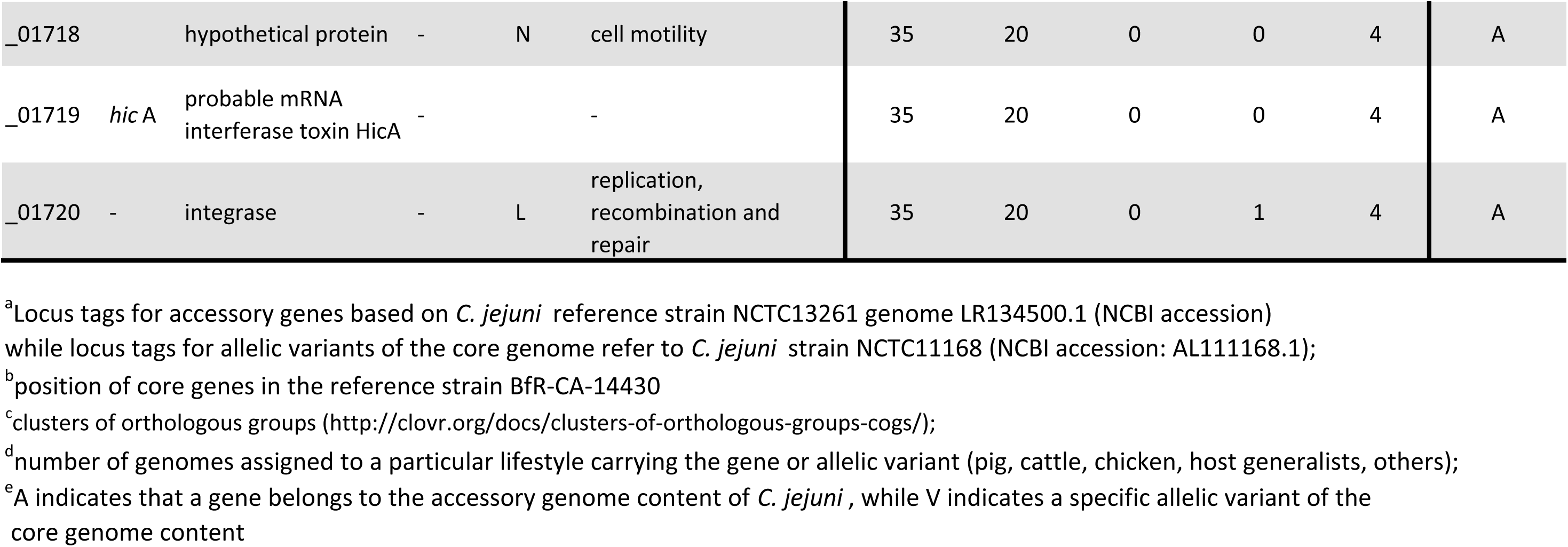
Selected accessory genes and allelic variants of the core genome content associated with the host cattle

### Accessory genes and allelic variants of the core genome associated with *C. jejuni* lineages assigned as cattle-specific

We further identified 66,491 *k-mers* for the cattle-associated genomes matching to 71 accessory genes and to 136 core gene variants (Table S3). According to our GWAS analysis, a particular accessory gene content which is representative for the lineages in both cattle-associated BAPS clusters (4 and 10) was not identified. However, 16 accessory genes were identified by *k-mers* significantly associated with CC-61 (BAPS cluster 10; Table S3). These genes belonged to a region of 9.9 kb size in *C. jejuni* (NCTC13261_01705 up to NCTC13261_01720). That particular locus contains 16 open reading frames encoding a HicA-HicB toxin/antitoxin system inhibiting the transfer of mRNA in case of nutrient limitation, a protein known to be involved in extracytoplasmatic stress response (YafQ) and regulatory protein RepA for plasmid DNA repair (Table S3).

Within the core genome we identified a 9.7 kb locus of 9 adjacent genes (Table 2) that encode for a ribosomal complex. While the allelic variants (non-synonymous substitutions) *dna*E and *ffh* (Figure 4b) were identified as cattle-specific, identical variants of *ars*C, *aroF*, *ura*H, *rpl*S, *trm*D, *rim*M and *rps*P were identified in host-generalist BAPS cluster 8, too. However, for the genes *ura*H, *arsC*, *rpl*S and *rps*P, detected SNPs lead to synonymous changes only, indicating their biological importance as conserved housekeeping genes within the *C. jejuni* lineages investigated here.

Additional non-synonymous, cattle-specific allelic variants were also identified on independent positions within the genome, including the alleles Cj0495(Figure 4b), *dsb*I and Cj1233 (Table 2).

### Accessory genes and allelic variants of the core genome associated with *C. jejuni* lineages assigned as chicken-specific

In comparison to the lineages associated with cattle, pig or even the host-generalists, chicken-associated lineages showed the broadest phylogenetic diversification in our study, mirrored by multiple lineages and CCs (Figure 1), including enhanced divergence within a specific CC (CC-353 or CC-1034). Accordingly, this particular heterogeneity resulted in less host-specific signatures. The 5 712 chicken-associated *k-mers* identified by our GWAS analysis cover 17 accessory genes and 25 core gene variants (Table S4). A gene for a TraG-like protein of the type IV secretion system [50] was detected among the accessory genomes in 59/90 chicken-associated genomes (Table 3). TraG-like proteins are known to play a crucial role in the conjugative transfer of plasmids [51]. Additionally, two genes for putative proteins of unknown function are carried by 66 and 68 of the chicken associated strains, respectively (Table 3).

**Table 3.**
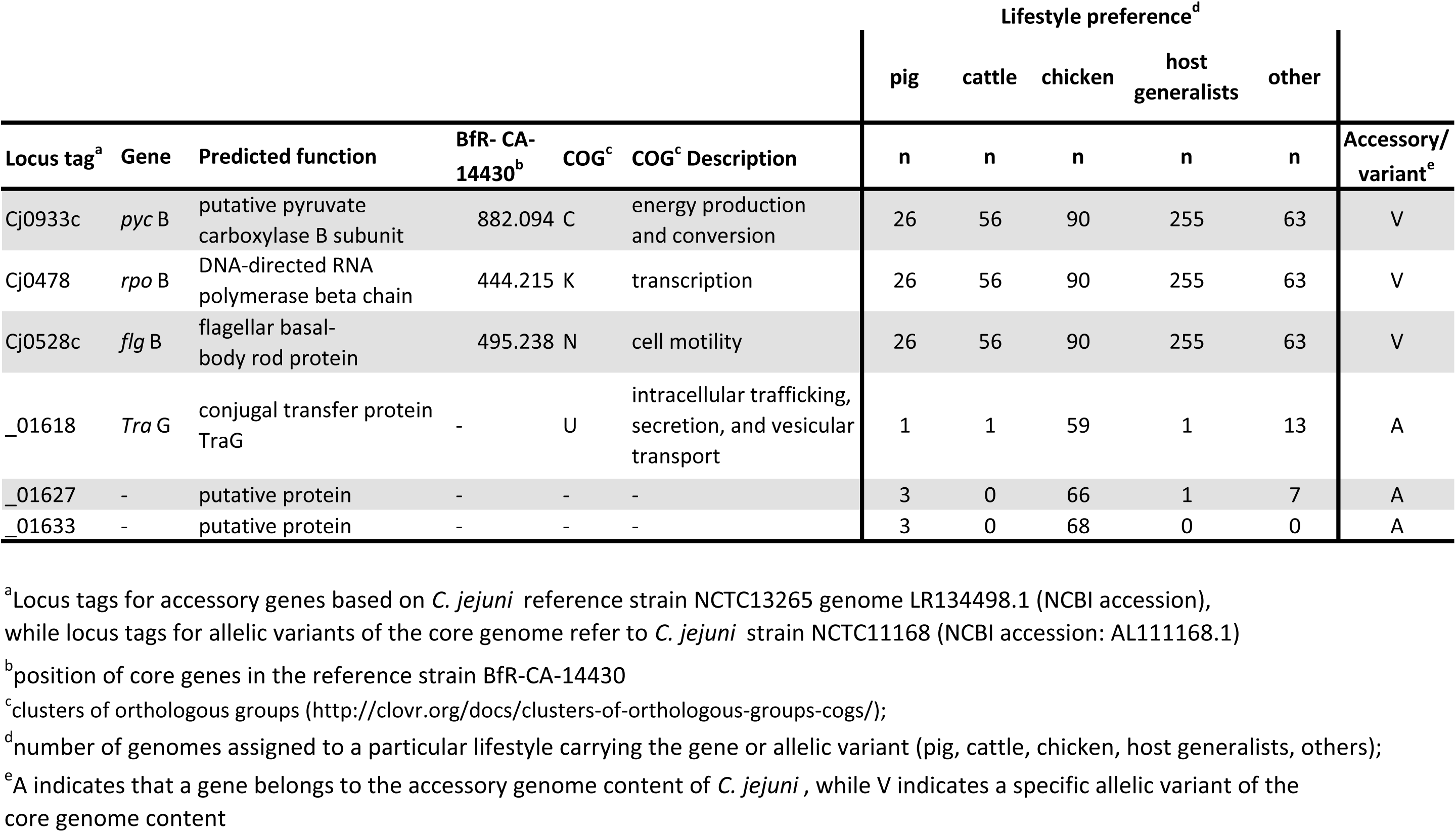
Selected accessory genes and allelic variants of the core genome content associated with the host chicken

**Table 4.**
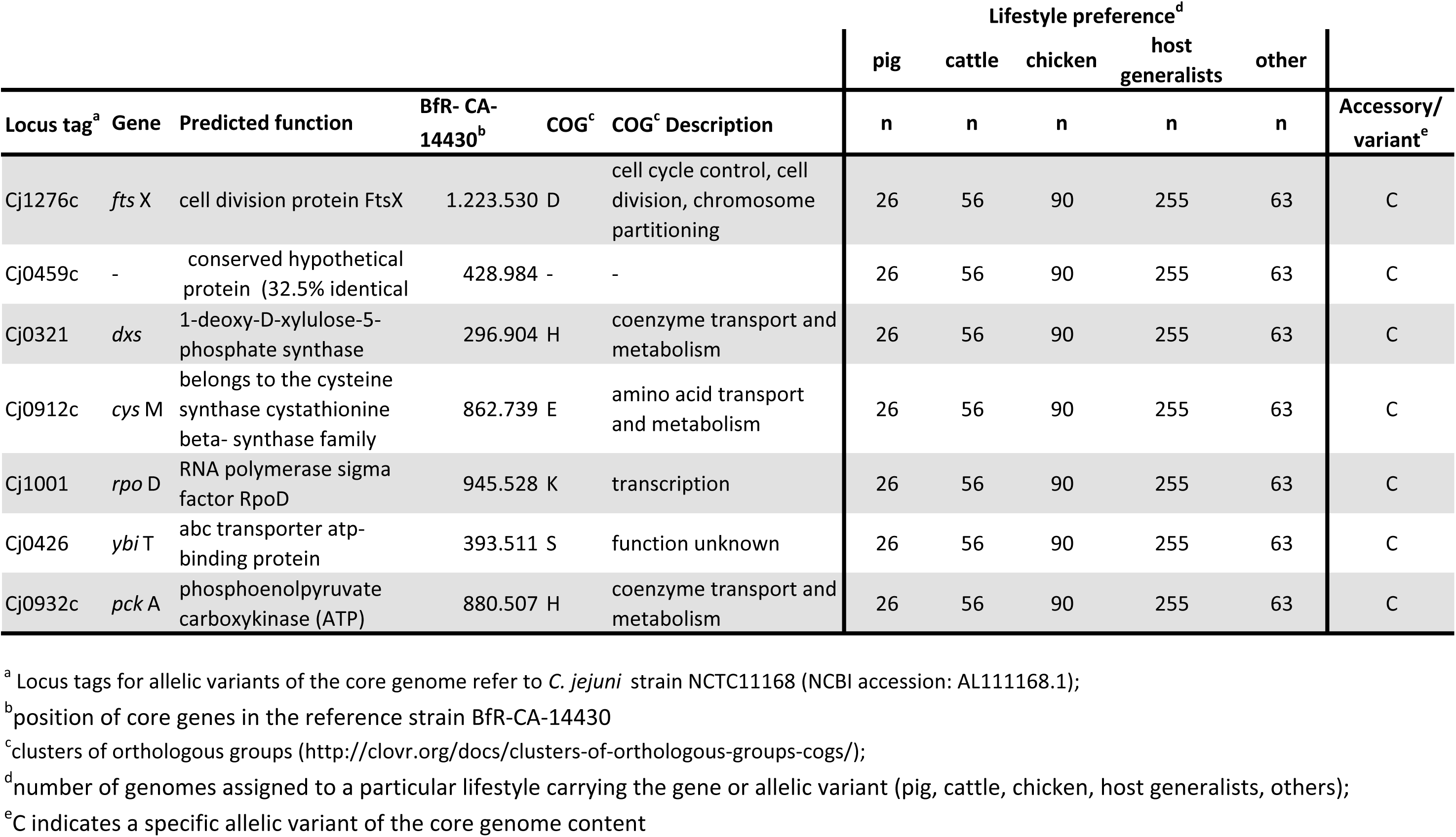
Selected allelic variants of the core genome content associated with host-generalists

Like the cattle-associated lineages, chicken-associated genomes carry host-adapted allelic variants (Table 3). The allele encoding a specific aa variant of *rpoB* was identified in most of the genomes in all three chicken-associated BAPS clusters (Table 3, Figure 4c). The gene variant encoding FlgB (Figure 4c) is identical in BAPS clusters 1 and 5 (chicken) and the host-generalist BAPS cluster 2 (CC-21). Furthermore, a very closely related aa variant was identified in BAPS cluster 9 (chicken) as well. Additionally, the same allelic variant of the *pycB* gene is carried by most genomes of BAPS clusters 1 and 9 (Table 3).

### Independent Adaptation of Host-Generalist Lineages

Considering the core genome phylogeny of the *C. jejuni* strains presented here, the host-generalist lineages of BAPS cluster 8 appear to have evolved from independent genomic backgrounds, while other host-generalist lineages, for instance those of BAPS clusters 2, 3 and 6, appeared to be linked to each other (Figure 1). In total, we have identified 37 339 *k-mers* which were mapped to 33 accessory genes and 87 core gene variants (Table S6). Accessory gene content exclusively associated with all host-generalist lineages was not identified by use of GWAS. A multitude of different allelic variants assigned to the core genome were identified for BAPS cluster 8 when compared with the genomes of the more closely related lineages of clusters 2, 3 and 6 (Table 4). Notably we also identified closely related variants for different core genes shared by all host-generalist lineages. These included *ftsX*, a gene involved in cell division, *ars*C, an arsenate reductase, further ribosomal genes (*rplS* and *rpsP*) and Cj0459c, known as a nicking endonuclease and purine-specific ribonuclease [52] (Table 4). While the ‘aa’ sequence encoded by *ftsX* shows a particular host-generalist-associated variant (Figure 4d), the ‘aa’ sequence determined by *ars*C, *rpl*S, *rps*P and Cj0459c are conserved in the *C. jejuni* population. Hence, *k-mers* identified for these coding sequences were associated with synonymous changes only. BAPS clusters 2, 3 and 6 harbour identical allelic variants for *dna*E and *ffh* (Figure 4b). The same is true for several other genes such as *dxs*, *cys*M and *pck*A (Table 4) that are broadly distributed across the *C. jejuni* genome and are involved in multiple metabolic pathways. Additionally, genes involved in transcriptional pathways such as *rpoD* and substrate transport functions like *ybi*T (Figure 4d) were identified.

## Discussion

We show how the recently emerging research field of bacterial GWAS was able to identify genetic signatures that play important roles for the host-specificity of *Campylobacter*. For each of the lifestyle preferences of *C. jejuni* investigated, we identified a broad set of allelic variants being associated with particular host-specific lineages from distantly related BAPS clusters, providing evidence for host-adaptive genetic signatures [53].

We also extended the scheme of lifestyle preferences based on MLST to a whole genome level by applying BAPS and identified 15 distinct phylogenetic clusters. The efficiency of the proposed approach to identify lifestyle preferences by assigning host-specific or host-generalist *C. jejuni* lineages was verified by performing a comparison of the predicted lifestyles. For instance, CC-42 or CC-61 (cattle), CC-354 or CC-692 (chicken) and CC-403 (mammalian/pig) lifestyle assignments were verified with previously published reports on these *C. jejuni* lineages [16,49,54]. Additionally, novel lifestyle preferences of distinct lineages, i.e. CC-22 (cattle-specific) and ST-2274 (chicken-specific), were identified using the definition described above.

*C. jejuni* isolates assigned to either chicken or host-generalist lineages showed a diverse population structure, as reported before [35]. Contrarily, we found *C. jejuni* genomes identified as cattle-specific (CC-42 and CC-61) or pig-specific (CC-403) were less diverse and more clonal. Previous studies assumed that the tight clonal structure of the cattle-associated lineages CC-42 and CC-61 resulted from a more recent onset of the colonization of cattle by *C. jejuni* and therefore may reflect a bottleneck event in its evolution [29, 53]. A similar host-adaptation process is possibly indicated by the limited diversity of CC-403 (pig-specific) assigned to BAPS cluster 11 in our study.

Genetic variation is known to be a pre-requisite to evolutionary change [55]. Since 2016, bacterial GWAS has advanced as a suitable method to identify genetic alterations associated with a phenotypical traits in large WGS datasets [56, 57], including studies on *C. jejuni* [21–24]. Acting like a “sieve”, genetic selection allows only a subset of mutations to persist and become an observable difference between genomes [55]. Allelic variants of *C. jejuni* core genes, independently acquired by different phylogenetic lineages leading to changes of known or predicted ‘aa’ sequences, likely reflect adaptation to a particular ecological niche and/or host [58, 59]. We have identified allelic variants of core genes which were clearly associated with the host species pig, cattle and chicken, even among distantly related BAPS clusters [BAPS 4 and 10 (cattle); BAPS 1, 5 and 9 (chicken)]. Further allelic variants (e.g. *ftsX* in CC-45 and CC-21) were identified as putative markers for host-generalist lineages. This observation is supported by the lack of notable recombination between CC-45 and CC-21, indicating that these variants occurred independently of phylogenetic background and geographic origin. Therefore, mutant selection leading to homoplasy would be the most reasonable assumption. More research on the subject, including isolates covering a broader time span is needed to gain further insight into the bacterial evolution of *C. jejuni*.

For each of the CC-42, CC-22 and CC-61 cattle-associated lineages in BAPS cluster 4 and 10, a different set of specific accessory genes was identified. This may reflect independent colonisation events of that particular host in the evolutionary history of *Campylobacter* [60]. In BAPS cluster 10 we have identified genes associated with a HicA-HicB toxin/antitoxin system, which is suspected to inhibit the bacterial mRNA transfer in case of limited nutrient availability [61–63].

Sharing the same host does not necessarily mean ample opportunities for DNA transfer with the host, since the preferred (sub-)niche of these CCs within the gut of cattle may differ, as it has been assumed for host-generalist lineages previously [49]. Furthermore, structure and composition of the gut microbiome may play a role, however little is known about the microbiome ecology and the putative lineage-specific differences among *C. jejuni* with respect to virulence-associated strategies such as attachment to host cell tissue [64, 65].

We identified a putative cattle-specific allelic variant of DNA polymerase III subunit alpha encoded by *dna*E, in which mutations have been shown to increase the overall mutation rate of *E. coli* [66, 67]. Since an increased mutation rate is well known as a factor influencing niche adaptation [53], the *dna*E variant may promote the host-specialization processes. In addition, we found cattle-specific changes of the gene encoding Ffh, a signal recognition particle protein (SRP). Ffh initiates the co-translational targeting of membrane and secretory proteins to the cytoplasmic bacterial membrane [68], indicating adaptation of transport processes. In *E. coli*, the SRP system plays an important role in membrane protein biosynthesis, and previous research also indicated that Ffh is involved in the regulation of membrane protein translation [69]. Notably, a GTPase (FlhF) possessing an active domain most similar to Ffh, was found to be involved in flagellar gene regulation and biosynthesis in *C. jejuni* [70]. Again, the lack of corresponding recombination patterns indicated that niche-specific environmental pressure induced the predicted ‘aa’ change of Ffh independently in distantly related lineages as we demonstrated in Figure 3. Indeed, *ffh* has already been described as a homoplasic gene on a nucleotide level in cattle-associated *C. jejuni* genomes by a recent study [29].

Most of the CC-403 and ST-1942 (pig-associated) *C. jejuni* in BAPS cluster 11 carry a unique set of genes encoding restriction modification (RM) systems (RM I and RM II) that may contribute to lineage-specific barriers shielding the bacteria from intrusion of foreign DNA, a phenomenon reported before [71–73]. As well, the frequency and pattern of intra-lineage recombination events was unique to CC-403 and its related STs, as noted before [74].

While ‘aa’ variants encoded by the *tenl* gene is thought to affect the thiamine metabolism and may serve as markers for cattle-specific niche adaption [29], in this study we identified pig- specific variations as well. The ‘aa’ changes associated with the allelic variant encoding final aromatase (TenI) needed in thiamine biosynthesis were extensive and may indicate functional alterations or even loss-of-function. Further research to characterise this gene would be useful for potential agrifood intervention strategies. Since industrial diets for pigs are generally supplemented with thiamine [75], reduction or even shutting-off the metabolic pathway might conserve energy and seems therefore beneficial for pig-specialized *C. jejuni* lineages. In addition, we identified a pig-specific variant of the putative thiamine-dependent synthase encoded by *dxs*, again underlining the general importance of specific alterations of the thiamine pathway for host adaptation of *C. jejuni* lineages. The majority of the accessory genome assigned in this study as chicken-specific included, among others, genes for a putative conjugative transfer protein (*Tra*G-like), which is commonly linked to a type IV secretion system essential for DNA transfer in bacterial conjugation [76, 77]. These findings are in concordance with the recombination analysis for the chicken-specific lineages (e.g. CC-257 or CC-354), which indicated multiple horizontal gene transfer events. With respect to *k-mers* that indicate sequence alterations of the core genomes and lead to aa variants of the respective proteins, we noted significant *k-mers* mapping to the gene encoding PycB, the second subunit of the anaplerotic and glucogenic pyruvate carboxylase in *C. jejuni* [78]. This finding indicates a specific adaptation of a basal metabolic pathway in *C. jejuni*. In addition, we detected significant *k-mers* associated with a *rpo*B variant, a housekeeping gene used for investigating genetic relatedness within the *Campylobacter* genus [79]. Interestingly, several different mutations of *rpoB* enhance growth at 42.2°C compared to the wildtype in *E. coli* [80]. Since the body temperature of poultry is commonly between 39 and 43°C [81], the *rpoB* variant might contribute to temperature– induced adaptive changes *in C. jejuni*.

The large host-generalist lineages belonging to either BAPS clusters 2, 3, 6 (CC-21/CC-48/CC-206) or BAPS cluster 8 (CC-45) showed clear differences concerning their accessory gene content, an observation confirmed by earlier results from Yahara et al., who tracked these lineages from the chicken flock through the meat production chain as well as in clinical samples of human origin [22]. Here, we have provided evidence that accessory gene patterns were mostly BAPS clusters-specific, irrespective of the sample origin (e.g. animal, human clinical or environment). Host-generalist BAPS clusters appear to possess a larger pool of accessory genes, possibly indicating a repertoire of genes promoting survival in different hosts and environments [82, 83]. This idea is supported by our recombination analysis, showing that host-generalist lineages are prone to DNA exchange, thus, natural transformation and recombination between host-generalist lineages enhances adaptive possibilities needed to survive in different hosts.

Variation of predicted aa sequences possibly associated with a host–generalist lifestyle of specific *C. jejuni* lineages were, for instance, identified for the cell division protein encoded by *fts*X. Recent work by Riedel et al. showed that *fts*X transcription is downregulated in *Campylobacter lari* after exposure to heat stress [84], possibly indicating certain allelic variants may differ with respect to their stress response. As mentioned earlier, allelic variants may have evolved individually in both lineages (CC-45 and CC-21/CC-48), since the recombination analysis suggests a limited number of recombination events between BAPS clusters 8 and 2, 3 and 6.

Distinct host-specific factors, such as body temperature, the structure and composition of the gut microbiota, mucosal structures and immune system shape the adaptation strategies of *C. jejuni* lineages. Focusing fundamental science research in these areas will enhance the opportunity to exploit this foodborne pathogen’s ability to thrive in niche environments with targeted intervention strategies in the future.

## Acknowledgment

We thank Petra Hahs and Corinna Fruth for their excellent assistance in the laboratory of the National Reference Centre for *Salmonella* and other Bacterial Enterics at the RKI. We also thank the Federal State Laboratories for isolating *Campylobacter* from food matrices and all members of the NRL for *Campylobacter* for technical support.

This research was accomplished within the PAC-CAMPY research network, a part of the national Zoonotic Infectious Diseases Research Network which is funded by the Federal Ministry of Education and Research (BMBF) with grant 01KI1725F, 01KI2007F, and 01KI1725B. Additional funding was received from the BMBF-funded research network #1HealthPREVENT (grant 01KI1727F) and SFB project of BfR 1322–646.

## References

1. Kaakoush NO, Castaño-Rodríguez N, Mitchell HM, Man SM. Global Epidemiology of Campylobacter Infection. Clinical microbiology reviews [Internet]. 2015 Jul 10;28(3):687– 720. Available from: https://cmr.asm.org/content/28/3/687

2. Burnham PM, Hendrixson DR. Campylobacter jejuni: collective components promoting a successful enteric lifestyle. Nature reviews Microbiology [Internet]. 2018 Sep 11;16(9):551–65. Available from: http://www.nature.com/articles/s41579-018-0037-9

3. Humphrey T, O’Brien S, Madsen M. *Campylobacters* as zoonotic pathogens: a food production perspective. International Journal of Food Microbiology [Internet]. 2007 Jul;117(3):237–57. Available from: https://linkinghub.elsevier.com/retrieve/pii/S0168160507000815

4. Hale CR, Scallan E, Cronquist AB, Dunn J, Smith K, Robinson T, et al. Estimates of enteric illness attributable to contact with animals and their environments in the United States. Clinical Infectious Diseases. 2012;

5. Friedman CR, Hoekstra RM, Samuel M, Marcus R, Bender J, Shiferaw B, et al. Risk Factors for Sporadic *Campylobacter* Infection in the United States: A Case-Control Study in FoodNet Sites. Clinical Infectious Diseases. 2004;

6. Marder EP, Cieslak PR, Cronquist AB, Dunn J, Lathrop S, Rabatsky-Ehr T, et al. Incidence and trends of infections with pathogens transmitted commonly through food and the effect of increasing use of culture-independent diagnostic tests on surveillance — Foodborne diseases active surveillance network, 10 U.S. Sites, 2013–2016. Morbidity and Mortality Weekly Report. 2017;

7. Tack DM, Marder EP, Griffin PM, Cieslak PR, Dunn J, Hurd S, et al. Preliminary incidence and trends of infections with pathogens transmitted commonly through food — foodborne diseases active surveillance network, 10 U.S. sites, 2015–2018. Morbidity and Mortality Weekly Report. 2019;

8. Hoffmann S, Maculloch B, Batz M. Economic burden of major foodborne illnesses acquired in the United States. In: Economic Cost of Foodborne Illnesses in the United States. 2015.

9. EFSA Panel on Biological Hazards (BIOHAZ). Scientific Opinion on Campylobacter in broiler meat production: control options and performance objectives and/or targets at different stages of the food chain. EFSA Journal [Internet]. 2011 Apr 1 [cited 2019 Sep 5];9(4):2105. Available from: http://doi.wiley.com/10.2903/j.efsa.2011.2105

10. European Food Safety Authority and European Centre for Disease Prevention and Control. The European Union summary report on trends and sources of zoonoses, zoonotic agents and food-borne outbreaks in 2016. EFSA Journal [Internet]. 2017 Dec;15(12). Available from: http://dx.doi.org/10.2903/j.efsa.2017.5077 LB - KxcE

11. Mangen M-JJ, Plass D, Havelaar AH, Gibbons CL, Cassini A, Mühlberger N, et al. The pathogen- and incidence-based DALY approach: an appropriate [corrected] methodology for estimating the burden of infectious diseases. PLoS One [Internet]. 2013;8(11):e79740. Available from: http://dx.doi.org/10.1371/journal.pone.0079740

12. Epps SVR, Harvey RB, Hume ME, Phillips TD, Anderson RC, Nisbet DJ. Foodborne *Campylobacter*: Infections, metabolism, pathogenesis and reservoirs [Internet]. Vol. 10, International Journal of Environmental Research and Public Health. Multidisciplinary Digital Publishing Institute (MDPI); 2013 [cited 2019 Oct 22]. p. 6292–304. Available from: http://www.ncbi.nlm.nih.gov/pubmed/24287853

13. Louwen R, van Baarlen P, van Vliet AHM, van Belkum A, Hays JP, Endtz HP. *Campylobacter* bacteremia: A rare and under-reported event? European Journal of Microbiology and Immunology. 2012;

14. Rees JH, Soudain SE, Gregson NA, Hughes RAC. *Campylobacter jejuni* Infection and Guillain–Barré Syndrome. New England Journal of Medicine [Internet]. 1995 Nov 23;333(21):1374–9. Available from: http://www.nejm.org/doi/abs/10.1056/NEJM199511233332102

15. Didelot X, Falush D. Inference of Bacterial Microevolution Using Multilocus Sequence Data. Genetics [Internet]. 2007 Mar;175(3):1251–66. Available from: http://www.genetics.org/lookup/doi/10.1534/genetics.106.063305

16. Sheppard SK, Colles FM, McCarthy ND, Strachan NJC, Ogden ID, Forbes KJ, et al. Niche segregation and genetic structure of *Campylobacter jejuni* populations from wild and agricultural host species. Molecular ecology [Internet]. 2011 Aug [cited 2019 Oct 22];20(16):3484–90. Available from: http://www.ncbi.nlm.nih.gov/pubmed/21762392

17. Griekspoor P, Colles FM, McCarthy ND, Hansbro PM, Ashhurst-Smith C, Olsen B, et al. Marked host specificity and lack of phylogeographic population structure of *Campylobacter jejuni* in wild birds. Molecular Ecology [Internet]. 2013 Mar [cited 2019 Oct 22];22(5):1463–72. Available from: http://www.ncbi.nlm.nih.gov/pubmed/23356487

18. Ogden ID, Dallas JF, MacRae M, Rotariu O, Reay KW, Leitch M, et al. *Campylobacter* excreted into the environment by animal sources: Prevalence, concentration shed, and host association. Foodborne Pathogens and Disease [Internet]. 2009 Dec [cited 2019 Oct 22];6(10):1161–70. Available from: http://www.ncbi.nlm.nih.gov/pubmed/19839759

19. Dearlove BL, Cody AJ, Pascoe B, Méric G, Wilson DJ, Sheppard SK. Rapid host switching in generalist *Campylobacter* strains erodes the signal for tracing human infections. The ISME Journal [Internet]. 2016 Mar 25 [cited 2019 Oct 22];10(3):721–9. Available from: http://www.nature.com/articles/ismej2015149

20. Hermans D, Van Deun K, Martel A, Van Immerseel F, Messens W, Heyndrickx M, et al. Colonization factors of *Campylobacter jejuni* in the chicken gut. Veterinary Research [Internet]. 2011;42(1):82. Available from: http://www.veterinaryresearch.org/content/42/1/82

21. Sheppard SK, Didelot X, Meric G, Torralbo A, Jolley KA, Kelly DJ, et al. Genome-wide association study identifies vitamin B5 biosynthesis as a host specificity factor in *Campylobacter*. Proc Natl Acad Sci U S A [Internet]. 2013 Jul 16 [cited 2018 Aug 15];110(29):11923–7. Available from: http://www.ncbi.nlm.nih.gov/pubmed/23818615

22. Yahara K, Méric G, Taylor AJ, de Vries SPW, Murray S, Pascoe B, et al. Genome-wide association of functional traits linked with *Campylobacter jejuni* survival from farm to fork. Environ Microbiol. 2017;19(1):361–80.

23. Thépault A, Méric G, Rivoal K, Pascoe B, Mageiros L, Touzain F, et al. Genome-Wide Identification of Host-Segregating Epidemiological Markers for Source Attribution in Campylobacter jejuni. Elkins CA, editor. Appl Environ Microbiol [Internet]. 2017 Apr 1;83(7):e03085--16. Available from: http://aem.asm.org/lookup/doi/10.1128/AEM.03085-16

24. Buchanan CJ, Webb AL, Mutschall SK, Kruczkiewicz P, Barker DORR, Hetman BM, et al. A Genome-Wide Association Study to Identify Diagnostic Markers for Human Pathogenic *Campylobacter jejuni* Strains. Frontiers in Microbiology [Internet]. 2017 Jun 30 [cited 2018 Aug 15];8:1224. Available from: http://journal.frontiersin.org/article/10.3389/fmicb.2017.01224/full

25. de Vries SPW, Gupta S, Baig A, Wright E, Wedley A, Jensen AN, et al. Genome-wide fitness analyses of the foodborne pathogen *Campylobacter jejuni* in in vitro and in vivo models. Scientific Reports [Internet]. 2017 Dec 28;7(1):1251. Available from: http://www.nature.com/articles/s41598-017-01133-4

26. Habib I, Uyttendaele M, De Zutter L. Survival of poultry-derived *Campylobacter jejuni* of multilocus sequence type clonal complexes 21 and 45 under freeze, chill, oxidative, acid and heat stresses. Food Microbiology. 2010;

27. Alter T, Scherer K. Stress response of *Campylobacter* spp. and its role in food processing. Journal of Veterinary Medicine Series B: Infectious Diseases and Veterinary Public Health. 2006.

28. Murphy C, Carroll C, Jordan KN. Environmental survival mechanisms of the foodborne pathogen *Campylobacter jejuni*. J Appl Microbiol [Internet]. 2006 Apr 1 [cited 2019 Oct 22];100(4):623–32. Available from: http://doi.wiley.com/10.1111/j.1365-2672.2006.02903.x

29. Mourkas E, Taylor AJ, Méric G, Bayliss SC, Pascoe B, Mageiros L, et al. Agricultural intensification and the evolution of host specialism in the enteric pathogen *Campylobacter jejuni*. Proceedings of the National Academy of Sciences [Internet]. 2020 May 4 [cited 2020 May 13]; Available from: https://www.pnas.org/content/early/2020/04/28/1917168117

30. Epping L, Golz JC, Knüver M-T, Huber C, Thürmer A, Wieler LH, et al. Comparison of different technologies for the decipherment of the whole genome sequence of *Campylobacter jejuni* BfR-CA-14430. Gut Pathogens [Internet]. 2019 Dec 16 [cited 2020 Jan 3];11(1):59. Available from: https://gutpathogens.biomedcentral.com/articles/10.1186/s13099-019-0340-7

31. Roehr JT, Dieterich C, Reinert K. Flexbar 3.0 – SIMD and multicore parallelization. Birol I, editor. Bioinformatics [Internet]. 2017 Sep 15;33(18):2941–2. Available from: https://academic.oup.com/bioinformatics/article/33/18/2941/3852078

32. Nikolenko SI, Korobeynikov AI, Alekseyev MA. BayesHammer: Bayesian clustering for error correction in single-cell sequencing. BMC Genomics. 2013;14 Suppl 1:S7.

33. Bankevich A, Nurk S, Antipov D, Gurevich AA, Dvorkin M, Kulikov AS, et al. SPAdes: A new genome assembly algorithm and its applications to single-cell sequencing. Journal of Computational Biology. 2012;19(5):455–77.

34. Seemann T. Prokka: Rapid prokaryotic genome annotation. Bioinformatics [Internet]. 2014 Jul 15;30(14):2068–9. Available from: https://academic.oup.com/bioinformatics/article-lookup/doi/10.1093/bioinformatics/btu153

35. Dingle KE, Colles FM, Wareing DRAA, Ure R, Fox AJ, Bolton FE, et al. Multilocus sequence typing system for *Campylobacter jejuni*. Journal of Clinical Microbiology [Internet]. 2001 Jan 1;39(1):14–23. Available from: http://jcm.asm.org/cgi/doi/10.1128/JCM.39.1.14-23.2001

36. Jolley KA, Maiden MCJ. BIGSdb: Scalable analysis of bacterial genome variation at the population level. BMC Bioinformatics. 2010;

37. Zhou Z, Alikhan NF, Sergeant MJ, Luhmann N, Vaz C, Francisco AP, et al. Grapetree: Visualization of core genomic relationships among 100,000 bacterial pathogens. Genome Research. 2018;

38. Page AJ, Cummins CA, Hunt M, Wong VK, Reuter S, Holden MTG, et al. Roary: rapid large-scale prokaryote pan genome analysis. Bioinformatics [Internet]. 2015 Nov 15;31(22):3691–3. Available from: https://academic.oup.com/bioinformatics/article-lookup/doi/10.1093/bioinformatics/btv421

39. Stamatakis A. RAxML version 8: a tool for phylogenetic analysis and post-analysis of large phylogenies. Bioinformatics [Internet]. 2014 May 1 [cited 2019 Jul 8];30(9):1312–3. Available from: https://academic.oup.com/bioinformatics/article-lookup/doi/10.1093/bioinformatics/btu033

40. Tavaré S. Some probabilistic and statistical problems in the analysis of DNA sequences [Internet]. Vol. 17, American Mathematical Society: Lectures on Mathematics in the Life Sciences. Providence, R.I. American Mathematical Society, c1986.; 1986 [cited 2019 Oct 22]. p. 57–86. Available from: http://agris.fao.org/agris-search/search.do?recordID=US201301755037

41. Didelot X, Wilson DJ. ClonalFrameML: Efficient Inference of Recombination in Whole Bacterial Genomes. Prlic A, editor. PLOS Computational Biology [Internet]. 2015 Feb 12;11(2):e1004041. Available from: https://dx.plos.org/10.1371/journal.pcbi.1004041

42. Tonkin-Hill G, Lees JA, Bentley SD, Frost SDWW, Corander J. RhierBAPs: An R implementation of the population clustering algorithm hierbaps [version 1; referees: 2 approved]. Wellcome Open Research. 2018;3:93.

43. Maaten L van der, Hinton G. Visualizing data using t-SNE. Journal of machine learning research [Internet]. 2008;9(Nov):2579–605. Available from: http://www.jmlr.org/papers/v9/vandermaaten08a.html

44. Marttinen P, Hanage WP, Croucher NJ, Connor TR, Harris SR, Bentley SD, et al. Detection of recombination events in bacterial genomes from large population samples. Nucleic Acids Research [Internet]. 2012 Jan [cited 2019 Jul 11];40(1):e6. Available from: http://www.ncbi.nlm.nih.gov/pubmed/22064866

45. Lees JA, Galardini M, Bentley SD, Weiser JN, Corander J. pyseer: A comprehensive tool for microbial pangenome-wide association studies. Bioinformatics. 2018;34(24):4310–2.

46. Li H, Durbin R. Fast and accurate short read alignment with Burrows-Wheeler transform. Bioinformatics. 2009;25(14):1754–60.

47. Huerta-Cepas J, Szklarczyk D, Forslund K, Cook H, Heller D, Walter MC, et al. eggNOG 4.5: a hierarchical orthology framework with improved functional annotations for eukaryotic, prokaryotic and viral sequences. Nucleic Acids Research. 2016;44(D1):D286--93.

48. Huerta-Cepas J, Forslund K, Coelho LP, Szklarczyk D, Jensen LJ, von Mering C, et al. Fast genome-wide functional annotation through orthology assignment by eggNOG-mapper. Molecular Biology and Evolution [Internet]. 2017 Aug 1;34(8):2115–22. Available from: https://academic.oup.com/mbe/article/34/8/2115/3782716

49. Sheppard SK, Cheng L, Méric G, De Haan CPA, Llarena AK, Marttinen P, et al. Cryptic ecology among host generalist *Campylobacter jejuni* in domestic animals. Molecular Ecology [Internet]. 2014 May [cited 2019 Oct 22];23(10):2442–51. Available from: http://www.ncbi.nlm.nih.gov/pubmed/24689900

50. Schröder G, Lanka E. TraG-like proteins of type IV secretion systems: Functional dissection of the multiple activities of TraG (RP4) and TrwB (R388). Journal of Bacteriology. 2003;

51. Poly F, Threadgill D, Stintzi A. Genomic diversity in *Campylobacter jejuni*: Identification of *C. jejuni* 81-176-specific genes. Journal of Clinical Microbiology. 2005 May;43(5):2330–8.

52. Lee KY, Lee KY, Kim JH, Lee IG, Lee SH, Sim DW, et al. Structure-based functional identification of *Helicobacter pylori* HP0268 as a nuclease with both DNA nicking and RNase activities. Nucleic Acids Research. 2015;

53. Sheppard SK, Guttman DS, Fitzgerald JR. Population genomics of bacterial host adaptation. Nature Reviews Genetics [Internet]. 2018 Sep 4 [cited 2018 Aug 14];19(9):549–65. Available from: http://www.nature.com/articles/s41576-018-0032-z

54. Mohan V, Stevenson M, Marshall J, Fearnhead P, Holland BR, Hotter G, et al. *Campylobacter jejuni* colonization and population structure in urban populations of ducks and starlings in New Zealand. MicrobiologyOpen [Internet]. 2013 Aug;2(4):659–73. Available from: http://doi.wiley.com/10.1002/mbo3.102

55. Hershberg R. Mutation—the engine of evolution: Studying mutation and its role in the evolution of bacteria. Cold Spring Harbor Perspectives in Biology. 2015;

56. Falush D. Bacterial genomics: Microbial GWAS coming of age. Nature microbiology [Internet]. 2016 May 26;1(5):16059. Available from: http://www.nature.com/articles/nmicrobiol201659

57. Power RA, Parkhill J, de Oliveira T. Microbial genome-wide association studies: lessons from human GWAS. Nature Reviews Genetics [Internet]. 2017 Jan 14;18(1):41–50. Available from: http://www.nature.com/articles/nrg.2016.132

58. Brandley MC, Warren DL, Leaché AD, McGuire JA. Homoplasy and clade support. Systematic Biology. 2009;

59. Hassanin A, Lecointre G, Tillier S. The ‘evolutionary signal’ of homoplasy in proteincoding gene sequences and its consequences for a priori weighting in phylogeny. Comptes Rendus de l’Académie des Sciences - Series III - Sciences de la Vie. 1998;

60. Sheppard SK, Maiden MCJ. The Evolution of *Campylobacter jejuni* and *Campylobacter coli*. Cold Spring Harbor Perspectives in Biology. 2015 Aug;7(8):a018119.

61. Motiejunaite R, Armalyte J, Markuckas A, Sužiedeliene E. Escherichia coli dinJ-yafQ genes act as a toxin-antitoxin module. FEMS Microbiology Letters. 2007;

62. Buts L, Lah J, Dao-Thi MH, Wyns L, Loris R. Toxin-antitoxin modules as bacterial metabolic stress managers. Trends in Biochemical Sciences. 2005.

63. Gerdes K, Christensen SK, Løbner-Olesen A. Prokaryotic toxin-antitoxin stress response loci. Nature Reviews Microbiology. 2005.

64. Han Z, Willer T, Li L, Pielsticker C, Rychlik I, Velge P, et al. Influence of the gut microbiota composition on *Campylobacter jejuni* colonization in chicken. Infection and Immunity. 2017;

65. Indikova I, Humphrey TJ, Hilbert F. Survival with a helping hand: *Campylobacter* and microbiota. Frontiers in Microbiology. 2015;

66. Fijalkowska IJ, Schaaper RM, Jonczyk P. DNA replication fidelity in *Escherichia coli*: A multi-DNA polymerase affair. FEMS Microbiology Reviews. 2012.

67. Vandewiele D, Fernández de Henestrosa AR, Timms AR, Bridges BA, Woodgate R. Sequence analysis and phenotypes of five temperature sensitive mutator alleles of dnaE, encoding modified α-catalytic subunits of Escherichia coli DNA polymerase III holoenzyme. Mutation Research - Fundamental and Molecular Mechanisms of Mutagenesis. 2002;

68. Shan SO, Stroud RM, Walter P. Mechanism of association and reciprocal activation of two GTPases. PLoS Biology. 2004;

69. Yosef I, Bochkareva ES, Bibi E. Escherichia coli SRP, its protein subunit Ffh, and the Ffh M domain are able to selectively limit membrane protein expression when overexpressed. mBio. 2010;

70. Balaban M, Joslin SN, Hendrixson DR. FlhF and its GTPase activity are required for distinct processes in flagellar gene regulation and biosynthesis in Campylobacter jejuni. Journal of Bacteriology. 2009;

71. Budroni S, Siena E, Dunning Hotopp JC, Seib KL, Serruto D, Nofroni C, et al. *Neisseria meningitidis* is structured in clades associated with restriction modification systems that modulate homologous recombination. Proceedings of the National Academy of Sciences of the United States of America. 2011;

72. McCarthy ND, Colles FM, Dingle KE, Bagnall MC, Manning G, Maiden MCJ, et al. Host-associated genetic import in *Campylobacter jejuni*. Emerging Infectious Diseases. 2007;13(2):267–72.

73. Asakura H, Brüggemann H, Sheppard SK, Ekawa T, Meyer TF, Yamamoto S, et al. Molecular Evidence for the Thriving of Campylobacter jejuni ST-4526 in Japan. Bereswill S, editor. PLoS ONE [Internet]. 2012 Nov 7;7(11):e48394. Available from: https://dx.plos.org/10.1371/journal.pone.0048394

74. Morley L, McNally A, Paszkiewicz K, Corander J, Méric G, Sheppard SK, et al. Gene Loss and Lineage-Specific Restriction-Modification Systems Associated with Niche Differentiation in the Campylobacter jejuni Sequence Type 403 Clonal Complex. Schloss PD, editor. Applied and environmental microbiology [Internet]. 2015 Jun 1 [cited 2019 Jul 11];81(11):3641–7. Available from: http://www.ncbi.nlm.nih.gov/pubmed/25795671

75. National Research Council. Nutrient Requirements of Swine [Internet]. National Research Council, editor. Nutrient Requirements of Swine. Washington, D.C.: National Academies Press; 2012. Available from: http://www.nap.edu/catalog/13298

76. Schröder G, Krause S, Zechner EL, Traxler B, Yeo HJ, Lurz R, et al. TraG-like proteins of DNA transfer systems and of the *Helicobacter pylori* type IV secretion system: Inner membrane gate for exported substrates? Journal of Bacteriology. 2002;

77. Kienesberger S, Trummler CS, Fauster A, Lang S, Sprenger H, Gorkiewicz G, et al. Interbacterial macromolecular transfer by the *Campylobacter fetus* subsp. *venerealis* type IV secretion system. Journal of Bacteriology. 2011;

78. Velayudhan J, Kelly DJ. Analysis of gluconeogenic and anaplerotic enzymes in *Campylobacter jejuni*: An essential role for phosphoenolpyruvate carboxykinase. Microbiology [Internet]. 2002 Mar 1 [cited 2019 Jul 16];148(3):685–94. Available from: http://mic.microbiologyresearch.org/content/journal/micro/10.1099/00221287-148-3-685

79. Korczak BM, Stieber R, Emler S, Burnens AP, Frey J, Kuhnert P. Genetic relatedness within the genus *Campylobacter* inferred from rpoB sequences. International Journal of Systematic and Evolutionary Microbiology. 2006;

80. González-González A, Hug SM, Rodríguez-Verdugo A, Patel JS, Gaut BS. Adaptive mutations in RNA polymerase and the transcriptional terminator rho have similar effects on *Escherichia coli* gene expression. Molecular Biology and Evolution [Internet]. 2017 Nov 1 [cited 2019 Sep 3];34(11):2839–55. Available from: http://www.ncbi.nlm.nih.gov/pubmed/28961910

81. Richards SA. The significance of changes in the temperature of the skin and body core of the chicken in the regulation of heat loss. The Journal of Physiology. 1971;

82. Hottes AK, Freddolino PL, Khare A, Donnell ZN, Liu JC, Tavazoie S. Bacterial Adaptation through Loss of Function. PLoS Genetics. 2013;

83. Iranzo J, Wolf YI, Koonin E V., Sela I. Gene gain and loss push prokaryotes beyond the homologous recombination barrier and accelerate genome sequence divergence. Nature Communications. 2019;

84. Riedel C, Förstner KU, Püning C, Alter T, Sharma CM, Gölz G. Differences in the Transcriptomic Response of *Campylobacter coli* and *Campylobacter lari* to Heat Stress. Frontiers in Microbiology. 2020;

