## Supplementary Material for "Genome-wide insights into population structure and host specificity of *Campylobacter jejuni*"

***Supplementary Material:* Population Structure and Host Specificity of *Campylobacter jejuni* in Germany - a Genomic Approach**

### Genome sequencing

DNA for whole genome sequencing (WGS) of the German isolates was prepared using the PureLink Genomic DNA Mini Kit (Thermo Fisher Scientific Inc., Waltham, MA, USA) or the DNeasy Blood & Tissue Kit (QIAGEN, Hilden, Germany). WGS sequencing libraries were generated with the Nextera XT (Illumina Inc., San Diego, CA, USA) library kit following the manufactures instructions. Sequencing was performed on a MiSeq sequencer (MiSeq Reagent Kit v.3; Illumina Inc., San Diego, CA, USA) resulting in 300-bp paired-end reads and an average coverage of 80x, and on a HiSeq 1500 using a PE Rapid Cluster Kit v2 and Rapid SBS Kit v2 (500 cycles; Illumina Inc., San Diego, CA) resulting in 250-bp paired-end reads and an average coverage of 80x.

**MLST Trees and Host Distribution**

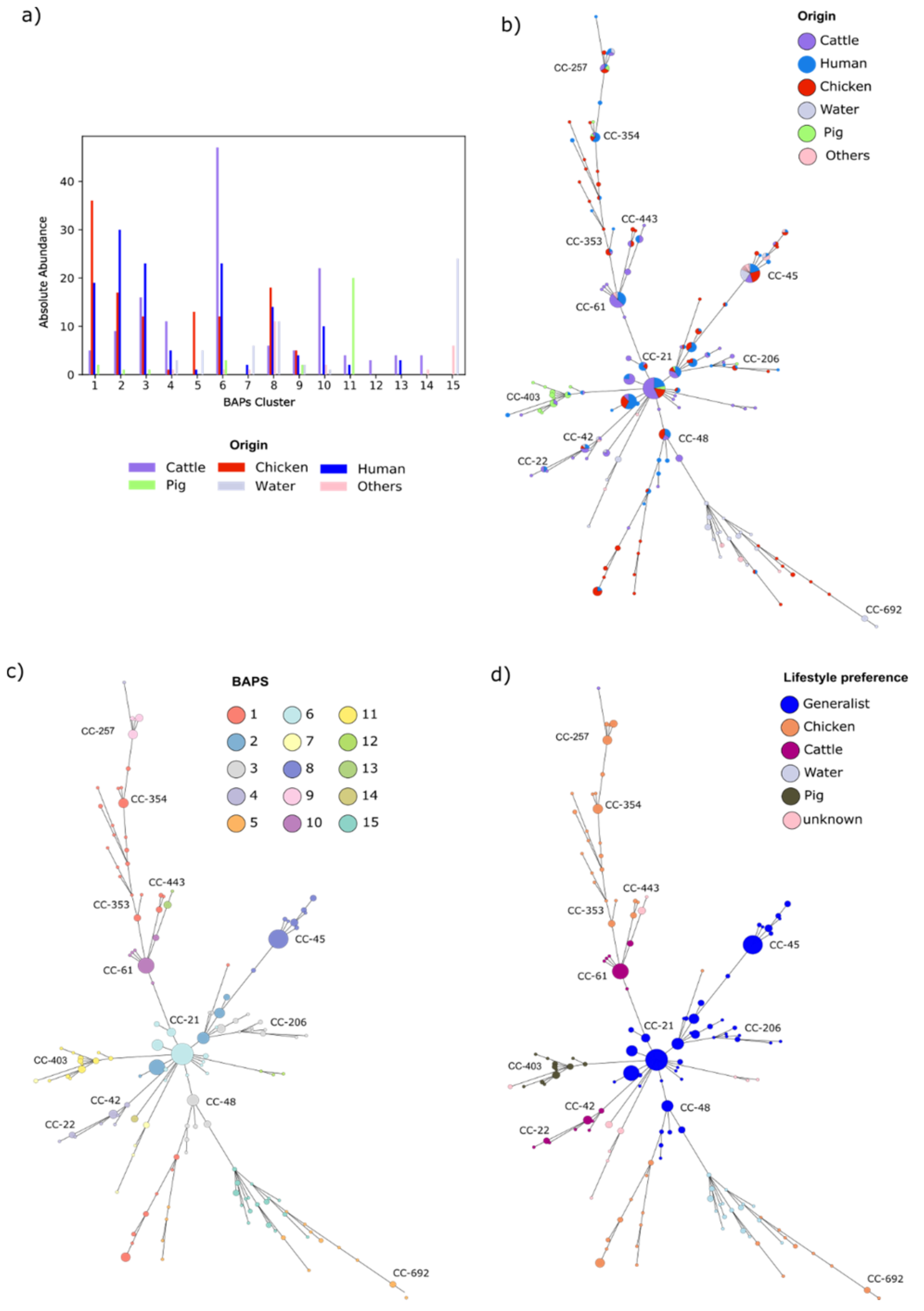


**Figure S1:** Subplot a) shows absolute distributions of host associations among the BAPs cluster that are later used for the stratified random sampling approach. Minimum spanning trees shows relationship between different MLSTs based on the 7 housekeeping genes colored by b) sampling origin, c) BAPS cluster and d) host association cluster. Subplot d) and e) show relative and absolute distributions of host associations among the BAPs cluster that are later used for the stratified random sampling approach.

**Genome-wide association study (GWAS)**

In order to perform an in-depth analysis of genomic alterations possibly associated with host specificity, pyseer v.1.1.2 (1) was used for GWAS based on variable-length *k-mer* composition (9 to 100 base pairs) for all 490 genomes, counted by fsm-lite v1.0 (https://github.com/nvalimak/fsm-lite). As a next step, *k-mer* counts of genomes representing distinct isolate origins (human, cattle, chicken or pig) were investigated with respect to significant differences: counts obtained for each group of isolate origins were compared against the combined counts obtained for genomes representing isolates of all other groups. For each comparison, significant *k-mers* were filtered by an individually calculated threshold (based on the Bonferroni correction) for the lineage corrected p-value obtained from pyseer and split into two groups based on their direction of effect. The significant *k-mers* were then mapped by bwa v0.7.17 (2) against selected reference genomes from this study set in order to identify putative origin-specific factors, genes and consecutive gene loci.

In order to reduce the false positive rate and account for highly unbalanced groups, we employed bootstrapping with a proportional stratified random sampling approach, which was weighted by size of the BAPS clusters. This approach was applied on the control group in order to compare equally sized groups. The whole approach was repeated 100 times for genomes representing each isolate origin to set up a consensus approach. Genes identified in at least 90% of these tests were selected as candidate genes. Of note, putative genes important for host-generalist lineages were not affected by the approach, since both, the host-generalist- and the host-specialist group contained an equal number of genomes. In this process genes associated with a max -log(p-value) ≥ 80 were selected. The expected frequency of significant *k-mers* is associated with the proportion of isolates assigned to a particular lifestyle (e.g. chicken, cattle, host-generalist) in relation to the total number of isolates.

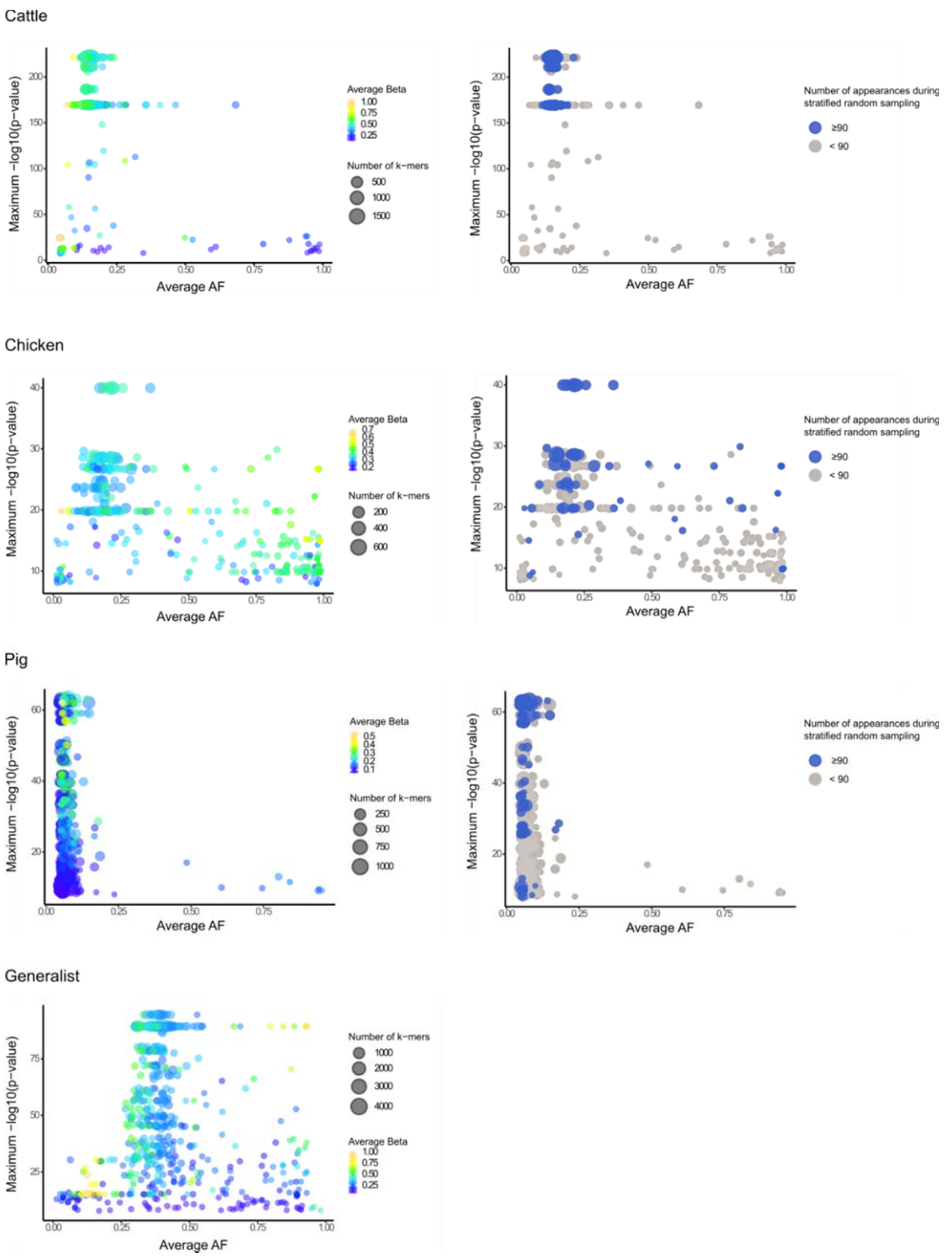


**Figure S2:** Visualizations of genes that were identified by mapping the significant *k-mers* for cattle, chicken, generalist and pig associated genomes. Plots in the left column show estimated GWAS results without the stratified sampling approach. The x-axis shows the average allele frequencies and y-axis the average p-value. The size accounts for the number of *k-mers* mapped to a specific gene whereas the color pattern estimates the average beta value (left column). Plots in the right column show the influence of the stratified random sampling GWAS for cattle, chicken and pig associated genomes. Blue dots where obtained in 90% of the times in the consensus GWAS (right column).

**Supp.Table 2.** BAPS clusters, original sample source and lifestyle preferences of 490 *C. jejuni* genomes

| | | | Sample origin | | | | | | Lifestyle classification | | |
|---|---|---|---|---|---|---|---|---|---|---|---|
| | | | chicken | human | cattle | pig | environment | other | | | |
| **BAPS** | **n** | **CC** | **n** | **n** | **n** | **n** | **n** | **n** | **this study** | **previous studies** | **reference** |
| **1** | **62** | CC-353 | 36 | 19 | 5 | 2 | 0 | 0 | chicken | chicken | (3) |
| | | CC-354 | | | | | | | chicken | chicken | (3) |
| | | CC-443 | | | | | | | chicken | chicken | (3) |
| | | CC-464 | | | | | | | chicken | chicken | (3,4) |
| | | CC-52 | | | | | | | chicken | chicken | (4) |
| | | ST-2274 | | | | | | | **chicken** | - | - |
| **2** | **57** | CC-21 | 17 | 30 | 9 | 1 | 0 | 0 | generalist | chicken | (3,5) |
| **3** | **52** | CC-48 | 12 | 23 | 16 | 1 | 0 | 0 | generalist | generalist | (3) |
| | | CC-206 | | | | | | | generalist | generalist | (3) |
| | | CC-21 | | | | | | | generalist | generalist | (3,5) |
| **4** | **21** | CC-42 | 1 | 5 | 11 | 0 | 3 | 1 | cattle | cattle | (3) |
| | | CC-22 | | | | | | | **cattle** | - | - |
| **5** | **19** | CC-1034 | 13 | 1 | 0 | 0 | 5 | 0 | chicken | wild birds, turkey | (6) |
| | | CC-692 | | | | | | | chicken | wild birds, turkey | (7) |
| **6** | **86** | CC-21 | 12 | 23 | 47 | 3 | 0 | 1 | generalist | generalist | (3) |
| **7** | **9** | CC-677 | 0 | 2 | 0 | 0 | 6 | 1 | n.a. | - | - |
| | | CC-682 | | | | | | | n.a. | - | - |
| **8** | **60** | CC-45 | 18 | 14 | 6 | 0 | 12 | 10 | generalist | generalist | (5) |
| | | CC-283 | | | | | | | generalist | generalist | (3) |
| **9** | **18** | CC-257 | 5 | 4 | 5 | 2 | 2 | 0 | chicken | chicken | (8) |
| **10** | **35** | CC-61 | 0 | 10 | 22 | 0 | 1 | 2 | cattle | cattle | (8) |

| | | | | | | | | | | | |
|---|---|---|---|---|---|---|---|---|---|---|---|
| **11** | **26** | CC-403 | 0 | 2 | 4 | 20 | 0 | 0 | pig | pig | (9) |
| **12** | **3** | ST-3098 | 0 | 0 | 3 | 0 | 0 | 0 | n.a. | - | - |
| | | ST-10282 | | | | | | | n.a. | - | - |
| **13** | **7** | ST-922 | 0 | 3 | 4 | 0 | 0 | 0 | n.a. | - | - |
| **14** | **5** | CC-508 | 0 | 0 | 4 | 0 | 1 | 0 | n.a. | | - |
| **15** | **30** | ST-693 | 0 | 0 | 0 | 0 | 24 | 6 | n.a. | - | - |
| | | ST-996 | | | | | | | n.a. | - | - |
| | | ST-1030 | | | | | | | n.a. | - | - |
| | | ST-1206 | | | | | | | n.a. | - | - |

***n.a. Clusters were not analyzed regarding their lifestyle preference due to the limited amount of isolates**

**References**


1. Lees JA, Galardini M, Bentley SD, Weiser JN, Corander J. pyseer: A comprehensive tool for microbial pangenome-wide association studies. Bioinformatics. 2018;34(24):4310–2.

2. Li H, Durbin R. Fast and accurate short read alignment with Burrows-Wheeler transform. Bioinformatics. 2009;25(14):1754–60.

3. Sheppard SK, Cheng L, Méric G, De Haan CPA, Llarena AK, Marttinen P, et al. Cryptic ecology among host generalist *Campylobacter jejuni* in domestic animals. Molecular Ecology [Internet]. 2014 May [cited 2019 Oct 22];23(10):2442–51. Available from: http://www.ncbi.nlm.nih.gov/pubmed/24689900

4. Noormohamed A, Fakhr M. Molecular Typing of *Campylobacter jejuni* and *Campylobacter coli* Isolated from Various Retail Meats by MLST and PFGE. Foods. 2014;

5. Gripp E, Hlahla D, Didelot X, Kops F, Maurischat S, Tedin K, et al. Closely related *Campylobacter jejuni* strains from different sources reveal a generalist rather than a specialist lifestyle. BMC Genomics. 2011;

6. Mohan V, Stevenson M, Marshall J, Fearnhead P, Holland BR, Hotter G, et al. *Campylobacter jejuni* colonization and population structure in urban populations of ducks and starlings in New Zealand. MicrobiologyOpen [Internet]. 2013 Aug;2(4):659–73. Available from: http://doi.wiley.com/10.1002/mbo3.102

7. Waldenström J, Broman T, Carlsson I, Hasselquist D, Achterberg RP, Wagenaar JA, et al. Prevalence of *Campylobacter jejuni*, *Campylobacter lari*, and *Campylobacter coli* in different ecological guilds and taxa of migrating birds. Applied and Environmental Microbiology. 2002;

8. Sheppard SK, Colles FM, McCarthy ND, Strachan NJC, Ogden ID, Forbes KJ, et al. Niche segregation and genetic structure of *Campylobacter jejuni* populations from wild and agricultural host species. Molecular ecology [Internet]. 2011 Aug [cited 2019 Oct 22];20(16):3484–90. Available from: http://www.ncbi.nlm.nih.gov/pubmed/21762392

9. Morley L, McNally A, Paszkiewicz K, Corander J, Méric G, Sheppard SK, et al. Gene Loss and Lineage-Specific Restriction-Modification Systems Associated with Niche Differentiation in the *Campylobacter jejuni* Sequence Type 403 Clonal Complex. Schloss PD, editor. Applied and environmental microbiology [Internet]. 2015 Jun 1 [cited 2019 Jul 11];81(11):3641–7. Available from: http://www.ncbi.nlm.nih.gov/pubmed/25795671
