## Supplementary material for "Genome-wide insights into population structure and host specificity of *Campylobacter jejuni*": Table S1

Table S1: Overview of sequenced genomes used in this study with additonal meta data information.

| SRR | ID | ST | CC | Origin | Association | BAPS1 | Source |
|---|---|---|---|---|---|---|---|
| SRR12302213 | 12-02934 | 354 | ST-354 | Human | Chicken | 1 | Germany: Federal State N |
| SRR12302212 | 12-02938 | 50 | ST-21 | Human | Generalist | 2 | Germany: Federal State H |
| SRR12302046 | 12-02939 | 10269 | ST-353 | Human | Chicken | 1 | Germany: Federal State H |
| SRR12302257 | 12-03052 | 48 | ST-48 | Human | Generalist | 3 | Germany: Federal State N |
| SRR12302223 | 12-03072 | 42 | ST-42 | Human | Cattle | 4 | Germany: Federal State J |
| SRR12302164 | 12-03499 | 2314 | ST-1034 | Human | Chicken | 5 | Germany: Federal State H |
| SRR12302153 | 12-03501 | 50 | ST-21 | Human | Generalist | 2 | Germany: Federal State G |
| SRR12302142 | 12-03536 | 429 | ST-48 | Human | Generalist | 3 | Germany: Federal State G |
| SRR12302107 | 12-03862 | 21 | ST-21 | Human | Generalist | 6 | Germany: Federal State H |
| SRR12302096 | 12-03906 | 19 | ST-21 | Human | Generalist | 2 | Germany: Federal State H |
| SRR12302211 | 12-03907 | 354 | ST-354 | Human | Chicken | 1 | Germany: Federal State H |
| SRR12302200 | 12-03908 | 19 | ST-21 | Human | Generalist | 2 | Germany: Federal State J |
| SRR12302189 | 12-03909 | 10270 | ST-41 | Human | unknown | 4 | Germany: Federal State J |
| SRR12302178 | 12-03950 | 464 | ST-464 | Human | unknown | 1 | Germany: Federal State N |
| SRR12302167 | 12-03951 | 677 | ST-677 | Human | unknown | 7 | Germany: Federal State H |
| SRR12302128 | 12-03952 | 21 | ST-21 | Human | Generalist | 6 | Germany: Federal State H |
| SRR12302117 | 12-03953 | 267 | ST-283 | Human | unknown | 8 | Germany: Federal State H |
| SRR12302079 | 12-03954 | 50 | ST-21 | Human | Generalist | 2 | Germany: Federal State H |
| SRR12302068 | 12-03967 | 677 | ST-677 | Human | unknown | 7 | Germany: Federal State P |
| SRR12302057 | 13-00141 | 53 | ST-21 | Human | Generalist | 6 | Germany: Federal State N |
| SRR12302045 | 13-00142 | 354 | ST-354 | Human | Chicken | 1 | Germany: Federal State P |
| SRR12302033 | 13-00156 | 21 | ST-21 | Human | Generalist | 6 | Germany: Federal State J |
| SRR12302022 | 13-00216 | 918 | ST-48 | Human | Generalist | 3 | Germany |
| SRR12302011 | 13-00388 | 1326 | ST-45 | Human | Generalist | 8 | Germany: Federal State H |
| SRR12302000 | 13-00578 | 572 | ST-206 | Human | Generalist | 3 | Germany: Federal State H |
| SRR12301989 | 13-00579 | 53 | ST-21 | Human | Generalist | 6 | Germany: Federal State J |
| SRR12301978 | 13-00893 | 21 | ST-21 | Human | Generalist | 6 | Germany: Federal State H |
| SRR12301967 | 13-01164 | 50 | ST-21 | Human | Generalist | 2 | Germany: Federal State P |
| SRR12302279 | 13-01279 | 21 | ST-21 | Human | Generalist | 6 | Germany: Federal State H |
| SRR12302268 | 13-01855 | 1519 | ST-21 | Human | Generalist | 2 | Germany: Federal State N |
| SRR12302256 | 13-01999 | 206 | ST-206 | Human | Generalist | 3 | Germany: Federal State H |
| SRR12302245 | 13-02263 | 50 | ST-21 | Human | Generalist | 2 | Germany |
| SRR12302234 | 13-02264 | 50 | ST-21 | Human | Generalist | 2 | Germany |

| | | | | | | | |
|---|---|---|---|---|---|---|---|
| SRR12302230 | 13-02292 | 354 | ST-354 | Human | Chicken | 1 | Germany: Federal State G |
| SRR12302229 | 13-02407 | 19 | ST-21 | Human | Generalist | 2 | Germany: Federal State H |
| SRR12302228 | 13-02841 | 45 | ST-45 | Human | Generalist | 8 | Germany |
| SRR12302227 | 13-02842 | 45 | ST-45 | Human | Generalist | 8 | Germany |
| SRR12302226 | 14-01211 | 46 | ST-206 | Human | Generalist | 3 | Germany: Federal State H |
| SRR12302225 | 14-01213 | 21 | ST-21 | Human | Generalist | 6 | Germany: Federal State H |
| SRR12302224 | 14-01252 | 990 | ST-257 | Human | Chicken | 9 | Germany: Federal State H |
| SRR12302222 | 14-01255 | 824 | ST-257 | Human | Chicken | 1 | Germany: Federal State H |
| SRR12302221 | 14-01256 | 46 | ST-206 | Human | Generalist | 3 | Germany: Federal State H |
| SRR12302220 | 14-01401 | 7231 | unknown | Human | unknown | 3 | Germany: Federal State H |
| SRR12302219 | 14-01734 | 46 | ST-206 | Human | Generalist | 3 | Germany: Federal State B |
| SRR12302218 | 14-01866 | 50 | ST-21 | Human | Generalist | 2 | Germany: Federal State H |
| SRR12302217 | 14-02206 | 2036 | ST-353 | Human | Chicken | 1 | Germany: Federal State P |
| SRR12302216 | 14-02234 | 1519 | ST-21 | Human | Generalist | 2 | Germany: Federal State J |
| SRR12302215 | 14-02237 | 50 | ST-21 | Human | Generalist | 2 | Germany: Federal State H |
| SRR12302214 | 14-02514 | 354 | ST-354 | Human | Chicken | 1 | Germany: Federal State H |
| SRR12302165 | 14-02648 | 50 | ST-21 | Human | Generalist | 2 | Germany: Federal State P |
| SRR12302163 | 14-02649 | 61 | ST-61 | Human | Cattle | 10 | Germany: Federal State H |
| SRR12302162 | 14-02926 | 19 | ST-21 | Human | Generalist | 2 | Germany: Federal State K |
| SRR12302161 | 14-02989 | 44 | ST-21 | Human | Generalist | 2 | Germany: Federal State K |
| SRR12302160 | 14-02992 | 45 | ST-45 | Human | Generalist | 8 | Germany: Federal State P |
| SRR12302159 | 14-03375 | 262 | ST-21 | Human | Generalist | 6 | Germany: Federal State P |
| SRR12302158 | 15-00051 | 50 | ST-21 | Human | Generalist | 2 | Germany |
| SRR12302157 | 15-00397 | 1044 | ST-658 | Human | unknown | 1 | Germany: Federal State J |
| SRR12302156 | 15-00432 | 10271 | ST-464 | Human | unknown | 1 | Germany: Federal State H |
| SRR12302155 | 15-00440 | 475 | ST-48 | Human | Generalist | 3 | Germany: Federal State H |
| SRR12302154 | 15-00441 | 475 | ST-48 | Human | Generalist | 3 | Germany: Federal State A |
| SRR12302152 | 15-01126 | 354 | ST-354 | Human | Chicken | 1 | Germany: Federal State N |
| SRR12302151 | 15-01144 | 1519 | ST-21 | Human | Generalist | 2 | Germany: Federal State H |
| SRR12302150 | 15-01231 | 824 | ST-257 | Human | Chicken | 1 | Germany: Federal State H |
| SRR12302149 | 15-01311 | 2036 | ST-353 | Human | Chicken | 1 | Germany: Federal State N |
| SRR12302148 | 15-01320 | 2274 | unknown | Human | unknown | 1 | Germany: Federal State H |
| SRR12302147 | 15-01321 | 1519 | ST-21 | Human | Generalist | 2 | Germany: Federal State H |
| SRR12302146 | 15-01399 | 50 | ST-21 | Human | Generalist | 2 | Germany: Federal State G |
| SRR12302145 | 15-01400 | 1003 | ST-45 | Human | Generalist | 8 | Germany: Federal State H |

| | | | | | | | |
|---|---|---|---|---|---|---|---|
| SRR12302144 | 15-01539 | 61 | ST-61 | Human | Cattle | 10 | Germany |
| SRR12302143 | 15-01638 | 61 | ST-61 | Human | Cattle | 10 | Germany |
| SRR12302141 | 15-01641 | 61 | ST-61 | Human | Cattle | 10 | Germany: Federal State J |
| SRR12302140 | 15-01705 | 257 | ST-257 | Human | Chicken | 9 | Germany |
| SRR12302139 | 15-01779 | 137 | ST-45 | Human | Generalist | 8 | Germany |
| SRR12302138 | 15-01780 | 1519 | ST-21 | Human | Generalist | 2 | Germany |
| SRR12302113 | 15-00398 | 50 | ST-21 | Human | Generalist | 2 | Germany: Federal State P |
| SRR12302112 | 16-00049 | 46 | ST-206 | Human | Generalist | 3 | Germany: Federal State H |
| SRR12302111 | 16-00112 | 122 | ST-206 | Human | Generalist | 3 | Germany: Federal State H |
| SRR12302110 | 16-00241 | 48 | ST-48 | Human | Generalist | 3 | Germany: Federal State H |
| SRR12302109 | 16-00242 | 6461 | ST-353 | Human | Chicken | 1 | Germany: Federal State N |
| SRR12302108 | 16-00263 | 658 | ST-658 | Human | unknown | 1 | Germany: Federal State H |
| SRR12302106 | 16-00264 | 1519 | ST-21 | Human | Generalist | 2 | Germany: Federal State H |
| SRR12302105 | 16-00268 | 48 | ST-48 | Human | Generalist | 3 | Germany: Federal State J |
| SRR12302104 | 16-00290 | 6175 | ST-21 | Human | Generalist | 3 | Germany: Federal State N |
| SRR12302103 | 16-00409 | 2274 | unknown | Human | unknown | 1 | Germany: Federal State P |
| SRR12302102 | 16-00431 | 22 | ST-22 | Human | unknown | 4 | Germany: Federal State G |
| SRR12302101 | 16-00576 | 1519 | ST-21 | Human | Generalist | 2 | Germany: Federal State H |
| SRR12302100 | 16-00632 | 50 | ST-21 | Human | Generalist | 2 | Germany: Federal State N |
| SRR12302099 | 16-00663 | 21 | ST-21 | Human | Generalist | 6 | Germany: Federal State H |
| SRR12302098 | 16-01339 | 1775 | ST-403 | Human | Pig | 11 | Germany: Federal State H |
| SRR12302097 | 16-01340 | 21 | ST-21 | Human | Generalist | 6 | Germany: Federal State H |
| SRR12302095 | 16-01341 | 50 | ST-21 | Human | Generalist | 2 | Germany: Federal State J |
| SRR12302094 | 19-00788 | 658 | ST-177 | Human | unknown | 1 | Germany |
| SRR12302093 | 19-00791 | 21 | ST-21 | Human | Generalist | 6 | Germany |
| SRR12302092 | 19-00792 | 22 | ST-22 | Human | unknown | 4 | Germany |
| SRR12302091 | 19-00811 | 61 | ST-61 | Human | Cattle | 10 | Germany |
| SRR12302090 | 19-00812 | 61 | ST-61 | Human | Cattle | 10 | Germany |
| SRR12302089 | BfRCA02030 | 435 | ST-403 | Pig | Pig | 11 | Germany |
| SRR12302088 | BfRCA05380 | 257 | ST-257 | Pig | Chicken | 9 | Germany |
| SRR12302087 | BfRCA05532NCT | 43 | ST-21 | Chicken | Generalist | 6 | Germany |
| SRR12302038 | BfRCA05915 | 257 | ST-257 | Pig | Chicken | 9 | Germany |
| SRR12302210 | BfRCA05925 | 9340 | ST-403 | Pig | Pig | 11 | Germany |
| SRR12302209 | BfRCA06044 | 21 | ST-21 | Pig | Generalist | 6 | Germany |
| SRR12302208 | BfRCA06058 | 10272 | ST-403 | Pig | Pig | 11 | Germany |

| | | | | | | | |
|---|---|---|---|---|---|---|---|
| SRR12302207 | BfRCA06059 | 435 | ST-403 | Pig | Pig | 11 | Germany |
| SRR12302206 | BfRCA06089 | 435 | ST-403 | Pig | Pig | 11 | Germany |
| SRR12302205 | BfRCA06331 | 21 | ST-21 | Pig | Generalist | 6 | Germany |
| SRR12302204 | BfRCA07435 | 1775 | ST-403 | Pig | Pig | 11 | Germany |
| SRR12302203 | BfRCA07767 | 1777 | ST-403 | Pig | Pig | 11 | Germany |
| SRR12302202 | BfRCA08326 | 1777 | ST-403 | Pig | Pig | 11 | Germany: Federal State L |
| SRR12302201 | BfRCA08665 | 583 | ST-45 | Chicken | Generalist | 8 | Germany: Federal State J |
| SRR12302199 | BfRCA10009 | 21 | ST-21 | Chicken | Generalist | 6 | Germany: Federal State B |
| SRR12302198 | BfRCA10050 | 51 | ST-443 | Chicken | Chicken | 1 | Germany: Federal State K |
| SRR12302197 | BfRCA10051 | 48 | ST-48 | Cattle | Generalist | 3 | Germany: Federal State H |
| SRR12302196 | BfRCA10084 | 1073 | ST-354 | Chicken | Chicken | 1 | Germany: Federal State J |
| SRR12302195 | BfRCA10129 | 53 | ST-21 | Chicken | Generalist | 6 | Germany: Federal State D |
| SRR12302194 | BfRCA10142 | 6409 | ST-1034 | Chicken | Chicken | 5 | Germany: Federal State K |
| SRR12302193 | BfRCA10153 | 21 | ST-21 | Chicken | Generalist | 6 | Germany: Federal State A |
| SRR12302192 | BfRCA10170 | 257 | ST-257 | Chicken | Chicken | 9 | Germany: Federal State A |
| SRR12302191 | BfRCA10173 | 48 | ST-48 | Chicken | Generalist | 3 | Germany: Federal State A |
| SRR12302190 | BfRCA10176 | 4754 | unknown | Chicken | Chicken | 5 | Germany: Federal State C |
| SRR12302188 | BfRCA10204 | 61 | ST-61 | Cattle | Cattle | 10 | Germany: Federal State G |
| SRR12302187 | BfRCA10221 | 48 | ST-48 | Chicken | Generalist | 3 | Germany: Federal State N |
| SRR12302186 | BfRCA10231 | 429 | ST-48 | Chicken | Generalist | 3 | Germany: Federal State E |
| SRR12302185 | BfRCA10238 | 51 | ST-443 | Chicken | Chicken | 1 | Germany: Federal State J |
| SRR12302184 | BfRCA10257 | 354 | ST-354 | Chicken | Chicken | 1 | Germany: Federal State H |
| SRR12302183 | BfRCA10272 | 3100 | unknown | Cattle | unknown | 3 | Germany: Federal State H |
| SRR12302182 | BfRCA10285 | 19 | ST-21 | Cattle | Generalist | 2 | Germany: Federal State H |
| SRR12302181 | BfRCA10287 | 273 | ST-206 | Cattle | Generalist | 3 | Germany: Federal State H |
| SRR12302180 | BfRCA10303 | 38 | ST-48 | Cattle | Generalist | 3 | Germany: Federal State O |
| SRR12302179 | BfRCA10338 | 45 | ST-45 | Chicken | Generalist | 8 | Germany: Federal State A |
| SRR12302177 | BfRCA10393 | 2274 | unknown | Chicken | unknown | 1 | Germany: Federal State D |
| SRR12302176 | BfRCA10394 | 46 | ST-206 | Chicken | Generalist | 3 | Germany: Federal State M |
| SRR12302175 | BfRCA10443 | 21 | ST-21 | Cattle | Generalist | 6 | Germany: Federal State O |
| SRR12302174 | BfRCA10483 | 38 | ST-48 | Cattle | Generalist | 3 | Germany: Federal State M |
| SRR12302173 | BfRCA10491 | 658 | ST-658 | Chicken | unknown | 1 | Germany: Federal State M |
| SRR12302172 | BfRCA10492 | 50 | ST-21 | Chicken | Generalist | 2 | Germany: Federal State J |
| SRR12302171 | BfRCA10572 | 2314 | ST-1034 | Chicken | Chicken | 5 | Germany: Federal State D |
| SRR12302170 | BfRCA10615 | 19 | ST-21 | Cattle | Generalist | 2 | Germany: Federal State J |

| | | | | | | | |
|---|---|---|---|---|---|---|---|
| SRR12302169 | BfRCA10644 | 1911 | unknown | Chicken | unknown | 1 | Germany: Federal State J |
| SRR12302168 | BfRCA10649 | 45 | ST-45 | Chicken | Generalist | 8 | Germany: Federal State J |
| SRR12302166 | BfRCA10651 | 7958 | unknown | Chicken | unknown | 5 | Germany: Federal State C |
| SRR12302137 | BfRCA10667 | 21 | ST-21 | Cattle | Generalist | 6 | Germany: Federal State H |
| SRR12302136 | BfRCA10675 | 3100 | unknown | Cattle | unknown | 3 | Germany: Federal State H |
| SRR12302135 | BfRCA10677 | 7041 | ST-21 | Cattle | Generalist | 3 | Germany: Federal State H |
| SRR12302134 | BfRCA10717 | 257 | ST-257 | Cattle | Chicken | 9 | Germany: Federal State H |
| SRR12302133 | BfRCA10738 | 10273 | ST-61 | Cattle | Cattle | 10 | Germany: Federal State H |
| SRR12302132 | BfRCA10753 | 21 | ST-21 | Cattle | Generalist | 6 | Germany: Federal State H |
| SRR12302131 | BfRCA10776b | 403 | ST-403 | Cattle | Pig | 11 | Germany: Federal State H |
| SRR12302130 | BfRCA10820 | 51 | ST-443 | Cattle | Chicken | 1 | Germany: Federal State A |
| SRR12302129 | BfRCA10827 | 21 | ST-21 | Cattle | Generalist | 6 | Germany: Federal State M |
| SRR12302127 | BfRCA10834 | 6728 | ST-353 | Chicken | Chicken | 1 | Germany: Federal State D |
| SRR12302126 | BfRCA10850 | 233 | ST-45 | Chicken | Generalist | 8 | Germany: Federal State J |
| SRR12302125 | BfRCA10877 | 48 | ST-48 | Cattle | Generalist | 3 | Germany: Federal State M |
| SRR12302124 | BfRCA10905 | 48 | ST-48 | Cattle | Generalist | 3 | Germany: Federal State A |
| SRR12302123 | BfRCA10926 | 21 | ST-21 | Cattle | Generalist | 6 | Germany: Federal State A |
| SRR12302122 | BfRCA10942 | 206 | ST-206 | Cattle | Generalist | 3 | Germany: Federal State J |
| SRR12302121 | BfRCA10958 | 5970 | unknown | Chicken | unknown | 5 | Germany: Federal State H |
| SRR12302120 | BfRCA10959 | 19 | ST-21 | Cattle | Generalist | 2 | Germany: Federal State O |
| SRR12302119 | BfRCA10961 | 233 | ST-45 | Chicken | Generalist | 8 | Germany: Federal State D |
| SRR12302118 | BfRCA10962 | 50 | ST-21 | Chicken | Generalist | 2 | Germany: Federal State D |
| SRR12302116 | BfRCA10963 | 61 | ST-61 | Cattle | Cattle | 10 | Germany: Federal State A |
| SRR12302115 | BfRCA10977 | 2156 | unknown | Cattle | unknown | 3 | Germany: Federal State B |
| SRR12302114 | BfRCA10984 | 21 | ST-21 | Cattle | Generalist | 6 | Germany: Federal State O |
| SRR12302086 | BfRCA10988 | 50 | ST-21 | Chicken | Generalist | 2 | Germany: Federal State P |
| SRR12302085 | BfRCA11042 | 61 | ST-61 | Cattle | Cattle | 10 | Germany: Federal State P |
| SRR12302084 | BfRCA11044 | 2274 | unknown | Chicken | unknown | 1 | Germany: Federal State J |
| SRR12302083 | BfRCA11054 | 2274 | unknown | Chicken | unknown | 1 | Germany: Federal State J |
| SRR12302082 | BfRCA11065 | 3098 | unknown | Cattle | unknown | 12 | Germany: Federal State D |
| SRR12302081 | BfRCA11066 | 10275 | ST-21 | Cattle | Generalist | 6 | Germany: Federal State H |
| SRR12302080 | BfRCA11083 | 257 | ST-257 | Cattle | Chicken | 9 | Germany: Federal State H |
| SRR12302078 | BfRCA11096 | 21 | ST-21 | Cattle | Generalist | 6 | Germany: Federal State H |
| SRR12302077 | BfRCA11177 | 21 | ST-21 | Cattle | Generalist | 6 | Germany: Federal State A |
| SRR12302076 | BfRCA11178 | 21 | ST-21 | Cattle | Generalist | 6 | Germany: Federal State A |

| | | | | | | | |
|---|---|---|---|---|---|---|---|
| SRR12302075 | BfRCA11192 | 10276 | unknown | Cattle | unknown | 12 | Germany: Federal State J |
| SRR12302074 | BfRCA11199 | 2274 | unknown | Chicken | unknown | 1 | Germany: Federal State B |
| SRR12302073 | BfRCA11202 | 21 | ST-21 | Chicken | Generalist | 6 | Germany: Federal State O |
| SRR12302072 | BfRCA11209 | 42 | ST-42 | Cattle | Cattle | 4 | Germany: Federal State A |
| SRR12302071 | BfRCA11214 | 61 | ST-61 | Cattle | Cattle | 10 | Germany: Federal State J |
| SRR12302070 | BfRCA11215 | 19 | ST-21 | Cattle | Generalist | 2 | Germany: Federal State J |
| SRR12302069 | BfRCA11219 | 48 | ST-48 | Chicken | Generalist | 3 | Germany: Federal State B |
| SRR12302067 | BfRCA11234 | 55 | ST-403 | Cattle | Pig | 11 | Germany: Federal State M |
| SRR12302066 | BfRCA11258 | 10277 | unknown | Chicken | unknown | 1 | Germany: Federal State A |
| SRR12302065 | BfRCA11315 | 21 | ST-21 | Cattle | Generalist | 6 | Germany: Federal State J |
| SRR12302064 | BfRCA11319 | 61 | ST-61 | Cattle | Cattle | 10 | Germany: Federal State J |
| SRR12302063 | BfRCA11330 | 45 | ST-45 | Cattle | Generalist | 8 | Germany: Federal State J |
| SRR12302062 | BfRCA11331 | 21 | ST-21 | Cattle | Generalist | 6 | Germany: Federal State J |
| SRR12302061 | BfRCA11344 | 61 | ST-61 | Cattle | Cattle | 10 | Germany: Federal State H |
| SRR12302060 | BfRCA11346 | 61 | ST-61 | Cattle | Cattle | 10 | Germany: Federal State H |
| SRR12302059 | BfRCA11347 | 21 | ST-21 | Cattle | Generalist | 6 | Germany: Federal State H |
| SRR12302058 | BfRCA11352 | 38 | ST-48 | Cattle | Generalist | 3 | Germany: Federal State H |
| SRR12302056 | BfRCA11375 | 50 | ST-21 | Cattle | Generalist | 2 | Germany: Federal State B |
| SRR12302055 | BfRCA11386 | 7515 | ST-42 | Cattle | Cattle | 4 | Germany: Federal State A |
| SRR12302054 | BfRCA11387 | 10278 | ST-42 | Cattle | Cattle | 4 | Germany: Federal State A |
| SRR12302053 | BfRCA11388 | 61 | ST-61 | Cattle | Cattle | 10 | Germany: Federal State A |
| SRR12302052 | BfRCA11390 | 50 | ST-21 | Chicken | Generalist | 2 | Germany: Federal State A |
| SRR12302051 | BfRCA11392 | 21 | ST-21 | Chicken | Generalist | 6 | Germany: Federal State A |
| SRR12302050 | BfRCA11421 | 45 | ST-45 | Chicken | Generalist | 8 | Germany: Federal State B |
| SRR12302049 | BfRCA11438 | 904 | ST-607 | Chicken | unknown | 1 | Germany: Federal State G |
| SRR12302048 | BfRCA11498 | 21 | ST-21 | Cattle | Generalist | 6 | Germany: Federal State J |
| SRR12302047 | BfRCA11566 | 2153 | unknown | Chicken | unknown | 1 | Germany: Federal State J |
| SRR12302044 | BfRCA11567 | 3766 | unknown | Chicken | unknown | 5 | Germany: Federal State J |
| SRR12302043 | BfRCA11573 | 464 | ST-464 | Cattle | unknown | 1 | Germany: Federal State J |
| SRR12302042 | BfRCA11581 | 45 | ST-45 | Cattle | Generalist | 8 | Germany: Federal State D |
| SRR12302041 | BfRCA11590 | 583 | ST-45 | Cattle | Generalist | 8 | Germany: Federal State K |
| SRR12302040 | BfRCA11610 | 933 | ST-403 | Cattle | Pig | 11 | Germany: Federal State P |
| SRR12302039 | BfRCA11627 | 21 | ST-21 | Cattle | Generalist | 6 | Germany: Federal State B |
| SRR12302037 | BfRCA11629 | 6461 | ST-353 | Chicken | Chicken | 1 | Germany: Federal State B |
| SRR12302036 | BfRCA11633 | 19 | ST-21 | Chicken | Generalist | 2 | Germany: Federal State K |

| | | | | | | | |
|---|---|---|---|---|---|---|---|
| SRR12302035 | BfRCA11654 | 50 | ST-21 | Cattle | Generalist | 2 | Germany: Federal State B |
| SRR12302034 | BfRCA11663 | 586 | unknown | Cattle | unknown | 4 | Germany: Federal State B |
| SRR12302032 | BfRCA11664 | 10280 | unknown | Chicken | unknown | 5 | Germany: Federal State K |
| SRR12302031 | BfRCA11665 | 21 | ST-21 | Cattle | Generalist | 6 | Germany: Federal State K |
| SRR12302030 | BfRCA11667 | 21 | ST-21 | Cattle | Generalist | 6 | Germany: Federal State K |
| SRR12302029 | BfRCA11700 | 1301 | ST-692 | Chicken | unknown | 5 | Germany: Federal State P |
| SRR12302028 | BfRCA11706 | 10281 | ST-21 | Cattle | Generalist | 6 | Germany: Federal State J |
| SRR12302027 | BfRCA11713 | 10282 | unknown | Cattle | unknown | 12 | Germany: Federal State J |
| SRR12302026 | BfRCA11722 | 21 | ST-21 | Cattle | Generalist | 6 | Germany: Federal State A |
| SRR12302025 | BfRCA11723 | 19 | ST-21 | Pig | Generalist | 2 | Germany: Federal State O |
| SRR12302024 | BfRCA11724 | 61 | ST-61 | Cattle | Cattle | 10 | Germany: Federal State O |
| SRR12302023 | BfRCA11725 | 432 | ST-61 | Cattle | Cattle | 10 | Germany: Federal State O |
| SRR12302021 | BfRCA11846 | 44 | ST-21 | Chicken | Generalist | 2 | Germany: Federal State I |
| SRR12302020 | BfRCA11848b | 51 | ST-443 | Cattle | Chicken | 1 | Germany: Federal State A |
| SRR12302019 | BfRCA11849 | 356 | ST-353 | Chicken | Chicken | 1 | Germany: Federal State A |
| SRR12302018 | BfRCA11850 | 19 | ST-21 | Cattle | Generalist | 2 | Germany: Federal State A |
| SRR12302017 | BfRCA11851 | 2274 | unknown | Chicken | unknown | 1 | Germany: Federal State A |
| SRR12302016 | BfRCA11852 | 21 | ST-21 | Cattle | Generalist | 6 | Germany: Federal State H |
| SRR12302015 | BfRCA11853 | 50 | ST-21 | Cattle | Generalist | 2 | Germany: Federal State A |
| SRR12302014 | BfRCA11884 | 583 | ST-45 | Chicken | Generalist | 8 | Germany: Federal State J |
| SRR12302013 | BfRCA11888 | 2066 | ST-52 | Chicken | unknown | 1 | Germany: Federal State B |
| SRR12302012 | BfRCA11917 | 42 | ST-42 | Chicken | Cattle | 4 | Germany: Federal State J |
| SRR12302010 | BfRCA11926 | 257 | ST-257 | Chicken | Chicken | 9 | Germany: Federal State J |
| SRR12302009 | BfRCA11946 | 50 | ST-21 | Chicken | Generalist | 2 | Germany: Federal State A |
| SRR12302008 | BfRCA12057 | 21 | ST-21 | Chicken | Generalist | 6 | Germany: Federal State B |
| SRR12302007 | BfRCA12154 | 5798 | unknown | Chicken | unknown | 5 | Germany: Federal State B |
| SRR12302006 | BfRCA12659 | 21 | ST-21 | Chicken | Generalist | 6 | Germany: Federal State B |
| SRR12302005 | BfRCA12662 | 5840 | ST-353 | Chicken | Chicken | 1 | Germany: Federal State B |
| SRR12302004 | BfRCA12663 | 50 | ST-21 | Chicken | Generalist | 2 | Germany: Federal State B |
| SRR12302003 | BfRCA12891 | 464 | ST-464 | Chicken | unknown | 1 | Germany: Federal State B |
| SRR12302002 | BfRCA12978 | 2254 | ST-257 | Chicken | Chicken | 9 | Germany: Federal State N |
| SRR12302001 | BfRCA13157 | 21 | ST-21 | Chicken | Generalist | 6 | Germany: Federal State B |
| SRR12301999 | BfRCA13162 | 6175 | ST-21 | Chicken | Generalist | 3 | Germany: Federal State B |
| SRR12301998 | BfRCA13163 | 46 | ST-206 | Chicken | Generalist | 3 | Germany: Federal State A |
| SRR12301997 | BfRCA13168 | 3628 | ST-443 | Chicken | Chicken | 1 | Germany: Federal State A |

| | | | | | | | |
|---|---|---|---|---|---|---|---|
| SRR12301996 | BfRCA13169 | 10283 | ST-443 | Chicken | Chicken | 1 | Germany: Federal State A |
| SRR12301995 | BfRCA13171 | 1519 | ST-21 | Chicken | Generalist | 2 | Germany: Federal State J |
| SRR12301994 | BfRCA13189 | 21 | ST-21 | Cattle | Generalist | 6 | Germany: Federal State M |
| SRR12301993 | BfRCA13199 | 861 | ST-21 | Cattle | Generalist | 6 | Germany: Federal State O |
| SRR12301992 | BfRCA13206 | 5103 | ST-22 | Cattle | unknown | 4 | Germany: Federal State A |
| SRR12301991 | BfRCA13207 | 42 | ST-42 | Cattle | Cattle | 4 | Germany: Federal State A |
| SRR12301990 | BfRCA13233b | 403 | ST-403 | Pig | Pig | 11 | Germany |
| SRR12301988 | BfRCA13265 | 354 | ST-354 | Pig | Chicken | 1 | Germany |
| SRR12301987 | BfRCA13281 | 403 | ST-403 | Pig | Pig | 11 | Germany: Federal State J |
| SRR12301986 | BfRCA13282 | 403 | ST-403 | Pig | Pig | 11 | Germany: Federal State J |
| SRR12301985 | BfRCA13292 | 22 | ST-22 | Cattle | unknown | 4 | Germany: Federal State J |
| SRR12301984 | BfRCA13298 | 22 | ST-22 | Cattle | unknown | 4 | Germany: Federal State J |
| SRR12301983 | BfRCA13324 | 10284 | unknown | Cattle | unknown | 3 | Germany: Federal State J |
| SRR12301982 | BfRCA13330 | 441 | unknown | Cattle | unknown | 13 | Germany: Federal State J |
| SRR12301981 | BfRCA13394 | 61 | ST-61 | Cattle | Cattle | 10 | Germany: Federal State A |
| SRR12301980 | BfRCA13398 | 2274 | unknown | Chicken | unknown | 1 | Germany: Federal State H |
| SRR12301979 | BfRCA13453 | 38 | ST-48 | Cattle | Generalist | 3 | Germany: Federal State J |
| SRR12301977 | BfRCA13463 | 50 | ST-21 | Chicken | Generalist | 2 | Germany: Federal State J |
| SRR12301976 | BfRCA13512 | 21 | ST-21 | Cattle | Generalist | 6 | Germany: Federal State J |
| SRR12301975 | BfRCA13514 | 21 | ST-21 | Cattle | Generalist | 6 | Germany: Federal State J |
| SRR12301974 | BfRCA13527 | 1709 | ST-1034 | Chicken | Chicken | 5 | Germany: Federal State N |
| SRR12301973 | BfRCA13538 | 10285 | ST-61 | Cattle | unknown | 10 | Germany: Federal State I |
| SRR12301972 | BfRCA13539 | 206 | ST-206 | Cattle | Generalist | 3 | Germany: Federal State I |
| SRR12301971 | BfRCA13541 | 21 | ST-21 | Cattle | Generalist | 6 | Germany: Federal State I |
| SRR12301970 | BfRCA13564 | 61 | ST-61 | Cattle | Cattle | 10 | Germany: Federal State J |
| SRR12301969 | BfRCA13582 | 38 | ST-48 | Cattle | Generalist | 3 | Germany: Federal State I |
| SRR12301968 | BfRCA13613 | 42 | ST-42 | Cattle | Cattle | 4 | Germany: Federal State J |
| SRR12301966 | BfRCA13729 | 3155 | ST-354 | Pig | Chicken | 1 | Germany |
| SRR12301965 | BfRCA13756 | 403 | ST-403 | Cattle | Pig | 11 | Germany: Federal State J |
| SRR12301964 | BfRCA13758 | 21 | ST-21 | Cattle | Generalist | 6 | Germany: Federal State J |
| SRR12301963 | BfRCA13795 | 19 | ST-21 | Chicken | Generalist | 2 | Germany: Federal State H |
| SRR12301962 | BfRCA13811 | 1459 | ST-21 | Cattle | Generalist | 6 | Germany: Federal State J |
| SRR12301961 | BfRCA13816 | 257 | ST-257 | Cattle | Chicken | 9 | Germany: Federal State J |
| SRR12301960 | BfRCA13821 | 42 | ST-42 | Cattle | Cattle | 4 | Germany: Federal State J |
| SRR12301959 | BfRCA13824 | 10286 | ST-61 | Cattle | unknown | 10 | Germany: Federal State J |

| | | | | | | | |
|---|---|---|---|---|---|---|---|
| SRR12302281 | BfRCA13826 | 19 | ST-21 | Cattle | Generalist | 2 | Germany: Federal State J |
| SRR12302280 | BfRCA13835 | 354 | ST-354 | Cattle | Chicken | 1 | Germany: Federal State J |
| SRR12302278 | BfRCA13836 | 21 | ST-21 | Cattle | Generalist | 6 | Germany: Federal State J |
| SRR12302277 | BfRCA13838 | 21 | ST-21 | Cattle | Generalist | 6 | Germany: Federal State J |
| SRR12302276 | BfRCA13918 | 1003 | ST-45 | Chicken | Generalist | 8 | Germany: Federal State B |
| SRR12302275 | BfRCA13937 | 1519 | ST-21 | Chicken | Generalist | 2 | Germany: Federal State K |
| SRR12302274 | BfRCA13939 | 607 | ST-607 | Chicken | unknown | 1 | Germany: Federal State A |
| SRR12302273 | BfRCA14088 | 48 | ST-48 | Chicken | Generalist | 3 | Germany: Federal State B |
| SRR12302272 | BfRCA14109 | 45 | ST-45 | Chicken | Generalist | 8 | Germany: Federal State M |
| SRR12302271 | BfRCA14180 | 354 | ST-354 | Chicken | Chicken | 1 | Germany: Federal State B |
| SRR12302270 | BfRCA14181 | 2066 | ST-52 | Chicken | unknown | 1 | Germany: Federal State B |
| SRR12302269 | BfRCA14304 | 2304 | unknown | Chicken | unknown | 3 | Germany: Federal State I |
| SRR12302267 | BfRCA14323 | 1519 | ST-21 | Chicken | Generalist | 2 | Germany: Federal State B |
| SRR10103069, SRR101( | BfRCA14430 | 44 | ST-21 | Chicken | Generalist | 2 | Germany |
| SRR12302266 | BfRCA14435 | 1519 | ST-21 | Chicken | Generalist | 2 | Germany: Federal State B |
| SRR12302265 | BfRCA14444 | 2274 | unknown | Chicken | unknown | 1 | Germany: Federal State J |
| SRR12302264 | BfRCA14553 | 44 | ST-21 | Chicken | Generalist | 2 | Germany: Federal State P |
| SRR12302263 | BfRCA14579 | 977 | ST-1034 | Chicken | Chicken | 5 | Germany: Federal State J |
| SRR12302262 | BfRCA14704 | 538 | ST-45 | Chicken | Generalist | 8 | Germany: Federal State P |
| SRR12302261 | BfRCA14734 | 977 | ST-1034 | Chicken | Chicken | 5 | Germany: Federal State J |
| SRR12302260 | BfRCA14814 | 2275 | ST-52 | Chicken | unknown | 1 | Germany: Federal State H |
| SRR12302259 | BfRCA14836 | 464 | ST-464 | Chicken | unknown | 1 | Germany: Federal State J |
| SRR12302258 | BfRCA14940 | 267 | ST-283 | Chicken | unknown | 8 | Germany: Federal State F |
| SRR12302255 | BfRCA14957 | 50 | ST-21 | Chicken | Generalist | 2 | Germany: Federal State O |
| SRR12302254 | BfRCA14962 | 2275 | ST-52 | Chicken | unknown | 1 | Germany: Federal State J |
| SRR12302253 | BfRCA14988 | 8334 | ST-353 | Chicken | Chicken | 1 | Germany: Federal State B |
| SRR12302252 | BfRCA14993 | 2275 | ST-52 | Chicken | unknown | 1 | Germany: Federal State J |
| SRR12302251 | BfRCA15004 | 2254 | ST-257 | Chicken | Chicken | 9 | Germany: Federal State P |
| SRR12302250 | BfRCA15023b | 1846 | ST-403 | Pig | Pig | 11 | Germany |
| SRR12302249 | BfRCA15024 | 1846 | ST-403 | Pig | Pig | 11 | Germany |
| SRR12302248 | BfRCA15054 | 1846 | ST-403 | Pig | Pig | 11 | Germany |
| SRR12302247 | BfRCA15085 | 9351 | unknown | Chicken | unknown | 5 | Germany: Federal State P |
| SRR12302246 | BfRCA15095 | 3335 | ST-206 | Pig | Generalist | 3 | Germany |
| SRR12302244 | BfRCA15119 | 45 | ST-45 | Chicken | Generalist | 8 | Germany: Federal State H |
| SRR12302243 | BfRCA15155 | 400 | ST-353 | Chicken | Chicken | 1 | Germany: Federal State M |

| | | | | | | | |
|---|---|---|---|---|---|---|---|
| SRR12302242 | BfRCA15166 | 9366 | ST-403 | Pig | Pig | 11 | Germany: Federal State E |
| SRR12302241 | BfRCA15240 | 257 | ST-257 | Chicken | Chicken | 9 | Germany: Federal State E |
| SRR12302240 | BfRCA15256 | 1942 | unknown | Pig | unknown | 11 | Germany |
| SRR12302239 | BfRCA15265 | 3628 | ST-443 | Chicken | Chicken | 1 | Germany: Federal State A |
| SRR12302238 | BfRCA15279 | 21 | ST-21 | Pig | Generalist | 6 | Germany |
| SRR12302237 | BfRCA15284 | 400 | ST-353 | Chicken | Chicken | 1 | Germany: Federal State M |
| SRR12302236 | BfRCA15332 | 1775 | ST-403 | Pig | Pig | 11 | Germany: Federal State H |
| SRR12302235 | BfRCA15395 | 464 | ST-464 | Chicken | unknown | 1 | Germany: Federal State P |
| SRR12302233 | IMT468 | 10287 | ST-403 | Pig | Pig | 11 | Germany |
| SRR12302232 | IMT538 | 1775 | ST-403 | Pig | Pig | 11 | Germany |
| SRR12302231 | IMT541 | 403 | ST-403 | Pig | Pig | 11 | Germany |
| SRR5209454 | SRR5209454 | 61 | ST-61 | Horse | Cattle | 10 | Canada: Alberta |
| SRR5209455 | SRR5209455 | 61 | ST-61 | Horse | Cattle | 10 | Canada: Alberta |
| SRR5209456 | SRR5209456 | 1244 | ST-61 | Cattle | Cattle | 10 | Canada: Alberta |
| SRR5209457 | SRR5209457 | 1244 | ST-61 | Cattle | Cattle | 10 | Canada: Alberta |
| SRR5209458 | SRR5209458 | 45 | ST-45 | Water | Generalist | 8 | Canada: South Nation Watershed\, Ontario |
| SRR5209459 | SRR5209459 | 2539 | ST-177 | Raccoon | unknown | 7 | Canada: Ontario |
| SRR5209460 | SRR5209460 | 137 | ST-45 | Raccoon | Generalist | 8 | Canada: Ontario |
| SRR5209461 | SRR5209461 | 137 | ST-45 | Raccoon | Generalist | 8 | Canada: Ontario |
| SRR5209462 | SRR5209462 | 137 | ST-45 | Raccoon | Generalist | 8 | Canada: Ontario |
| SRR5209463 | SRR5209463 | 45 | ST-45 | Water | Generalist | 8 | Canada: Ontario |
| SRR5209464 | SRR5209464 | 61 | ST-61 | Cattle | Cattle | 10 | Canada: Alberta |
| SRR5209465 | SRR5209465 | 1244 | ST-61 | Cattle | Cattle | 10 | Canada: Alberta |
| SRR5209466 | SRR5209466 | 45 | ST-45 | Water | Generalist | 8 | Canada: Ontario |
| SRR5209467 | SRR5209467 | 682 | ST-682 | Water | unknown | 7 | Canada: Alberta |
| SRR5209468 | SRR5209468 | 45 | ST-45 | Water | Generalist | 8 | Canada: Ontario |
| SRR5209469 | SRR5209469 | 10289 | ST-177 | Water | unknown | 7 | Canada: Quebec |
| SRR5209470 | SRR5209470 | 682 | ST-682 | Water | unknown | 7 | Canada: Ontario |
| SRR5209471 | SRR5209471 | 45 | ST-45 | Water | Generalist | 8 | Canada: Ontario |
| SRR5209472 | SRR5209472 | 10290 | ST-42 | Water | Cattle | 4 | Canada: Ontario |
| SRR5209473 | SRR5209473 | 132 | ST-508 | Cattle | unknown | 14 | Canada: Alberta |
| SRR5209474 | SRR5209474 | 132 | ST-508 | Cattle | unknown | 14 | Canada: Alberta |
| SRR5209475 | SRR5209475 | 42 | ST-42 | Water | Cattle | 4 | Canada: Ontario |
| SRR5209476 | SRR5209476 | 45 | ST-45 | Water | Generalist | 8 | Canada: Alberta |
| SRR5209477 | SRR5209477 | 45 | ST-45 | Water | Generalist | 8 | Canada: Ontario |

| | | | | | | | |
|---|---|---|---|---|---|---|---|
| SRR5209478 | SRR5209478 | 137 | ST-45 | Water | Generalist | 8 | Canada: Alberta |
| SRR5209479 | SRR5209479 | 45 | ST-45 | Water | Generalist | 8 | Canada: Ontario |
| SRR5209480 | SRR5209480 | 682 | ST-682 | Water | unknown | 7 | Canada: New Brunswick |
| SRR5209481 | SRR5209481 | 45 | ST-45 | Chicken | Generalist | 8 | Canada: Alberta |
| SRR5209482 | SRR5209482 | 45 | ST-45 | Duck | Generalist | 8 | Canada: Alberta |
| SRR5209483 | SRR5209483 | 45 | ST-45 | Dog | Generalist | 8 | Canada: Alberta |
| SRR5209484 | SRR5209484 | 45 | ST-45 | Goose | Generalist | 8 | Canada: Alberta |
| SRR5209485 | SRR5209485 | 45 | ST-45 | Water | Generalist | 8 | Canada: Alberta |
| SRR5209486 | SRR5209486 | 1244 | ST-61 | Cattle | Cattle | 10 | Canada: Alberta |
| SRR5209487 | SRR5209487 | 682 | ST-682 | Water | unknown | 7 | Canada: Ontario |
| SRR5209488 | SRR5209488 | 5128 | ST-682 | Water | unknown | 7 | Canada: New Brunswick |
| SRR5209489 | SRR5209489 | 45 | ST-45 | Sheep | Generalist | 8 | Canada: Alberta |
| SRR5209490 | SRR5209490 | 45 | ST-45 | Cattle | Generalist | 8 | Canada: Alberta |
| SRR5209491 | SRR5209491 | 45 | ST-45 | Sheep | Generalist | 8 | Canada: Alberta |
| SRR5209492 | SRR5209492 | 61 | ST-61 | Cattle | Cattle | 10 | Canada: Alberta |
| SRR5209493 | SRR5209493 | 459 | ST-42 | Cattle | Cattle | 4 | Canada: Alberta |
| SRR5209494 | SRR5209494 | 48 | ST-48 | Chicken | Generalist | 3 | Canada: Alberta |
| SRR5209495 | SRR5209495 | 48 | ST-48 | Chicken | Generalist | 3 | Canada: Alberta |
| SRR5209496 | SRR5209496 | 45 | ST-45 | Sewage | Generalist | 8 | Canada: Alberta |
| SRR5209497 | SRR5209497 | 137 | ST-45 | Water | Generalist | 8 | Canada: Alberta |
| SRR5209498 | SRR5209498 | 459 | ST-42 | Cat | Cattle | 4 | Canada: Alberta |
| SRR5209499 | SRR5209499 | 132 | ST-508 | Sewage | unknown | 14 | Canada: Alberta |
| SRR5209500 | SRR5209500 | 806 | ST-21 | Sheep | Generalist | 6 | Canada: Alberta |
| SRR5209501 | SRR5209501 | 61 | ST-61 | Cattle | Cattle | 10 | Canada: Alberta |
| SRR5209502 | SRR5209502 | 45 | ST-45 | Cattle | Generalist | 8 | Canada: Alberta |
| SRR5209503 | SRR5209503 | 61 | ST-61 | Water | Cattle | 10 | Canada: Alberta |
| SRR5209504 | SRR5209504 | 132 | ST-508 | Cattle | unknown | 14 | Canada: Alberta |
| SRR5209505 | SRR5209505 | 132 | ST-508 | Cattle | unknown | 14 | Canada: Alberta |
| SRR5209506 | SRR5209506 | 459 | ST-42 | Water | Cattle | 4 | Canada: Alberta |
| SRR5209507 | SRR5209507 | 262 | ST-21 | Human | Generalist | 6 | Canada: Alberta |
| SRR5209508 | SRR5209508 | 61 | ST-61 | Human | Cattle | 10 | Canada: Alberta |
| SRR5209509 | SRR5209509 | 10291 | ST-48 | Human | unknown | 3 | Canada: Alberta |
| SRR5209510 | SRR5209510 | 45 | ST-45 | Human | Generalist | 8 | Canada: Alberta |
| SRR5209511 | SRR5209511 | 19 | ST-21 | Human | Generalist | 2 | Canada: Alberta |
| SRR5209512 | SRR5209512 | 45 | ST-45 | Human | Generalist | 8 | Canada: Alberta |

| | | | | | | | |
|---|---|---|---|---|---|---|---|
| SRR5209513 | SRR5209513 | 45 | ST-45 | Human | Generalist | 8 | Canada: Alberta |
| SRR5209514 | SRR5209514 | 48 | ST-48 | Human | Generalist | 3 | Canada: Alberta |
| SRR5209515 | SRR5209515 | 10291 | ST-48 | Human | unknown | 3 | Canada: Ontario |
| SRR5209516 | SRR5209516 | 48 | ST-48 | Human | Generalist | 3 | Canada: Alberta |
| SRR5209517 | SRR5209517 | 61 | ST-61 | Human | Cattle | 10 | Canada: Alberta |
| SRR5209518 | SRR5209518 | 45 | ST-45 | Human | Generalist | 8 | Canada: Alberta |
| SRR5209519 | SRR5209519 | 61 | ST-61 | Human | Cattle | 10 | Canada: Alberta |
| SRR5209520 | SRR5209520 | 50 | ST-21 | Human | Generalist | 2 | Canada: Alberta |
| SRR5209521 | SRR5209521 | 61 | ST-61 | Human | Cattle | 10 | Canada: Alberta |
| SRR5209522 | SRR5209522 | 45 | ST-45 | Chicken | Generalist | 8 | Canada: British Columbia |
| SRR5209523 | SRR5209523 | 267 | ST-283 | Chicken | unknown | 8 | Canada: British Columbia |
| SRR5209524 | SRR5209524 | 267 | ST-283 | Turkey | unknown | 8 | Canada: Ontario |
| SRR5209525 | SRR5209525 | 45 | ST-45 | Chicken | Generalist | 8 | Canada: Ontario |
| SRR5209526 | SRR5209526 | 21 | ST-21 | Chicken | Generalist | 6 | Canada: Ontario |
| SRR5209527 | SRR5209527 | 45 | ST-45 | Chicken | Generalist | 8 | Canada: Ontario |
| SRR5209528 | SRR5209528 | 45 | ST-45 | Cattle | Generalist | 8 | Canada: Ontario |
| SRR5209529 | SRR5209529 | 933 | ST-403 | Human | Pig | 11 | Canada: Ontario |
| SRR5209530 | SRR5209530 | 42 | ST-42 | Human | Cattle | 4 | Canada: Ontario |
| SRR5209531 | SRR5209531 | 21 | ST-21 | Human | Generalist | 6 | Canada: Ontario |
| SRR5209532 | SRR5209532 | 45 | ST-45 | Human | Generalist | 8 | Canada: Ontario |
| SRR5209533 | SRR5209533 | 45 | ST-45 | Chicken | Generalist | 8 | Canada: Ontario |
| SRR5209534 | SRR5209534 | 4080 | unknown | Water | unknown | 15 | Canada: Ontario |
| SRR5209535 | SRR5209535 | 1030 | unknown | Water | unknown | 15 | Canada: Quebec |
| SRR5209536 | SRR5209536 | 3112 | unknown | Water | unknown | 15 | Canada: Ontario |
| SRR5209537 | SRR5209537 | 1294 | unknown | Water | unknown | 15 | Canada: Alberta |
| SRR5209538 | SRR5209538 | 1030 | unknown | Water | unknown | 15 | Canada: Ontario |
| SRR5209539 | SRR5209539 | 3495 | unknown | Water | unknown | 15 | Canada: Ontario |
| SRR5209540 | SRR5209540 | 996 | unknown | Water | unknown | 15 | Canada: British Columbia |
| SRR5209541 | SRR5209541 | 996 | unknown | Water | unknown | 15 | Canada: Alberta |
| SRR5209542 | SRR5209542 | 693 | unknown | Water | unknown | 15 | Canada: Alberta |
| SRR5209543 | SRR5209543 | 699 | ST-692 | Water | unknown | 5 | Canada: Alberta |
| SRR5209544 | SRR5209544 | 991 | ST-692 | Water | unknown | 5 | Canada: British Columbia |
| SRR5209545 | SRR5209545 | 991 | ST-692 | Water | unknown | 5 | Canada: British Columbia |
| SRR5209546 | SRR5209546 | 6516 | unknown | Water | unknown | 15 | Canada: Alberta |
| SRR5209547 | SRR5209547 | 9353 | ST-1034 | Water | Chicken | 15 | Canada: British Columbia |

| | | | | | | | |
|---|---|---|---|---|---|---|---|
| SRR5209548 | SRR5209548 | 4071 | ST-1034 | Water | Chicken | 15 | Canada: Alberta |
| SRR5209549 | SRR5209549 | 693 | unknown | Water | unknown | 15 | Canada: Ontario |
| SRR5209550 | SRR5209550 | 991 | ST-692 | Water | unknown | 5 | Canada: British Columbia |
| SRR5209551 | SRR5209551 | 693 | unknown | Water | unknown | 15 | Canada: Alberta |
| SRR5209552 | SRR5209552 | 4071 | ST-1034 | Water | Chicken | 15 | Canada: Alberta |
| SRR5209553 | SRR5209553 | 5452 | unknown | Water | unknown | 15 | Canada: British Columbia |
| SRR5209554 | SRR5209554 | 991 | ST-692 | Water | unknown | 5 | Canada: British Columbia |
| SRR5209555 | SRR5209555 | 10293 | unknown | Water | unknown | 15 | Canada: Ontario |
| SRR5209556 | SRR5209556 | 5705 | unknown | Duck | unknown | 15 | Canada: Ontario |
| SRR5209557 | SRR5209557 | 5705 | unknown | Goose | unknown | 15 | Canada: Ontario |
| SRR5209558 | SRR5209558 | 995 | unknown | Water | unknown | 15 | Canada: British Columbia |
| SRR5209559 | SRR5209559 | 1030 | unknown | Water | unknown | 15 | Canada: Alberta |
| SRR5209560 | SRR5209560 | 3495 | unknown | Water | unknown | 15 | Canada: British Columbia |
| SRR5209561 | SRR5209561 | 710 | unknown | Goose | unknown | 15 | Canada: Alberta |
| SRR5209562 | SRR5209562 | 996 | unknown | Water | unknown | 15 | Canada: Alberta |
| SRR5209563 | SRR5209563 | 3112 | unknown | Water | unknown | 15 | Canada: Alberta |
| SRR5209564 | SRR5209564 | 10296 | unknown | Water | unknown | 15 | Canada: Alberta |
| SRR5209565 | SRR5209565 | 693 | unknown | Water | unknown | 15 | Canada: Alberta |
| SRR5209566 | SRR5209566 | 1206 | unknown | Goose | unknown | 15 | Canada: Alberta |
| SRR5209567 | SRR5209567 | 1206 | unknown | Goose | unknown | 15 | Canada: Alberta |
| SRR5209568 | SRR5209568 | 1206 | unknown | Goose | unknown | 15 | Canada: Alberta |
| SRR5209569 | SRR5209569 | 929 | ST-257 | Water | Chicken | 9 | Canada: Alberta |
| SRR5209570 | SRR5209570 | 929 | ST-257 | Water | Chicken | 9 | Canada: Alberta |
| SRR5209571 | SRR5209571 | 929 | ST-257 | Cattle | Chicken | 9 | Canada: Alberta |
| SRR5209572 | SRR5209572 | 929 | ST-257 | Cattle | Chicken | 9 | Canada: Alberta |
| SRR5209573 | SRR5209573 | 929 | ST-257 | Human | Chicken | 9 | Canada: Alberta |
| SRR5209574 | SRR5209574 | 929 | ST-257 | Human | Chicken | 9 | Canada: Alberta |
| SRR5209575 | SRR5209575 | 982 | ST-21 | Cattle | Generalist | 6 | Canada: Alberta |
| SRR5209576 | SRR5209576 | 21 | ST-21 | Cattle | Generalist | 6 | Canada: Alberta |
| SRR5209577 | SRR5209577 | 21 | ST-21 | Cattle | Generalist | 6 | Canada: Alberta |
| SRR5209578 | SRR5209578 | 21 | ST-21 | Cattle | Generalist | 6 | Canada: Alberta |
| SRR5209579 | SRR5209579 | 8 | ST-21 | Cattle | Generalist | 6 | Canada: Alberta |
| SRR5209580 | SRR5209580 | 8 | ST-21 | Cattle | Generalist | 6 | Canada: Alberta |
| SRR5209581 | SRR5209581 | 8 | ST-21 | Cattle | Generalist | 6 | Canada: Alberta |
| SRR5209582 | SRR5209582 | 8 | ST-21 | Cattle | Generalist | 6 | Canada: Alberta |

| | | | | | | | |
|---|---|---|---|---|---|---|---|
| SRR5209583 | SRR5209583 | 8 | ST-21 | Cattle | Generalist | 6 | Canada: Alberta |
| SRR5209584 | SRR5209584 | 3391 | ST-21 | Cattle | Generalist | 6 | Canada: Alberta |
| SRR5209585 | SRR5209585 | 3391 | ST-21 | Cattle | Generalist | 6 | Canada: Alberta |
| SRR5209586 | SRR5209586 | 3391 | ST-21 | Cattle | Generalist | 6 | Canada: Alberta |
| SRR5209587 | SRR5209587 | 8 | ST-21 | Cattle | Generalist | 6 | Canada: Alberta |
| SRR5209588 | SRR5209588 | 8 | ST-21 | Cattle | Generalist | 6 | Canada: Alberta |
| SRR5209589 | SRR5209589 | 982 | ST-21 | Chicken | Generalist | 6 | Canada: Alberta |
| SRR5209590 | SRR5209590 | 982 | ST-21 | Chicken | Generalist | 6 | Canada: Alberta |
| SRR5209591 | SRR5209591 | 922 | unknown | Cattle | unknown | 13 | Canada: Alberta |
| SRR5209592 | SRR5209592 | 45 | ST-45 | Sheep | Generalist | 8 | Canada: Alberta |
| SRR5209593 | SRR5209593 | 21 | ST-21 | Cattle | Generalist | 6 | Canada: Alberta |
| SRR5209594 | SRR5209594 | 922 | unknown | Cattle | unknown | 13 | Canada: Alberta |
| SRR5209595 | SRR5209595 | 8 | ST-21 | Cattle | Generalist | 6 | Canada: Alberta |
| SRR5209596 | SRR5209596 | 8 | ST-21 | Cattle | Generalist | 6 | Canada: Alberta |
| SRR5209597 | SRR5209597 | 922 | unknown | Cattle | unknown | 13 | Canada: Alberta |
| SRR5209598 | SRR5209598 | 8 | ST-21 | Cattle | Generalist | 6 | Canada: Alberta |
| SRR5209599 | SRR5209599 | 679 | ST-45 | Human | Generalist | 8 | Canada: Alberta |
| SRR5209600 | SRR5209600 | 982 | ST-21 | Human | Generalist | 6 | Canada: Alberta |
| SRR5209601 | SRR5209601 | 922 | unknown | Human | unknown | 13 | Canada: Alberta |
| SRR5209602 | SRR5209602 | 8 | ST-21 | Human | Generalist | 6 | Canada: Alberta |
| SRR5209603 | SRR5209603 | 21 | ST-21 | Human | Generalist | 6 | Canada: Alberta |
| SRR5209604 | SRR5209604 | 922 | unknown | Human | unknown | 13 | Canada: Alberta |
| SRR5209605 | SRR5209605 | 8 | ST-21 | Human | Generalist | 6 | Canada: Alberta |
| SRR5209606 | SRR5209606 | 8 | ST-21 | Human | Generalist | 6 | Canada: Alberta |
| SRR5209607 | SRR5209607 | 982 | ST-21 | Human | Generalist | 6 | Canada: Alberta |
| SRR5209608 | SRR5209608 | 982 | ST-21 | Human | Generalist | 6 | Canada: Alberta |
| SRR5209609 | SRR5209609 | 982 | ST-21 | Human | Generalist | 6 | Canada: Alberta |
| SRR5209610 | SRR5209610 | 982 | ST-21 | Human | Generalist | 6 | Canada: Alberta |
| SRR5209611 | SRR5209611 | 922 | unknown | Human | unknown | 13 | Canada: Alberta |
| SRR5209612 | SRR5209612 | 679 | ST-45 | Human | Generalist | 8 | Canada: Alberta |
| SRR5209613 | SRR5209613 | 2306 | ST-21 | Human | Generalist | 2 | Canada: Alberta |
| SRR5209614 | SRR5209614 | 46 | ST-206 | Human | Generalist | 3 | Canada: Alberta |
| SRR5209615 | SRR5209615 | 46 | ST-206 | Human | Generalist | 3 | Canada: Alberta |
| SRR5209616 | SRR5209616 | 19 | ST-21 | Human | Generalist | 2 | Canada: Alberta |
| SRR5209617 | SRR5209617 | 52 | ST-52 | Cattle | unknown | 1 | Canada: Ontario |

| | | | | | | | |
|---|---|---|---|---|---|---|---|
| SRR5209618 | SRR5209618 | 429 | ST-48 | Chicken | Generalist | 3 | Canada: Ontario |
| SRR5209619 | SRR5209619 | 918 | ST-48 | Human | Generalist | 3 | Canada: Ontario |
