## Supplementary material for "Genome-wide insights into population structure and host specificity of *Campylobacter jejuni*": Table S3

Table S3: Overview of cattle associated accessory genes and core gene variants identified by k-mer mapping.

| Reference Gene Nane[a] | Alias | Predicted Function | COG Family[b] | AA | # Cattle[c] | # Chicken[c] | # Pig[c] | # Generalist[c] | # Others[c] | Total | Core/Accessory[d] |
|---|---|---|---|---|---|---|---|---|---|---|---|
| NCTC13261_01393 | - | plasmid stabilization system protein, RelE/ParE family (HicA/HicB) | S | MKIVPTPKFKNEL | 55 | 1 | 0 | 59 | 7 | 122 | Accessory |
| NCTC13261_01392 | - | hypothetical protein | - | MNLEFSKETQHF | 55 | 2 | 0 | 61 | 7 | 125 | Accessory |
| Cj1355 | *ceu* E | Enterochelin uptake periplasmic binding protein | P | MKKSLVFAFFAFF | 55 | 90 | 26 | 255 | 64 | 490 | Core |
| Cj1278c | *trm* B | putative tRNA (guanine-N(7)-)-methyltransferase | J | MPNFKSKKIKEIN | 55 | 90 | 26 | 255 | 64 | 490 | Core |
| Cj1250 | *pur* D | phosphoribosylamine--glycine ligase | F | MKIMILGSGAREY | 55 | 90 | 26 | 255 | 64 | 490 | Core |
| Cj1248 | *gua* A | GMP synthase (glutamine-hydrolyzing) | F | MKKADILVLDFGS | 55 | 90 | 26 | 255 | 64 | 490 | Core |
| Cj1246c | *uvr* C | excinuclease ABC subunit C | L | MTKENLENELKTI | 53 | 84 | 26 | 247 | 63 | 473 | Accessory |
| Cj1240c | - | Hypothetical protein | - | MKKIILLLGLSLVS | 55 | 86 | 21 | 249 | 64 | 475 | Accessory |
| Cj1234 | *gly* S | glycyl-tRNA synthetase beta chain | J | MSELLIEIGTEELP | 55 | 90 | 26 | 255 | 64 | 490 | Core |
| Cj1233 | *ppa* X | putative HAD-superfamily hydrolase | S | MINVFFDMDGTL | 55 | 87 | 25 | 255 | 63 | 485 | Accessory |
| Cj1228c | *htr* A | serine protease (protease DO) | M | MKKIFLSLSLASAL | 55 | 90 | 26 | 255 | 64 | 490 | Core |
| Cj1163c | *czc* D | putative cation transport protein / putative heavy-metal-associated domain protein | P | MYKFLSHEPLANI | 55 | 90 | 26 | 255 | 64 | 490 | Core |
| Cj1162c | *cop* Z | putative cation transport protein / putative heavy-metal-associated domain protein | P | MYKFLSHEPLANI | 55 | 90 | 26 | 255 | 64 | 490 | Core |
| Cj1048c | *dap* E | succinyl-diaminopimelate desuccinylase | E | MNAKEFLIELLKF | 55 | 90 | 26 | 255 | 64 | 490 | Core |
| NCTC13261_01099 | - | dna methylase-type I restriction-modification system | V | MIEEILKDSDYKLD | 30 | 34 | 0 | 46 | 27 | 137 | Accessory |
| NCTC13261_01098 | *rlf* A | RlfA | S | MDNMKEFKIFCD | 29 | 8 | 13 | 2 | 15 | 67 | Accessory |
| NCTC13261_01097 | - | hypothetical protein | - | MYYLLDFSTCKCE | 36 | 35 | 14 | 50 | 34 | 169 | Accessory |
| NCTC13261_01096 | - | Putative uncharacterized protein | V | MNLNQTLQEKYP | 36 | 30 | 0 | 47 | 21 | 134 | Accessory |
| Cj1047c | *thi* S | thiamine biosynthesis protein | H | MIINGEKFEFKELI | 55 | 90 | 14 | 254 | 51 | 464 | Accessory |
| Cj1008c | *aro* B | 3-dehydroquinate synthase | E | MQVEVKLKENAY | 55 | 90 | 26 | 255 | 63 | 489 | Core |
| Cj1002c | *six* A | putative phosphoglycerate/bisphosphoglycerate mutase | T | MKKIYIIRHAKASI | 42 | 88 | 23 | 216 | 32 | 401 | Accessory |
| Cj1001 | *rpo* D | RNA polymerase sigma factor (sigma-70) | K | MNAKTQEAELEE | 55 | 90 | 26 | 255 | 64 | 490 | Core |
| Cj0999c | *yei* H | putative integral membrane protein | S | MKTSFLAHSVAIV | 55 | 90 | 26 | 255 | 64 | 490 | Core |

| | | | | | | | | | | | |
|---|---|---|---|---|---|---|---|---|---|---|---|
| Cj0995c | *hem* B | delta-aminolevulinic acid dehydratase | H | MFKRFRRLRLNE | 55 | 90 | 26 | 255 | 64 | 490 | Core |
| Cj0994c | *arg* F | ornithine carbamoyltransferase | E | MKHFLTLRDFSKE | 55 | 90 | 26 | 255 | 64 | 490 | Core |
| Cj0991c | *glp* C | putative oxidoreductase ferredoxin-type electron transport protein | C | MKFSQISDACVK | 55 | 90 | 26 | 255 | 64 | 490 | Core |
| Cj0967 | - | putative periplasmic protein | - | MIIHGPSKAYASL | 37 | 79 | 26 | 192 | 45 | 379 | Accessory |
| Cj0967 | - | periplasmic protein | - | MREDEVLSFKAR | 37 | 54 | 26 | 60 | 34 | 211 | Accessory |
| Cj0964 | - | putative periplasmic protein | E | MKKIFLIFFLSCFL | 55 | 90 | 26 | 255 | 64 | 490 | Core |
| Cj0959c | *yid* D | hypothetical protein | S | MGIEKTRMFQM | 55 | 90 | 26 | 255 | 63 | 489 | Core |
| Cj0946 | *spr* 7 | putative lipoprotein | M | MKSCLYFTFIVLFL | 55 | 90 | 26 | 255 | 64 | 490 | Core |
| Cj0944c | - | putative periplasmic protein | - | MKKIFLSVFLVLSL | 55 | 90 | 25 | 255 | 62 | 487 | Core |
| Cj0940c | *gln* M | putative glutamine transport system permease | P | MSLFTQKNKNFK | 55 | 90 | 26 | 255 | 64 | 490 | Core |
| Cj0939c | - | hypothetical protein | - | MLIIGHELLKNLD | 53 | 90 | 26 | 255 | 63 | 487 | Core |
| Cj0938c | *aas* | putative 2-acylglycerophosphoethanolamine acyltransferase / acyl-acyl carrier protein synthetase | EGP | MQKKSFLKIYGLI | 45 | 82 | 25 | 215 | 34 | 401 | Accessory |
| Cj0934c | IV02_29000 | putative sodium:amino-acid symporter family protein | P | MRTYFSKIGFVLA | 32 | 81 | 0 | 175 | 15 | 303 | Accessory |
| Cj0933c | *pyc* B | putative pyruvate carboxylase B subunit | C | MAKKFIDVMDTS | 55 | 90 | 26 | 255 | 64 | 490 | Core |
| Cj0932c | *pck* A | phosphoenolpyruvate carboxykinase (ATP) | H | MKKFDNLGLDNI | 55 | 90 | 26 | 255 | 64 | 490 | Core |
| Cj0920c | *glt* J | putative ABC-type amino-acid transporter permease protein | P | MNESVGFVEHLR | 55 | 90 | 26 | 255 | 64 | 490 | Core |
| Cj0919c | *tcy* B_2 | Amino acid ABC transporter, permease protein PEB1 | P | MENVFNAQNIEF | 55 | 90 | 26 | 255 | 64 | 490 | Core |
| Cj0917c | *cst* A | putative integral membrane protein (CstA ) | T | MTQLSTKILWLFV | 52 | 88 | 26 | 245 | 63 | 474 | Accessory |
| Cj0916c | *ybd* D | hypothetical protein | S | MNFKKIKYYYEKA | 55 | 90 | 26 | 254 | 64 | 489 | Core |
| Cj0915 | *yci* A | putative hydrolase | I | MRDMGEPKLKIV | 55 | 90 | 26 | 255 | 64 | 490 | Core |
| Cj0912c | *cys* K | cysteine synthase | E | MKVYEKVSELIGN | 52 | 88 | 26 | 249 | 64 | 479 | Accessory |
| Cj0911 | hyaE | putative periplasmic protein | S | MKKNIILFIVIVAII | 55 | 90 | 26 | 255 | 64 | 490 | Core |
| Cj0909 | VY92_09940 | putative periplasmic protein | S | MKKILLLGALFAA | 55 | 89 | 26 | 252 | 64 | 486 | Core |
| Cj0908 | - | putative periplasmic protein | - | MIRNFFIGMSFLG | 46 | 80 | 24 | 211 | 35 | 396 | Accessory |
| Cj0905c | *alr* | alanine racemase | E | MSLIKIDQKAYEY | 55 | 90 | 26 | 255 | 64 | 490 | Core |
| Cj0903c | *agc* S | putative amino-acid transport protein | E | MKLDIMLDFANK | 55 | 90 | 25 | 254 | 64 | 488 | Core |
| Cj0902 | *art* M | putative glutamine transport ATP-binding protein | E | MIEVKNLQKKYG | 55 | 90 | 26 | 255 | 64 | 490 | Core |
| Cj0886c | *fts* K | putative cell division protein | D | MLAPGMGEWVY | 55 | 90 | 26 | 255 | 64 | 490 | Core |
| Cj0879c | - | putative periplasmic protein | - | MKKCILIFFSLYSLS | 38 | 57 | 4 | 140 | 57 | 296 | Accessory |

| | | | | | | | | | | | |
|---|---|---|---|---|---|---|---|---|---|---|---|
| NCTC13261_00864 | - | Arylsulfotransferase | M | MQPVDKNGKKIV | 33 | 36 | 1 | 74 | 25 | 169 | Accessory |
| NCTC13261_00861 | - | Arylsulfotransferase | M | MRLSKTLCMALLA | 48 | 65 | 25 | 118 | 26 | 282 | Accessory |
| Cj0865 | *dsb* B | disulfide oxidoreductase | C | MKDNCRNFSLSK | 55 | 90 | 26 | 254 | 64 | 489 | Core |
| NCTC13261_00859 | *dsb* A | thiol:disulfide interchange protein DsbA | O | MKFPVKLARSIVV | 55 | 78 | 26 | 255 | 27 | 441 | Accessory |
| Cj0863c | *xer* H | DNA recombinase | L | MKYPLDCEENFEI | 55 | 90 | 26 | 255 | 64 | 490 | Core |
| Cj0821 | *glm* U | UDP-N-acetylglucosamine pyrophosphorylase | M | MKTSILILAAGLGT | 55 | 90 | 26 | 255 | 64 | 490 | Core |
| Cj0794 | - | hypothetical protein | S | MADTIRQYYSKFN | 43 | 81 | 25 | 20 | 44 | 213 | Accessory |
| Cj0814 | - | hypothetical protein | S | MSIFSINDNSNYN | 42 | 18 | 0 | 102 | 18 | 180 | Accessory |
| Cj0812 | *thr* C | threonine synthase Functional classification-Amino acid biosynthesis-Aspartate family protein | E | MKLVESRNVNNV | 55 | 80 | 26 | 255 | 64 | 480 | Accessory |
| Cj0800c | - | putative ATPase | L | MALDEKIIAYTEN | 55 | 90 | 26 | 255 | 64 | 490 | Core |
| Cj0799c | *ruv* A | putative Holliday junction ATP-dependent DNA helicase | L | MVVGIEGIITKKEI | 55 | 90 | 26 | 255 | 64 | 490 | Core |
| Cj0798c | *ddl* | D-alanine--D-alanine ligase | F | MKFAILFGGNSYE | 55 | 90 | 26 | 255 | 64 | 490 | Core |
| Cj0794 | - | hypothetical protein | S | MITQIIILYFHKQK | 48 | 76 | 19 | 174 | 22 | 339 | Accessory |
| Cj0793 | *flg* S | signal transduction histidine kinase | T | MNESILKSLDSNE | 55 | 90 | 26 | 237 | 61 | 469 | Accessory |
| Cj0792 | - | hypothetical protein | S | MLYKRKNIKFKNF | 55 | 90 | 26 | 250 | 62 | 483 | Accessory |
| Cj0791c | *csd* A | putative aminotransferase | E | MQISDLKKELILKK | 55 | 90 | 26 | 255 | 63 | 489 | Core |
| Cj0783 | *nap* B | periplasmic nitrate reductase small subunit (cytochrome C-type protein) | C | MKKKLVLLGSAAY | 55 | 90 | 26 | 255 | 64 | 490 | Core |
| Cj0780 | *nap* A | periplasmic nitrate reductase | C | MNRRDFIKNTAIA | 55 | 90 | 26 | 254 | 64 | 489 | Core |
| Cj0777 | *rep* | putative ATP-dependent DNA helicase | L | MPLSKLNNEQYLA | 55 | 90 | 26 | 255 | 64 | 490 | Core |
| Cj0776c | - | putative periplasmic protein | - | MNLKIFSIIVGILIA | 55 | 90 | 26 | 255 | 64 | 490 | Core |
| Cj0775c | *val* S | valyl-tRNA synthetase | J | MYDKNLEKEYYQ | 55 | 90 | 26 | 255 | 64 | 490 | Core |
| Cj0772c | *met* Q | putative NLPA family lipoprotein | P | MKIKSLFIASILTLS | 55 | 90 | 26 | 255 | 64 | 490 | Core |
| Cj0771c | *met* Q | putative NLPA family lipoprotein | M | MNLFKIIILACILNI | 55 | 90 | 26 | 255 | 64 | 490 | Core |
| Cj0763c | *cys* E | serine acetyltransferase | E | MNFWGIIKEDFSG | 55 | 90 | 26 | 255 | 64 | 490 | Core |
| Cj0762c | *asp* C | aspartate aminotransferase | E | MLTKRSQVLEESI | 55 | 90 | 26 | 255 | 64 | 490 | Core |
| Cj0757 | *hrc* A | putative heat shock regulator | K | MKSRDKKDLILES | 36 | 74 | 0 | 195 | 15 | 320 | Accessory |
| Cj0755 | *cfr* A | ferric receptor CfrA | P | MKKLCLSVCAIGL | 26 | 55 | 0 | 144 | 1 | 226 | Accessory |
| Cj0753c | *ton* B | energy transducer TonB | M | MKAFISNHKNQS | 36 | 73 | 0 | 192 | 12 | 313 | Accessory |
| Cj0752 | - | ISCco1, transposase orfB | - | MRYHRDLPNTYK | 29 | 65 | 0 | 176 | 12 | 282 | Accessory |
| Cj0718 | *dna* E | DNA polymerase III, alpha chain | L | MSQFTHLHLHTE | 55 | 90 | 26 | 255 | 64 | 490 | Core |
| Cj0717 | *spx* A | putative ArsC family protein | P | MKLYGIKNCNSVI | 55 | 90 | 26 | 255 | 64 | 490 | Core |

| | | | | | | | | | | | |
|---|---|---|---|---|---|---|---|---|---|---|---|
| Cj0716 | *aro* F | putative phospho-2-dehydro-3-deoxyheptonate aldolase | E | MWTKNSWKNYI | 55 | 90 | 26 | 255 | 64 | 490 | Core |
| Cj0715 | *ura* H | transthyretin-like periplasmic protein | S | MFSIKKTLLILASV | 55 | 90 | 26 | 255 | 64 | 490 | Core |
| Cj0714 | *rpl* S | 50S ribosomal protein L19 | J | MKNKYIEQFEAK | 55 | 90 | 26 | 255 | 64 | 490 | Core |
| Cj0713 | *trm* D | tRNA (guanine-N1)-methyltransferase | J | MKFSFVSLFPNLN | 55 | 90 | 26 | 255 | 64 | 490 | Core |
| Cj0712 | *rim* M | putative 16S rRNA processing protein | J | MSEKDFVQVAKL | 55 | 89 | 26 | 254 | 63 | 487 | Core |
| Cj0710 | *rps* P | 30S ribosomal protein S16 | J | MTVIRLTRMGRT | 55 | 90 | 26 | 255 | 64 | 490 | Core |
| Cj0709 | *ffh* | signal recognition particle protein | U | MFELVSESFKSAI | 55 | 90 | 26 | 255 | 64 | 490 | Core |
| Cj0535 | *oor* D | OORD subunit of 2-oxoglutarate:acceptor oxidoreductase | C | MSMIAPKDTPVV | 55 | 90 | 26 | 255 | 64 | 490 | Core |
| Cj0534 | *suc* D | succinyl-coA synthetase alpha chain | C | MSILVNKNTKVIV | 55 | 90 | 26 | 255 | 64 | 490 | Core |
| Cj0532 | *mdh* | malate dehydrogenase | C | MKITVIGAGNVG | 55 | 90 | 26 | 255 | 64 | 490 | Core |
| Cj0495 | - | putative methyltransferase domain protein | S | MSDLITLAQLSQ | 55 | 90 | 26 | 255 | 64 | 490 | Core |
| Cj0494 | - | putative exporting protein | - | MIKIAFFITFVISFL | 46 | 53 | 25 | 72 | 25 | 221 | Accessory |
| Cj0493 | *fus* A | elongation factor G | J | MSRNTPLKKVRN | 55 | 90 | 26 | 255 | 64 | 490 | Core |
| Cj0492 | *rps* G | 30S ribosomal protein S7 | J | MRRRKAPVREVL | 55 | 90 | 26 | 255 | 64 | 490 | Core |
| Cj0491 | *rps* L | 30S ribosomal protein S12 | J | MPTINQLVRKERI | 55 | 90 | 26 | 255 | 64 | 490 | Core |
| Cj0479 | *rpo* C | DNA-directed RNA polymerase beta' chain | K | MSKFKVIEIKEDA | 55 | 90 | 26 | 255 | 64 | 490 | Core |
| Cj0478 | *rpo* B | DNA-directed RNA polymerase beta chain | K | MLDNKLGNRLRV | 55 | 90 | 26 | 255 | 64 | 490 | Core |
| Cj0464 | *rec* G | ATP-dependent DNA helicase | L | MKIKESDFEFFKK | 55 | 89 | 26 | 250 | 63 | 483 | Accessory |
| Cj0463 | *ymx* G | zinc protease-like protein | S | MQYLESRGVKIPI | 55 | 90 | 26 | 255 | 64 | 490 | Core |
| Cj0462 | *mqn* C | putative radical SAM domain protein | H | MKRLDKKEALDLI | 55 | 90 | 26 | 255 | 64 | 490 | Core |
| Cj0461c | *bac* E | putative MFS (Major Facilitator Superfamily) transport protein | EGP | MNYIELLKNNKNI | 55 | 90 | 26 | 255 | 64 | 490 | Core |
| Cj0460 | *nus* A | transcription termination factor | K | MEKIADIIESIANE | 55 | 90 | 26 | 255 | 64 | 490 | Core |
| Cj0459c | - | hypothetical protein | - | MELKLARTLINEK | 55 | 90 | 26 | 255 | 64 | 490 | Core |
| Cj0458c | *mia* B | putative tRNA 2-methylthioadenosine synthase | J | MSAKKLFIQTLGC | 55 | 90 | 25 | 255 | 64 | 489 | Core |
| Cj0456c | - | hypothetical protein | - | MLFLLKKIFPQLFI | 55 | 90 | 26 | 255 | 64 | 490 | Core |
| Cj0455c | *pil* N | hypothetical protein | NU | MTYSFIQPRKKPI | 55 | 90 | 26 | 255 | 64 | 490 | Core |
| Cj0454c | - | putative membrane protein | - | MKDKSLEEIDLLK | 55 | 90 | 26 | 255 | 64 | 490 | Core |
| Cj0453 | *thi* C | thiamin biosynthesis protein ThiC | H | MKTQMNYAKEG | 3 | 0 | 0 | 3 | 0 | 6 | Accessory |

| | | | | | | | | | | | |
|---|---|---|---|---|---|---|---|---|---|---|---|
| Cj0451 | *rpe* | Ribulose-phosphate 3-epimerase | G | MYVAPSLLSANFL | 55 | 90 | 26 | 255 | 64 | 490 | Core |
| Cj0450c | *rpm* B | 50S ribosomal protein L28 | J | MARVCQITGKGP | 55 | 90 | 26 | 255 | 64 | 490 | Core |
| Cj0449c | *ydc* H | hypothetical protein | S | MLHEYRELMSEL | 55 | 90 | 26 | 255 | 64 | 490 | Core |
| Cj0448c | - | putative MCP-type signal transduction protein | NT | MFGSKINHSDLQ | 21 | 24 | 26 | 81 | 58 | 210 | Accessory |
| Cj0444 | *cirA* _3 | TonB-dependent receptor | P | MVALYGENEYFIT | 35 | 66 | 0 | 174 | 6 | 281 | Accessory |
| Cj0444 | *cirA* _3 | Ferric receptor CfrA | P | MIEDTLSISARLKY | 35 | 53 | 0 | 140 | 0 | 228 | Accessory |
| Cj0437 | *frd* A | succinate dehydrogenase flavoprotein subunit | C | MGEFSRRDFIKTA | 55 | 78 | 26 | 249 | 62 | 470 | Accessory |
| Cj0435 | - | 3-oxoacyl-[acyl-carrier protein] reductase | IQ | MKFSGKNVLITGA | 55 | 90 | 26 | 255 | 64 | 490 | Core |
| Cj0434 | *gpm* I | 2,3-bisphosphoglycerate-independent phosphoglycerate mutase | G | MKQKCVLIITDGI | 55 | 90 | 26 | 255 | 64 | 490 | Core |
| Cj0432c | *mur* D | UDP-N-acetylmuramoylalanine--D-glutamate ligase | M | MKISLFGYGKTTR | 55 | 90 | 26 | 255 | 64 | 490 | Core |
| NCTC13261_01757 | *rpl* V | ribosomal protein L22 | J | MSKALIKFIRLSPT | 55 | 90 | 26 | 255 | 64 | 490 | Core |
| Cj1701c | *rps* C | 30S ribosomal protein S3 | J | MGQKVNPIGLRL | 55 | 90 | 26 | 255 | 64 | 490 | Core |
| Cj1688c | *sec* Y | preprotein translocase subunit | U | MNRALTNKILITLA | 55 | 90 | 26 | 255 | 64 | 490 | Core |
| Cj1687 | - | putative efflux protein | EGP | MVVWTISGISGFI | 37 | 77 | 4 | 196 | 50 | 364 | Accessory |
| Cj1687 | - | Major Facilitator Superfamily protein | EGP | MNGKTYKFHPND | 54 | 14 | 25 | 82 | 19 | 194 | Accessory |
| Cj1686c | *top* A | DNA topoisomerase I | L | MKKNLIIVESPAK | 55 | 90 | 26 | 255 | 64 | 490 | Core |
| Cj1685c | *bio* B | putative biotin synthase | H | MQIMLCAISNIAS | 55 | 90 | 26 | 255 | 64 | 490 | Core |
| Cj1681c | *cys* Q | CysQ protein | P | MLNLDKFLEIAIN | 55 | 90 | 26 | 255 | 64 | 490 | Core |
| Cj1673c | *rec* A | recA protein | L | MDDNKRKSLDAA | 55 | 90 | 26 | 255 | 64 | 490 | Core |
| Cj1672c | *eno* | enolase | G | MLVIEDVRAYEVL | 55 | 90 | 26 | 255 | 64 | 490 | Core |
| Cj1670c | *cgp* A | putative periplasmic protein | S | MKTRILAIFFIFTSI | 55 | 90 | 26 | 255 | 64 | 490 | Core |
| Cj1669c | *lig* | putative ATP-dependent DNA ligase | L | MRFVFLICCACLV | 55 | 90 | 26 | 255 | 64 | 490 | Core |
| NCTC13261_01720 | - | integrase | L | MLIHSGRRRDEVI | 35 | 20 | 0 | 1 | 4 | 60 | Accessory |
| NCTC13261_01719 | *Hic* A | HicA | - | MLGLEFIFSKQRL | 35 | 20 | 0 | 0 | 4 | 59 | Accessory |
| NCTC13261_01718 | - | hypothetical protein | N | MHLKYFKIIIYNTY | 35 | 20 | 0 | 0 | 4 | 59 | Accessory |
| NCTC13261_01717 | *Hic* B | HicB | S | MKKDINYYLNLPY | 34 | 14 | 0 | 1 | 4 | 53 | Accessory |
| NCTC13261_01716 | - | putative protein | - | MNLILENPSEKLIK | 35 | 0 | 0 | 0 | 0 | 35 | Accessory |
| NCTC13261_01715 | *yaf* Q | putative RelE/StbE family addiction module toxin | - | MIKYTLKKSKEFKI | 35 | 0 | 0 | 0 | 0 | 35 | Accessory |
| NCTC13261_01714 | - | helix-turn-helix domain-containing protein | - | MQDTNKDFLTIQ | 35 | 19 | 0 | 1 | 4 | 59 | Accessory |
| NCTC13261_01713 | - | hypothetical protein | - | MNDRQALEYLKD | 35 | 0 | 0 | 1 | 0 | 36 | Accessory |
| NCTC13261_01712 | - | hypothetical protein | - | MSISLFNDEIKAYI | 34 | 7 | 0 | 1 | 4 | 46 | Accessory |

| | | | | | | | | | | | |
|---|---|---|---|---|---|---|---|---|---|---|---|
| NCTC13261_01711 | *dna* G | DnaB-like protein helicase-like protein | L | MKDYSKEINDLKK | 30 | 19 | 0 | 0 | 4 | 53 | Accessory |
| NCTC13261_01710 | - | hypothetical protein | - | MTRLEFDEKIKQL | 35 | 0 | 0 | 0 | 0 | 35 | Accessory |
| NCTC13261_01709 | - | acyl carrier protein | K | MTKEKFNQLLKQ | 34 | 0 | 0 | 0 | 0 | 34 | Accessory |
| NCTC13261_01708 | - | hypothetical protein | - | MEKVDYCRMSCL | 35 | 0 | 0 | 0 | 0 | 35 | Accessory |
| NCTC13261_01707 | - | hypothetical protein | - | MQTITIKAKEQEI | 35 | 0 | 0 | 0 | 0 | 35 | Accessory |
| NCTC13261_01706 | - | RelE/ParE family plasmid stabilization system protein | S | MQILESDLFTNEL | 35 | 20 | 0 | 0 | 4 | 59 | Accessory |
| NCTC13261_01705 | - | putative periplasmic protein | - | MKKLGLALAVLV | 35 | 38 | 0 | 193 | 43 | 309 | Accessory |
| Cj0313 | CP_0860 | putative integral membrane protein | S | MWIFFRFISGIYLI | 55 | 90 | 26 | 255 | 64 | 490 | Core |
| Cj0314 | *lys* A | diaminopimelate decarboxylase | E | MDYKQLKQEFNT | 55 | 90 | 26 | 255 | 64 | 490 | Core |
| Cj0316 | *phe* A | chorismate mutase/prephenate dehydratase | E | MPNLEEFRNKIDA | 55 | 90 | 26 | 255 | 64 | 490 | Core |
| Cj0317 | *his* C | histidinol-phosphate aminotransferase | E | MKFNEFLNHLSN | 55 | 90 | 26 | 255 | 64 | 490 | Core |
| Cj0318 | *fli* F | flagellar M-ring protein | N | MDFKNMLHQIG | 55 | 90 | 26 | 255 | 64 | 490 | Core |
| Cj0360 | *glm* M | phosphoglucosamine mutase | G | MKLFGTDGVRGK | 55 | 90 | 26 | 255 | 64 | 490 | Core |
| Cj0392c | *pyk* | pyruvate kinase I | G | MLKKTKIVATVGF | 55 | 90 | 25 | 255 | 64 | 489 | Core |
| Cj0393c | *mqo* | putative malate:quinone oxidoreductase | C | MSQQEFDVLVIG | 55 | 90 | 26 | 255 | 64 | 490 | Core |
| Cj0394c | *coa* X | putative transcriptional activator | F | MLLCDIGNSNAN | 55 | 90 | 26 | 255 | 64 | 490 | Core |
| Cj0396c | *pgb* B | putative lipoprotein | - | MKKIFLTLFCLIFL | 55 | 90 | 26 | 255 | 63 | 489 | Core |
| Cj0397c | - | hypothetical protein | - | MGLKDNLKAVKN | 55 | 89 | 26 | 252 | 64 | 486 | Core |
| Cj0398 | *gat* C | Glu-tRNAGln amidotransferase subunit C | J | MQIDEKLLSKLEK | 55 | 90 | 26 | 255 | 64 | 490 | Core |
| Cj0399 | *cvp* A | cvpA family protein | S | MNFYWFDAFILG | 55 | 90 | 26 | 255 | 64 | 490 | Core |
| Cj0401 | *lys* S | lysyl-tRNA synthetase | J | MFDNILEQQRIEK | 55 | 90 | 26 | 255 | 64 | 490 | Core |
| Cj0404 | - | putative transmembrane protein | S | MENQKNEFDDIII | 55 | 90 | 26 | 255 | 64 | 490 | Core |
| Cj0405 | *aro* E | shikimate 5-dehydrogenase | E | MKFLAVIGDPISH | 55 | 90 | 26 | 255 | 64 | 490 | Core |
| Cj0406c | - | lipoprotein, putative | - | MKKYILLASSMLII | 55 | 90 | 25 | 255 | 59 | 484 | Accessory |
| Cj0418c | Cj0418c | hypothetical protein | M | MQISSSYNSYDYY | 54 | 34 | 26 | 93 | 41 | 248 | Accessory |
| Cj0420 | Cj0420 | putative periplasmic protein | S | MKKVVLSSLVAVS | 55 | 90 | 26 | 255 | 64 | 490 | Core |
| Cj0421c | Cj0421c | putative integral membrane protein | - | MFLLDFLISLSFIFE | 55 | 90 | 26 | 255 | 64 | 490 | Core |
| Cj0422c | - | putative H-T-H containing protein | - | MTKKSKRDMAYE | 36 | 85 | 26 | 255 | 49 | 451 | Accessory |
| NCTC13261_00426 | - | integral membrane protein | - | MDFNSVLFSIGDI | 35 | 0 | 23 | 44 | 14 | 116 | Accessory |
| NCTC13261_00427 | - | lipoprotein, putative | - | MKCFNAKFMFIG | 35 | 0 | 23 | 44 | 15 | 117 | Accessory |
| NCTC13261_00428 | - | Integral membrane protein | - | MDYNVFLLNIQD | 26 | 0 | 23 | 30 | 11 | 90 | Accessory |

| | | | | | | | | | | | |
|---|---|---|---|---|---|---|---|---|---|---|---|
| Cj0426 | *ybi* T | putative ABC transporter ATP-binding protein | S | MVEVKNLTMRFA | 55 | 90 | 26 | 255 | 64 | 490 | Core |
| Cj0427 | - | hypothetical protein | - | MAKFRIQYSAGF | 55 | 90 | 26 | 255 | 64 | 490 | Core |
| Cj0428 | - | hypothetical protein | - | MQVNYRTISSYEY | 55 | 90 | 26 | 255 | 64 | 490 | Core |
| Cj0429c | *yig* Z | hypothetical protein | S | MQTIDQIFQTQID | 55 | 90 | 26 | 255 | 64 | 490 | Core |
| Cj0430 | - | putative protein, PMT family | M | MEKIKNYKLIIILLS | 55 | 90 | 26 | 255 | 64 | 490 | Core |
| Cj0431 | - | hypothetical protein | NU | MIKAFSLLEFVFIIL | 55 | 90 | 26 | 255 | 64 | 490 | Core |
| Cj0129c | *bam* A | outer membrane protein assembly complex, YaeT protein | M | MKKHLISICALVA | 55 | 90 | 26 | 255 | 64 | 490 | Core |
| Cj0127c | *acc* D | acetyl-coenzyme A carboxylase carboxyl transferase subunit beta | I | MNFADIFSKIRRC | 55 | 90 | 26 | 255 | 64 | 490 | Core |
| Cj0105 | *atp* A | ATP synthase F1 sector alpha subunit | C | MKFKADEISSIIKE | 55 | 90 | 26 | 255 | 64 | 490 | Core |
| Cj0100 | *par* A | parA family protein | D | MSEIITIANQKGG | 55 | 90 | 26 | 255 | 64 | 490 | Core |
| Cj0099 | *bir* A | putative biotin--[acetyl-CoA-carboxylase] synthetase | H | MEKGLKVKIVCVE | 55 | 90 | 26 | 255 | 64 | 490 | Core |
| Cj0098 | *fmt* | methionyl-tRNA formyltransferase | J | MKKIIFMGTPSYA | 55 | 90 | 26 | 255 | 64 | 490 | Core |
| Cj0096 | *obg* | GTP-binding protein, GTP1/Obg family | S | MFIDSVKITLASG | 55 | 90 | 26 | 255 | 64 | 490 | Core |
| Cj0093 | - | putative periplasmic protein | M | MKIIKILFLGLFLSL | 55 | 88 | 26 | 214 | 42 | 425 | Accessory |
| Cj0089 | - | putative lipoprotein | S | MKIKVGLIFSGIVC | 55 | 90 | 26 | 255 | 64 | 490 | Core |
| Cj0088 | *dcu* A | anaerobic C4-dicarboxylate transporter | U | MDIMIILQVIVLLC | 55 | 88 | 26 | 221 | 62 | 452 | Accessory |
| Cj0087 | *asp* A | aspartate ammonia-lyase | E | MGTRKEHDFIGE | 55 | 90 | 26 | 255 | 64 | 490 | Core |
| Cj0086c | *ung* | uracil-DNA glycosylase | L | MEEITINIDKIKINI | 55 | 90 | 26 | 255 | 64 | 490 | Core |
| Cj0085c | *rac* D | putative amino acid recemase | M | MKTIGIIGGMSFE | 36 | 90 | 26 | 187 | 28 | 367 | Accessory |
| Cj0082 | *cyd* B | cytochrome bd oxidase subunit II | C | MFFGLELEGLQIY | 55 | 90 | 26 | 255 | 64 | 490 | Core |
| Cj0076c | *lld* P | L-lactate permease | C | MEQILTWQQIYD | 54 | 87 | 1 | 254 | 64 | 460 | Accessory |
| Cj0197c | *dap* B | dihydrodipicolinate reductase | E | EAAKNATLAHGLY | 55 | 90 | 26 | 255 | 64 | 490 | Core |
| Cj0196c | *pur* F | amidophosphoribosyltransferase | F | RKFSQNYDCVLVI | 55 | 90 | 26 | 255 | 64 | 490 | Core |
| Cj0195 | *fli* I | flagellum-specific ATP synthase | NU | QNTPLILDANCFL | 55 | 90 | 26 | 255 | 64 | 490 | Core |
| Cj0193c | *tig* | trigger factor (peptidyl-prolyl cis /trans isomerase, chaperone) | D | SAGTLAGLGALNI | 55 | 90 | 26 | 255 | 64 | 490 | Core |
| Cj0192c | *clp* P | ATP-dependent Clp protease proteolytic subunit | O | EDFAKEFVGRIKIA | 55 | 90 | 26 | 255 | 64 | 490 | Core |
| Cj0191c | *def* | polypeptide deformylase | J | CILGTGSNRCLDE | 55 | 90 | 26 | 255 | 64 | 490 | Core |
| Cj0189c | - | hypothetical protein | S | GGNNASDGLVAL | 55 | 90 | 26 | 255 | 64 | 490 | Core |
| Cj0188c | *nnr* D | putative kinase | - | MKAIIDNIKILKQN | 55 | 90 | 26 | 255 | 64 | 490 | Core |

| | | | | | | | | | | | |
|---|---|---|---|---|---|---|---|---|---|---|---|
| Cj0184c | - | Ser/Thr protein phosphatase family protein | T | MRILLVLRGNYYA | 55 | 90 | 26 | 255 | 64 | 490 | Core |
| Cj0183 | *cor* C | putative integral membrane protein with haemolysin domain protein | S | MDPSQVLDLNQT | 55 | 90 | 26 | 255 | 64 | 490 | Core |
| Cj0182 | *sbm* A | putative transmembrane transport protein | I | MFSSFFKSKKWA | 55 | 76 | 26 | 255 | 29 | 441 | Accessory |
| Cj0179 | *exb* B | biopolymer transport protein | U | MLKKMFLFCFLLL | 36 | 64 | 0 | 194 | 14 | 308 | Accessory |
| Cj0178 | - | putative TonB-denpendent outer membrane receptor | P | MQNLFDKNYMD | 34 | 9 | 0 | 58 | 6 | 107 | Accessory |
| Cj0178 | - | putative TonB-denpendent outer membrane receptor | P | MKKLSLFCAVGLC | 23 | 3 | 0 | 24 | 3 | 53 | Accessory |

[a]Locus tags for accessory genes based on C. jejuni reference strain NCTC13261 genome LR134500.1 (NCBI accession) while locus tags for allelic variants of the core genome refer to C. jejuni strain NCTC11168 (NCBI accession: AL111168.1)

[b]clusters of orthologous groups (http://clovr.org/docs/clusters-of-orthologous-groups-cogs/);

[c]number of genomes assigned to a particular lifestyle carrying the gene or allelic variant (pig, cattle, chicken, host generalists, others)

[d]indicates that a gene belongs to the accessory (A) or the core (C) genome content of C. jejuni
