## Supplementary material for "Genome-wide insights into population structure and host specificity of *Campylobacter jejuni*": Table S4

Table S4: Overview of chicken associated accessory genes and core gene variants identified by k-mer mapping.

| Reference Gene Nane[a] | Alias | Predicted Function | COG Family[b] | AA | # Cattle[c] | # Chicken[c] | # Pig[c] | # Generalist[c] | # Others[c] | Total | Core/accessory[d] |
|---|---|---|---|---|---|---|---|---|---|---|---|
| Cj1033 | *cme* F | integral membrane component of efflux system (multidrug efflux system CmeDEF) | V | MFKLAINRPITVLMFFLALN | 54 | 85 | 26 | 220 | 60 | 445 | Accessory |
| NCTC13265_01618 | *tra* G | conjugal transfer protein TraG | U | MNEIMQAVKGITTETGYIV | 1 | 59 | 1 | 1 | 13 | 75 | Accessory |
| NCTC13265_01619 | - | TraG-like protein | - | MNEALSDKSSFSETDKKAL | 0 | 16 | 0 | 0 | 0 | 16 | Accessory |
| NCTC13265_01620 | - | TraG-like protein | - | MAENYIAENNLGETREQQ | 0 | 17 | 0 | 0 | 0 | 17 | Accessory |
| NCTC13265_01623 | - | putative protein | - | MKTTEYNTQQKNNKEYLA | 0 | 35 | 0 | 0 | 5 | 40 | Accessory |
| NCTC13265_01624 | - | death-on-curing family protein | - | MKYIELSEAIAIHEKIIEKTG | 0 | 35 | 0 | 0 | 5 | 40 | Accessory |
| LR59_01905 | *doc* | death-on-curing family protein | S | MKYIELSEAIAIHEKIIEKTG | 0 | 30 | 0 | 0 | 5 | 35 | Accessory |
| NCTC13265_01627 | - | putative protein | - | MAIKIEISGFDKDFLPVFES | 0 | 66 | 3 | 1 | 7 | 77 | Accessory |
| NCTC13265_01633 | - | putative protein | - | MIIVNDKQLKTLIVKKEHIL | 0 | 68 | 3 | 0 | 0 | 71 | Accessory |
| Cj0976 | *cmo* B | putative methyltransferase | J | MQENLLEKQFLNHPLYAKI | 20 | 63 | 24 | 109 | 37 | 253 | Accessory |
| NCTC13265_01672 | - | putative periplasmic protein | U | MQTTDEDPNINEDDKQAS | 19 | 20 | 0 | 21 | 14 | 74 | Accessory |
| Cj0933c | *pyc* B | putative pyruvate carboxylase B subunit | C | MAKKFIDVMDTSFRDGFQ | 55 | 90 | 26 | 255 | 64 | 490 | Core |
| Cj0912c | *cys* K | cysteine synthase | E | MKVYEKVSELIGNTPIIHLK | 52 | 88 | 26 | 249 | 64 | 479 | Accessory |
| Cj0861c | *trp* G | glutamine amidotransferase | EH | MKKILFIDNYDSFSYTIIYYL | 55 | 90 | 26 | 242 | 57 | 470 | Accessory |
| Cj0849c | - | FIG00469420: hypothetical protein | - | MINTQLASQIANTQKNDL | 0 | 65 | 26 | 14 | 9 | 114 | Accessory |
| Cj0780 | *nap* A | periplasmic nitrate reductase | C | MNRRDFIKNTAIASAASVA | 55 | 90 | 26 | 254 | 64 | 489 | Core |
| Cj0652 | *pbp* C | penicillin-binding protein 2 | M | MRMRLVVGFILLFFIFLLSR | 55 | 90 | 26 | 255 | 64 | 490 | Core |
| Cj0651 | - | putative integral membrane protein | - | MRRNLSAYKAKFNGAYVF | 55 | 90 | 26 | 255 | 64 | 490 | Core |
| Cj0577c | *que* A | S-adenosylmethionine:tRNA ribosyltransferase-isomerase | F | MNKDLLLSSYDYTLANELIA | 55 | 90 | 26 | 255 | 64 | 490 | Core |
| Cj0528c | *flg* B | flagellar basal-body rod protein | N | MINPFKSKELVTGALAGRN | 55 | 90 | 26 | 255 | 64 | 490 | Core |
| Cj0509c | *clp* B | ATP-dependent chaperone protein ClpB | O | MANIQDFLTDNMLSNLES | 55 | 90 | 26 | 255 | 64 | 490 | Core |
| Cj0508 | *pbp* A | penicillin-binding protein | M | MKILKYIFSFFTLLFIAGFIYV | 55 | 90 | 26 | 255 | 64 | 490 | Core |
| Cj0507 | *maf* | Maf-like protein | D | MLILASSSISRANLLKTAKID | 55 | 90 | 26 | 255 | 64 | 490 | Core |
| Cj0497 | - | ATP-dependent nuclease subunit B | S | MYRYLLFVLAAFFLAACGS | 55 | 90 | 26 | 255 | 64 | 490 | Core |
| Cj0496 | - | hypothetical protein | S | MIFDKNFSYAFDENACEK | 55 | 90 | 26 | 255 | 64 | 490 | Core |
| Cj0495 | - | putative methyltransferase domain protein | S | MSDLITLAQLSQGYRYNSD | 55 | 90 | 26 | 255 | 64 | 490 | Core |
| Cj0493 | *fus* A | elongation factor G | J | MSRSTPLKKVRNIGIAAHID | 55 | 90 | 26 | 255 | 64 | 490 | Core |
| Cj0492 | *rps* G | 30S ribosomal protein S7 | J | MRRRKAPVREVLPDPIYGN | 55 | 90 | 26 | 255 | 64 | 490 | Core |
| Cj0479 | *rpo* C | DNA-directed RNA polymerase beta' chain | K | MSKFKVIEIKEDARPRDFEA | 55 | 90 | 26 | 255 | 64 | 490 | Core |
| Cj0478 | *rpo* B | DNA-directed RNA polymerase beta chain | K | MLDNKLGNRLRVDFSNISK | 55 | 90 | 26 | 255 | 64 | 490 | Core |
| Cj0444 | *cir* A_3 | TonB-dependent receptor, putative, degenerate | P | MVYHLNDNIALKGGVSKG | 35 | 53 | 0 | 140 | 0 | 228 | Accessory |

| | | | | | | | | | | | |
|---|---|---|---|---|---|---|---|---|---|---|---|
| Cj1623 | - | putative membrane protein | - | MAFMNFSGFFYARNDLRL | 55 | 90 | 26 | 255 | 64 | 490 | Core |
| Cj1633 | *til* S | thiamine biosynthesis protein:ExsB | D | MKALALFSGGLDSMLAMK | 55 | 90 | 26 | 255 | 64 | 490 | Core |
| Cj0391c | - | hypothetical protein | - | MQVNTFSNIASMAQTQVS | 55 | 90 | 25 | 255 | 64 | 489 | Core |
| NCTC13265_00510 | *bio* C | hypothetical protein | S | MKISNLIQNENSQELILVFG | 14 | 31 | 0 | 0 | 0 | 45 | Accessory |
| Cj0131 | - | putative peptidase M23 family protein | M | MAKRKGKTYLSVLILVLIAIL | 55 | 90 | 26 | 255 | 64 | 490 | Core |
| Cj0662c | *hsl* U | ATP-dependent Hsl protease ATP-binding subunit | O | MNLTPKEIVKFLDDYVIGQ | 55 | 90 | 26 | 255 | 64 | 490 | Core |
| Cj0718 | *dna* E | DNA polymerase III, alpha chain | L | MSQFTHLHLHTEYSLLDGA | 55 | 90 | 26 | 255 | 64 | 490 | Core |
| Cj1481c | *add* A | putative helicase | - | MSQFEPFLALEASAGSGKT | 55 | 90 | 26 | 255 | 64 | 490 | Core |
| Cj1478c | *opr* F | outer membrane fibronectin-binding protein | M | MKKIFLCLGLASVLFGADN | 55 | 90 | 26 | 255 | 64 | 490 | Core |
| Cj1477c | *ppa* X | putative hydrolase | S | MNKTILFDLDGTLIDSTDAI | 55 | 90 | 26 | 255 | 64 | 490 | Core |
| NCTC13265_01596 | - | hypothetical protein | I | MQEKINELKDYAELAQASY | 0 | 20 | 0 | 0 | 0 | 20 | Accessory |

[a]Locus tags for accessory genes based on C. jejuni reference strain NCTC13265 genome LR134498.1 (NCBI accession), while locus tags for allelic variants of the core genome refer to C. jejuni strain NCTC11168 (NCBI accession: AL111168.1)

[b]clusters of orthologous groups (http://clovr.org/docs/clusters-of-orthologous-groups-cogs/);

[c]number of genomes assigned to a particular lifestyle carrying the gene or allelic variant (pig, cattle, chicken, host generalists, others)

[d]indicates that a gene belongs to the accessory (A) or the core (C) genome content of C. jejuni
