## Supplementary material for "Genome-wide insights into population structure and host specificity of *Campylobacter jejuni*": Table S5

Table S5: Overview of pig associated accessory genes and core gene variants identified by k-mer mapping.

| Reference Gene Nane[a] | Alias | Predicted Function | COG Family[b] | AA | # Cattle[c] | # Chicken[c] | # Pig[c] | # Generalist[c] | # Others[c] | Total | Core/Accessory[d] |
|---|---|---|---|---|---|---|---|---|---|---|---|
| Cj1019c | *liv* J | branched-chain amino-acid ABC transport system periplasmic binding protein | E | MKKLTLTLSVLTMVNCL | 55 | 90 | 26 | 255 | 64 | 490 | Core |
| Cj1033 | *cme* F | integral membrane component of efflux system (multidrug efflux system CmeDEF) | V | MFKLAINRPITVLMFFLA | 54 | 85 | 26 | 220 | 60 | 445 | Accessory |
| Cj1034c | - | putative DnaJ-like protein | O | MQIVQTLETINVNTDDI | 55 | 90 | 26 | 255 | 64 | 490 | Core |
| Cj1037c | *pyc* A | acetyl-CoA carboxylase, biotin carboxylase | I | MNQIHKILIANRAEIAV | 55 | 90 | 26 | 255 | 64 | 490 | Core |
| Cj1039 | *mur* G | putative undecaprenyldiphospho-muramoylpentapeptide b-N-acetylglucosaminyltransferase | M | MTIALTGGGTGGHLAIV | 53 | 90 | 26 | 255 | 64 | 488 | Core |
| Cj1040c | *yea* N | membrane protein, putative | P | MLFVFDSKTSIIISAFIM | 0 | 0 | 25 | 0 | 10 | 35 | Accessory |
| Cj1040c | *yea* N | putative transmembrane transport protein | P | MVDFFVLRKKALKKINL | 1 | 0 | 26 | 14 | 0 | 41 | Accessory |
| Cj1040c | *yea* N | Putative transmembrane transport protein | P | MIVAFNLRAPITAIGPM | 0 | 0 | 26 | 0 | 4 | 30 | Accessory |
| Cj1043c | *ten* I | putative thiamine-phosphate pyrophosphorylase | H | MWDKKIIAISDRKCVEI | 55 | 90 | 26 | 255 | 64 | 490 | Core |
| Cj1044c | *thi* H | thiazole biosynthesis protein ThiH | C | MQDYMQYLPHMQEIK | 55 | 90 | 26 | 255 | 64 | 490 | Core |
| A6J90_06670 | Z012_06150 | Type II restriction endonuclease | L | MNKVELGSNTAKNGFK | 0 | 0 | 26 | 0 | 1 | 27 | Accessory |
| A6J90_06675 | *dcm* | cytosine-specific methyltransferase NlaX | H | MKFIDFCSGIGGGRLGL | 0 | 0 | 26 | 0 | 1 | 27 | Accessory |
| Cj1045c | *thi* G | Thiazole biosynthesis protein ThiG | H | MQENLKNDKLKIGKYD | 55 | 90 | 26 | 255 | 64 | 490 | Core |
| Cj1052c | *mut* S2 | putative mismatch repair protein | L | MNDTKEELISKLDLNSYL | 55 | 77 | 26 | 255 | 62 | 475 | Accessory |
| Cj1097 | *sst* T | putative transmembrane transport protein | E | MFSKIIQSYAKGNLIVQI | 49 | 88 | 25 | 236 | 48 | 446 | Accessory |
| Cj1124c | *pgl* A | GalNAc transferase | M | MRIGFLSHAGASIYHFR | 55 | 86 | 26 | 253 | 64 | 484 | Accessory |
| Cj1133 | *waa* C | heptosyltransferase I | M | MKIAIVRLSALGDIIQSA | 55 | 90 | 26 | 255 | 64 | 490 | Core |
| Cj1210 | *yoh* D | putative integral membrane protein | S | MQDMIDTLIKYGYIVLF | 55 | 90 | 26 | 252 | 64 | 487 | Core |
| Cj1211 | *com* EC | putative competence family protein | S | MSLWNSFFYSFKEFHYL | 54 | 87 | 25 | 231 | 60 | 457 | Accessory |
| Cj1213c | *glc* D | putative glycolate oxidase subunit D | C | MKKEFEQYFKRFLGEEN | 55 | 90 | 26 | 255 | 62 | 488 | Core |
| Cj1218c | *rib* E | putative riboflavin synthase alpha chain | H | MFNGLIREIAEVQSYQN | 55 | 90 | 26 | 255 | 64 | 490 | Core |
| Cj1219c | *ytf* N | putative periplasmic protein | S | MKKIFYGVIAFVVVLLIA | 55 | 90 | 26 | 255 | 64 | 490 | Core |
| Cj1221 | *gro* L | 60 kD chaperonin (cpn60) | O | MAKEIIFSDEARNKLYE | 55 | 90 | 26 | 255 | 64 | 490 | Core |

| | | | | | | | | | | | |
|---|---|---|---|---|---|---|---|---|---|---|---|
| Cj1226c | *cpr* S | putative two-component sensor (histidine kinase) | T | MNKSSIFYTITFIFIFAGV | 55 | 90 | 26 | 254 | 64 | 489 | Core |
| Cj1228c | *htr* A | serine protease (protease DO) | M | MKKIFLSLSLASALFAASI | 55 | 90 | 26 | 255 | 64 | 490 | Core |
| Cj1231 | *kef* B | putative glutathione-regulated potassium-efflux system protein | P | MDNFLEIFLITVAIAIVLN | 55 | 89 | 26 | 254 | 64 | 488 | Core |
| Cj1253 | *pnp* | polynucleotide phosphorylase | J | MQYSIEINKNTEIFDIDK | 55 | 90 | 25 | 253 | 64 | 487 | Core |
| Cj1257c | *mdt* G | putative efflux pump | EGP | MENFNRTLLVCWFGVF | 54 | 88 | 26 | 245 | 57 | 470 | Accessory |
| Cj1283 | *ktr* B | putative K+ uptake protein | P | MKQFGLDRRTFKILLAG | 55 | 90 | 26 | 255 | 64 | 490 | Core |
| Cj1290c | *acc* C | biotin carboxylase | I | MEIKSILIANRGEIALRAL | 55 | 90 | 26 | 255 | 64 | 490 | Core |
| Cj1311 | *pse* F | putative acylneuraminate cytidylyltransferase | M | MKNLCIIPARGGSKRIPR | 55 | 90 | 26 | 254 | 62 | 487 | Core |
| Cj1312 | *pse* G | nucleotidase specific for PseC product,UDP-4-amino-4,6-dideoxy-beta-L-AltNAc | M | MKVLFRSDSSSQIGFGH | 55 | 72 | 26 | 251 | 64 | 468 | Accessory |
| Cj0719c | *ygg* S | hypothetical protein | S | MTLEQILEKTKNIRLVAA | 55 | 90 | 26 | 255 | 64 | 490 | Core |
| Cj0718 | *dna* E | DNA polymerase III, alpha chain | L | MSQFTHLHLHTEYSLLD | 55 | 90 | 26 | 255 | 64 | 490 | Core |
| Cj0709 | *ffh* | signal recognition particle protein | U | MFELVSESFKSAINKLRF | 55 | 90 | 26 | 255 | 64 | 490 | Core |
| Cj0686 | *isp* G | 4-hydroxy-3-methylbut-2-en-1-yl diphosphate synthase | I | MEYKRFKTRQIKVGNVL | 55 | 90 | 26 | 255 | 64 | 490 | Core |
| Cj0685c | - | invasion phenotype protein | S | MQNLLLYIKNNLTPTLA | 15 | 39 | 10 | 58 | 13 | 135 | Accessory |
| A6J90_04300 | - | invasion phenotype protein | S | MGVYIDKNDFKQLEQN | 55 | 85 | 26 | 199 | 59 | 424 | Accessory |
| Cj0684 | *pri* A | putative primosomal protein N' | L | MQRSDIRYYELAICGLYL | 55 | 90 | 26 | 255 | 64 | 490 | Core |
| Cj0608 | *cus* C | putative outer membrane efflux protein | MU | MKIIFSIFLAFFLSACGAK | 55 | 90 | 26 | 255 | 64 | 490 | Core |
| Cj0607 | *mac* B | ABC-type transmembrane transport protein | V | MIFLKNICKNIGENAILK | 35 | 55 | 21 | 165 | 28 | 304 | Accessory |
| Cj0606 | *mac* A | amidohydrolase | M | MKKKVILIILIAILGSVGA | 35 | 55 | 21 | 163 | 24 | 298 | Accessory |
| Cj0605 | - | putative amidohydrolase | E | MQKLVENLALKYYDKV | 55 | 90 | 22 | 254 | 64 | 485 | Accessory |
| Cj0597 | *fba* A | Fructose-bisphosphate aldolase class II | G | MGVLDIVKAGVISGDEL | 55 | 90 | 26 | 255 | 64 | 490 | Core |
| Cj0586 | *lig* A | DNA ligase | L | MKKEEYLEKVALANLWI | 55 | 90 | 26 | 255 | 64 | 490 | Core |
| Cj0574 | *ilv* I | acetolactate synthase large subunit | H | MKELSGSAMICEALKEE | 55 | 90 | 26 | 255 | 64 | 490 | Core |
| Cj0557c | - | putative integral membrane protein | S | MLKQHFKEIAKLNSSEQ | 55 | 85 | 24 | 254 | 59 | 477 | Accessory |
| Cj0551 | *efp* | elongation factor P | J | MASYSMGDLKKGLKIEI | 55 | 90 | 26 | 255 | 64 | 490 | Core |
| Cj0545 | *ubi* D | putative 3-octaprenyl-4-hydroxybenzoate carboxy-lyase | H | MKEFIQILKENDLLRVIE | 55 | 90 | 26 | 255 | 64 | 490 | Core |
| Cj0531 | *icd* | isocitrate dehydrogenase, NADP-dependent | C | MQITYTLTDESPALATYS | 55 | 90 | 26 | 255 | 64 | 490 | Core |
| Cj0530 | - | putative periplasmic protein | M | MKKKILYIVIFFVVLILALF | 55 | 90 | 26 | 254 | 63 | 488 | Core |

| | | | | | | | | | | | |
|---|---|---|---|---|---|---|---|---|---|---|---|
| Cj0525c | *fts* I | putative penicillin-binding protein | M | MQEYKKNRVSKVAFAY | 55 | 90 | 26 | 255 | 64 | 490 | Core |
| Cj0518 | *htp* G | hsp90 family heat shock protein | O | MQFQTEVNQLLQLMIH | 55 | 90 | 26 | 255 | 64 | 490 | Core |
| Cj0506 | *ala* S | alanyl-tRNA synthetase | J | MDIRKAYLDFFASKRHE | 55 | 90 | 26 | 255 | 64 | 490 | Core |
| Cj0503c | *hem* H | ferrochelatase | H | MKLVLFLNMGGATNLQ | 55 | 90 | 26 | 255 | 64 | 490 | Core |
| Cj0500 | *sel* U | tRNA 2-selenouridine synthase | H | MLSEVEFTEFQKENFSLI | 31 | 24 | 18 | 83 | 59 | 215 | Accessory |
| Cj0499 | *hit* | putative histidine triad (HIT) family protein | FG | MQYLYAPWRSEYFEKVI | 31 | 25 | 18 | 84 | 59 | 217 | Accessory |
| Cj0479 | *rpo* C | DNA-directed RNA polymerase beta' chain | K | MSKFKVIEIKEDARPRDF | 55 | 90 | 26 | 255 | 64 | 490 | Core |
| Cj0465c | *ctb* | group III truncated haemoglobin | S | MKFETINQESIAKLMEIF | 55 | 90 | 26 | 255 | 64 | 490 | Core |
| Cj0464 | *rec* G | ATP-dependent DNA helicase | L | MKIKESDYEFFKKLKIRSA | 55 | 89 | 26 | 250 | 63 | 483 | Accessory |
| Cj0463 | *ymx* G | zinc protease-like protein | S | MQYLESRGVKIPFIFEKN | 55 | 90 | 26 | 255 | 64 | 490 | Core |
| Cj0462 | *mqn* C | putative radical SAM domain protein | H | MKRLDKKEALDLLHHAS | 55 | 90 | 26 | 255 | 64 | 490 | Core |
| Cj0444 | *cir* A_3 | TonB-dependent receptor, putative, degenerate | P | MILNIKIIIFKGINLTQIKN | 55 | 89 | 26 | 254 | 63 | 487 | Core |
| Cj1576c | *nuo* D | NADH dehydrogenase I chain D | C | MQIPSKLKPYYENIAFEC | 55 | 90 | 26 | 255 | 64 | 490 | Core |
| A6J90_00190 | - | putative protein | - | MLENLKEIYEEFELLIAQI | 0 | 0 | 25 | 0 | 0 | 25 | Accessory |
| A6J90_00195 | - | FIG00470712: hypothetical protein | S | MENDIIKKFINTDYKLLEI | 0 | 0 | 26 | 0 | 0 | 26 | Accessory |
| A6J90_00200 | - | FIG00471113: hypothetical protein | - | MDNKKLVINPKKILQNL | 1 | 0 | 26 | 0 | 0 | 27 | Accessory |
| Cj0571 | - | transcriptional regulator | K | MLSEIQELYPKLHKDFLL | 0 | 0 | 18 | 0 | 0 | 18 | Accessory |
| A6J90_00270 | - | putative protein | - | MTYQNFKNKLDTKIFGE | 0 | 0 | 26 | 0 | 0 | 26 | Accessory |
| A6J90_00275 | *ccr* M | DNA methylase | L | MQKDIIFQGNCLEILKTII | 0 | 0 | 26 | 0 | 0 | 26 | Accessory |
| Cj1612 | *prf* A | peptide chain release factor 1 | J | MLASKLDPFLKRFEELNS | 55 | 90 | 26 | 255 | 64 | 490 | Core |
| Cj1630 | *ton* B | putative TonB transport protein | U | MKTLFLNHKYQASYITFI | 17 | 19 | 25 | 168 | 13 | 242 | Accessory |
| Cj1631c | - | FIG00469530: hypothetical protein | - | MDKLTINDFNVSLPSSQ | 53 | 90 | 26 | 255 | 54 | 478 | Accessory |
| Cj1633 | *til* S | thiamine biosynthesis protein:ExsB | D | MKALALFSGGLDSMLAI | 55 | 90 | 26 | 255 | 64 | 490 | Core |
| Cj1634c | *aro* C | chorismate synthase | E | MNTFGTRLKFTSFGESH | 55 | 90 | 26 | 255 | 64 | 490 | Core |
| Cj0274 | *lpx* A | acyl-[acyl-carrier-protein]--UDP-N- acetylglucosamine O-acyltransferase | M | MKKIHPSAVIEEGAQLG | 55 | 90 | 26 | 255 | 64 | 490 | Core |
| Cj0279 | *car* B | carbamoyl-phosphate synthase large chain | F | MPKRTDVKSILLIGSGPIV | 55 | 90 | 26 | 255 | 64 | 490 | Core |
| A6J90_02340 | - | putative protein | - | MQNHLNREIMINLGSYY | 0 | 0 | 25 | 0 | 0 | 25 | Accessory |
| A6J90_02350 | - | R Pab1 restriction endonuclease | L | MNFKIDYELPLTSVAGKI | 0 | 0 | 25 | 0 | 0 | 25 | Accessory |
| A6J90_02350 | *sua* 5 | hypothetical protein | J | MIYLAQTDTTAGFLSKD | 0 | 0 | 26 | 0 | 0 | 26 | Accessory |

| | | | | | | | | | | | |
|---|---|---|---|---|---|---|---|---|---|---|---|
| A6J90_02420 | *pgt* P | MFS transporter%2C OPA family%2C phosphoglycerate transporter protein | G | MALGLILCALVNVLLGFS | 33 | 29 | 26 | 104 | 52 | 244 | Accessory |
| Cj0293 | *sur* E | multifunctional protein SurE | S | MKEILITNDDGYESEGLK | 55 | 90 | 26 | 255 | 64 | 490 | Core |
| Cj0300c | *mod* C | putative molybdenum transport ATP-binding protein | P | MIKIDINHPMNTAKGRL | 40 | 61 | 26 | 255 | 54 | 436 | Accessory |
| Cj0321 | *dxs* | L-deoxy-D-xylulose-5-phosphate synthase | H | MSKKFAHTQEELEKLSLI | 55 | 90 | 26 | 254 | 64 | 489 | Core |
| Cj0328c | *fab* H | 3-oxoacyl-[acyl-carrier-protein] synthase | I | MLKAGLKSIASYIPEKILS | 55 | 90 | 26 | 255 | 64 | 490 | Core |
| Cj0329c | *pls* X | putative fatty acid/phospholipid synthesis protein | I | MINIAIDAMGGDFGEKP | 55 | 90 | 26 | 255 | 64 | 490 | Core |
| Cj0352 | *fli* Z | putative transmembrane protein | N | MRLLVLFFLILPLYSVELIS | 55 | 90 | 26 | 255 | 64 | 490 | Core |
| Cj0356c | *fol* B | putative dihydroneopterin aldolase | H | MQSHIKIKFHFKCIIGILD | 55 | 90 | 26 | 255 | 64 | 490 | Core |
| Cj0428 | - | hypothetical protein | - | MQVNYRTISSYEYDAISC | 55 | 90 | 26 | 255 | 64 | 490 | Core |
| A6J90_00035 | *mlo* A | MloA protein, putative | S | MKEFIPSKLPLNIELNTAI | 2 | 18 | 12 | 8 | 0 | 40 | Accessory |
| Cj1543 | *kip* A | hypothetical protein | E | MSIKIIEASINSSLQDFGR | 52 | 86 | 20 | 235 | 52 | 445 | Accessory |
| A6J90_08990 | *hsd* R | type I restriction enzyme EcoR124II R protein | V | MNNLMTQNDNTTIITEY | 0 | 1 | 26 | 0 | 0 | 27 | Accessory |
| Cj1500 | *yed* E | putative integral membrane protein | S | MNFFKQKYLINFWDNS | 55 | 90 | 26 | 255 | 64 | 490 | Core |
| Cj1484c | - | putative membrane protein | - | MKKILQDGFFLAIFFFVL | 55 | 90 | 25 | 255 | 64 | 489 | Core |
| Cj1482c | *add* B | hypothetical protein | L | MKLRIFSSSRQIREYYNQ | 55 | 90 | 26 | 255 | 64 | 490 | Core |
| Cj1481c | *add* A | putative helicase | L | MSQFEPFLALEASAGSG | 55 | 90 | 26 | 255 | 64 | 490 | Core |
| Cj1476c | *nif* J | pyruvate-flavodoxin oxidoreductase | C | MGKIMKTMDGNEAAA | 55 | 90 | 26 | 255 | 64 | 490 | Core |
| Cj1457c | *tru* D | tRNA pseudouridine synthase D | J | MDLAEENTIFKPLYSLKH | 55 | 90 | 26 | 255 | 64 | 490 | Core |
| Cj0574 | *ilv* A | threonine dehydratase biosynthetic | E | MLELNKIYKAKQQISGFV | 55 | 90 | 26 | 255 | 64 | 490 | Core |
| Cj0886c | *fts* K | putative cell division protein | D | MLAPSMGEWVYKANLI | 55 | 90 | 26 | 255 | 64 | 490 | Core |
| Cj0887c | *flg* L | putative flagellin | N | MRITNKLNFTNSVNNSN | 55 | 90 | 26 | 255 | 57 | 483 | Accessory |
| Cj0891c | *ser* A | D-3-phosphoglycerate dehydrogenase | E | MKKKIIVCDAILDKGVDI | 55 | 90 | 26 | 255 | 64 | 490 | Core |
| Cj0924c | *che* B | putative MCP protein-glutamate methylesterase | NT | MKLILIGSSTGGPNQLKP | 55 | 90 | 26 | 255 | 64 | 490 | Core |
| Cj0929 | *pep* A | aminopeptidase | E | MKFELNDKKLDAIKADF | 55 | 90 | 26 | 255 | 63 | 489 | Core |
| Cj0930 | *ych* F | putative GTP-binding protein | J | MSLSVGIVGLPNVGKST | 55 | 90 | 26 | 255 | 64 | 490 | Core |
| Cj0931c | *arg* H | argininosuccinate lyase | E | MKNEMWSGRFSDASD | 53 | 86 | 26 | 229 | 53 | 447 | Accessory |
| A6J90_01640 | - | hypothetical protein | - | MINSLNSNLNYDYNTSN | 0 | 0 | 26 | 0 | 0 | 26 | Accessory |

| | | | | | | | | | | | |
|---|---|---|---|---|---|---|---|---|---|---|---|
| Cj0107 | *atp* D | ATP synthase F1, beta subunit | F | MQGFISQVLGPVVDVD | 55 | 90 | 26 | 255 | 64 | 490 | Core |
| Cj0105 | *atp* A | ATP synthase F1 sector alpha subunit | C | MKFKADEISSIIKERIENF | 55 | 90 | 26 | 255 | 64 | 490 | Core |
| Cj0101 | *par* B | parB family protein | K | MGLNKDRGLSSLISDMI | 55 | 90 | 26 | 254 | 64 | 489 | Core |
| A6J90_01500/A6J90_0150 | - | putative protein | V | MPKNDSFFNQVIKAKID | 0 | 0 | 25 | 0 | 0 | 25 | Accessory |
| A6J90_01490 | - | putative protein | - | MSDLNKIPKETKARQTG | 0 | 0 | 26 | 0 | 0 | 26 | Accessory |
| Cj0077c | *cdt* C | cytolethal distending toxin%2C subunit C | S | MIKIWNIKEIVLSDELKQ | 0 | 0 | 26 | 4 | 0 | 30 | Accessory |
| Cj0076c | *lld* P | L-lactate permease | C | MEQILTWQQIYDPFSNI | 55 | 90 | 26 | 255 | 64 | 490 | Core |
| A6J90_01375 | - | Non-heme iron protein, hemerythrin family | P | MIDIQHQKLFELAGKVE | 0 | 0 | 12 | 0 | 0 | 12 | Accessory |
| A6J90_01275 | - | anion transporter | P | MLLLLWAGALGLFFGISI | 12 | 77 | 23 | 113 | 17 | 242 | Accessory |
| Cj0038c | - | putative poly(A) polymerase family protein | - | MDLKSLENNRLYILKRLG | 49 | 81 | 25 | 241 | 45 | 441 | Accessory |
| Cj1414c | *kps* C | capsule polysaccharide export protein KpsC | M | MNLAKKHKAKFILLEDG | 4 | 0 | 26 | 2 | 1 | 33 | Accessory |
| Cj0943 | *lol* A | putative outer-membrane lipoprotein carrier protein precursor | M | MKKTFLIFFIFIGELFALD | 55 | 90 | 26 | 255 | 64 | 490 | Core |
| Cj0944c | - | putative periplasmic protein | - | MKKIFLSVFLVLSLNAQN | 55 | 90 | 25 | 255 | 62 | 487 | Core |
| Cj0945c | - | putative helicase | L | MLDKLEKILAYDNVFLSC | 55 | 90 | 23 | 252 | 64 | 484 | Accessory |
| Cj0946 | *spr* 7 | putative lipoprotein | M | MKSCLYFTFIVLFLTACST | 55 | 90 | 26 | 255 | 64 | 490 | Core |
| Cj0995c | *hem* B | delta-aminolevulinic acid dehydratase | H | MFKRFRRLRLNENLRAN | 55 | 90 | 26 | 255 | 64 | 490 | Core |
| Cj1011 | *cor* A | putative CorA-like Mg2+ transporter protein | P | MLELHENLKKILQAKNLI | 55 | 90 | 26 | 255 | 64 | 490 | Core |
| Cj0773c | *met* I | putative ABC transport system permease protein | P | MNEENISIISAFFSRISQF | 55 | 90 | 26 | 255 | 64 | 490 | Core |
| A6J90_04615 | - | hypothetical protein | - | MDHILSGKAENFVAGW | 17 | 14 | 24 | 57 | 49 | 161 | Accessory |
| Cj0796c | *mhp* C | putative hydrolase | S | MAQTQLSYKNKTYQISY | 55 | 90 | 26 | 255 | 64 | 490 | Core |

[a]Locus tag for accessory genes based on C. jejuni reference genome CP022076.1 (NCBI accession). Locus tags for allelic variants of the core genome refer to C. jejuni strain NCTC11168 (NCBI accession: AL111168.1)

[b]clusters of orthologous groups (http://clovr.org/docs/clusters-of-orthologous-groups-cogs/);

[c]number of genomes assigned to a particular lifestyle carrying the gene or allelic variant (pig, cattle, chicken, host generalists, others)

[d]indicates that a gene belongs to the accessory (A) or the core (C) genome content of C. jejuni
