## Supplementary material for "Genome-wide insights into population structure and host specificity of *Campylobacter jejuni*": Table S6

Table S6: Overview of host-generalist associated accessory genes and core gene variants identified by k-mer mapping.

| Reference Gene Nane[a] | Alias | Predicted Function | COG Family[b] | AA | # Cattle[c] | # Chicken[c] | # Pig[c] | # Generalist[c] | # Others[c] | Total | Core/Accessory[d] |
|---|---|---|---|---|---|---|---|---|---|---|---|
| Cj1342c | - | no Hp match. A member of the 617 family of C.j. proteins containing homopolymeric tracts | E | MIVSKAYEIDSCDDVE | 12 | 39 | 25 | 167 | 30 | 273 | Accessory |
| Cj1341c | *pse* E | Protein of unknown function DUF115 | S | MTILEKNIQALLSGVN | 1 | 33 | 2 | 146 | 17 | 199 | Accessory |
| Cj1276c | *fts* X | Cell division protein FtsX | D | MKFFKTHLSLILPLLFN | 55 | 90 | 26 | 255 | 64 | 490 | Core |
| Cj1266c | *hyd* B | Belongs to the NiFe NiFeSe hydrogenase large subunit family | C | MSQKIIVDPITRIEGHI | 55 | 90 | 26 | 255 | 64 | 490 | Core |
| Cj1252 | *lpt* D | Involved in the assembly of lipopolysaccharide (LPS) at the surface of the outer membrane | M | MWRKFSLLLGTSIALN | 55 | 90 | 25 | 255 | 64 | 489 | Core |
| Cj1250 | *pur* D | Belongs to the GARS family | F | MKIMILGSGAREYSIA | 55 | 90 | 26 | 255 | 64 | 490 | Core |
| Cj1240c | - | - | - | MKKIILLLGLSLASLGA | 55 | 86 | 21 | 249 | 64 | 475 | Accessory |
| Cj1019c | *liv* J | amino acid abc transporter | E | MKKLTLTLSVLTMVN | 55 | 90 | 26 | 255 | 64 | 490 | Core |
| Cj1008c | *aro* B | Catalyzes the conversion of 3-deoxy-D-arabino- heptulosonate 7-phosphate (DAHP) to dehydroquinate (DHQ) | E | MQVEVKLKENAYKVY | 55 | 90 | 26 | 255 | 63 | 489 | Core |
| Cj1001 | *rpo* D | Sigma factors are initiation factors that promote the attachment of RNA polymerase to specific initiation sites and are then released. This sigma factor is the primary sigma factor during exponential growth | K | MNAKTQEAELEELFQ | 55 | 90 | 26 | 255 | 64 | 490 | Core |
| Cj0975 | *hxu* B | Haemolysin secretion/activation protein ShlB/FhaC/HecB | U | MLEQSPYKEDANSKN | 37 | 90 | 25 | 195 | 43 | 390 | Accessory |
| Cj0964 | - | leucine binding | E | MKKIFLIFFLSCFLLNA | 55 | 90 | 26 | 255 | 64 | 490 | Core |
| Cj0961c | *rpm* H | Belongs to the bacterial ribosomal protein bL34 family | J | MKRTYQPHGTPRKRT | 55 | 90 | 26 | 255 | 64 | 490 | Core |
| Cj0960c | *rnp* A | RNaseP catalyzes the removal of the 5'-leader sequence from pre-tRNA to produce the mature 5'-terminus. It can also cleave other RNA substrates such as 4.5S RNA. The protein component plays an auxiliary but essential role in vivo by binding to the 5'-leader sequence and broadening the substrate specificity of the ribozyme | J | MKNFDKFSTNEEFSSY | 55 | 90 | 26 | 255 | 64 | 490 | Core |
| Cj0958c | *yid* C | Required for the insertion and or proper folding and or complex formation of integral membrane proteins into the membrane. Involved in integration of membrane proteins that insert both dependently and independently of the Sec translocase complex, as well as at least some lipoproteins. Aids folding of multispanning membrane proteins | U | MNNSNNIFQQKRILL | 55 | 90 | 26 | 255 | 64 | 490 | Core |
| Cj0956c | *mnm* E | Exhibits a very high intrinsic GTPase hydrolysis rate. Involved in the addition of a carboxymethylaminomethyl (cmnm) group at the wobble position (U34) of certain tRNAs, forming tRNA- cmnm(5)s(2)U34 | J | MSDTIAAIATAHGVG | 55 | 90 | 26 | 255 | 64 | 490 | Core |
| Cj0945c | - | COG0507 ATP-dependent exoDNAse (exonuclease V) alpha subunit - helicase superfamily I member | L | MLDKLEKILAYDNVFL | 55 | 90 | 23 | 252 | 64 | 484 | Accessory |
| Cj0943 | *lol* A | Participates in the translocation of lipoproteins from the inner membrane to the outer membrane. Only forms a complex with a lipoprotein if the residue after the N-terminal Cys is not an aspartate (The Asp acts as a targeting signal to indicate that the lipoprotein should stay in the inner membrane) | M | MKKTFLIFFIFIGQLFA | 55 | 90 | 26 | 255 | 64 | 490 | Core |
| Cj0934c | IV02_29000 | Belongs to the sodium neurotransmitter symporter (SNF) (TC 2.A.22) family | P | MRTYFSKIGFVLTVAG | 32 | 81 | 0 | 175 | 15 | 303 | Accessory |
| Cj0932c | *pck* A | Involved in the gluconeogenesis. Catalyzes the conversion of oxaloacetate (OAA) to phosphoenolpyruvate (PEP) through direct phosphoryl transfer between the nucleoside triphosphate and OAA | H | MKKFDKLGLDNIKEIF | 55 | 90 | 26 | 255 | 64 | 490 | Core |
| Cj0917c | *cst* A | Carbon starvation protein | T | MTQLSTKILWLFVAA | 52 | 88 | 26 | 245 | 63 | 474 | Accessory |
| Cj0915 | *yci* A | hmm pf03061 | I | MRDMGEPKLKIVAM | 55 | 90 | 26 | 255 | 64 | 490 | Core |
| Cj0912c | *cys* M | Belongs to the cysteine synthase cystathionine beta- synthase family | E | MKVHEKVSELIGNTP | 52 | 88 | 26 | 249 | 64 | 479 | Accessory |
| Cj0911 | *hya* E | SCO1 SenC | S | MKKNIILFIVIVAIILGV | 55 | 90 | 26 | 255 | 64 | 490 | Core |

| | | | | | | | | | | | |
|---|---|---|---|---|---|---|---|---|---|---|---|
| Cj0909 | VY92_09940 | hmm pf04314 | S | MKKILLLGALFAVNLV | 55 | 89 | 26 | 252 | 64 | 486 | Core |
| Cj0905c | *alr* | Catalyzes the interconversion of L-alanine and D- alanine. May also act on other amino acids | E | MSLIKIDQKAYEYNLR | 55 | 90 | 26 | 255 | 64 | 490 | Core |
| Cj0898 | *hin* T | Hit family | FG | MQEKTIFELIVEGKLP | 55 | 90 | 26 | 254 | 64 | 489 | Core |
| Cj0886c | *fts* K | Belongs to the FtsK SpoIIIE SftA family | D | MLAPGMGEWVYKA | 55 | 90 | 26 | 255 | 64 | 490 | Core |
| Cj0879c | - | - | - | MKKCILIFFSLYSLSFA | 38 | 57 | 4 | 140 | 57 | 296 | Accessory |
| Cj0874c | *pet* A | cytochrome C | C | MNKFSIVLTLLLCGSC | 52 | 73 | 26 | 239 | 54 | 444 | Accessory |
| Cj0874c | *pe* tA | cytochrome C | C | MDKCAVCHQENGLG | 0 | 24 | 0 | 87 | 7 | 118 | Accessory |
| Cj0866 | - | hmm pf05935 | M | MRLSKTLCMALLACS | 0 | 37 | 0 | 97 | 0 | 134 | Accessory |
| Cj0865 | *dsb* B | Required for disulfide bond formation in some proteins. Part of a redox system composed of DsbI and DsbL that mediates formation of an essential disulfide bond in AssT (By similarity) | C | MKDKCRNFSLSKWQ | 55 | 90 | 26 | 254 | 64 | 489 | Core |
| Cj0835c | *acn* B | Belongs to the aconitase IPM isomerase family | C | MSFMQEYNKLVEERA | 55 | 90 | 26 | 255 | 64 | 490 | Core |
| Cj0812 | *thr* C | threonine synthase | E | MKLVESRNVNNVSSF | 55 | 80 | 26 | 255 | 64 | 480 | Accessory |
| Cj0800c | - | Flagellar Assembly Protein A | L | MALDEKIIAYTENPAR | 55 | 90 | 26 | 255 | 64 | 490 | Core |
| Cj0799c | *ruv* A | The RuvA-RuvB complex in the presence of ATP renatures cruciform structure in supercoiled DNA with palindromic sequence, indicating that it may promote strand exchange reactions in homologous recombination. RuvAB is a helicase that mediates the Holliday junction migration by localized denaturation and reannealing. RuvA stimulates, in the presence of DNA, the weak ATPase activity of RuvB | L | MVVGIEGIITKKEPTFI | 55 | 90 | 26 | 255 | 64 | 490 | Core |
| Cj0798c | *ddl* | Belongs to the D-alanine--D-alanine ligase family | F | MKFAILFGGNSYEHEI | 55 | 90 | 26 | 255 | 64 | 490 | Core |
| Cj0794 | - | Annotation was generated automatically without manual curation | S | MSIFSINDNSNYGSILS | 48 | 76 | 19 | 174 | 22 | 339 | Accessory |
| Cj0791c | *csd* A | Aminotransferase | E | MQISDLKKELILKKGIL | 55 | 90 | 26 | 255 | 63 | 489 | Core |
| Cj0780 | *nap* A | nitrate reductase (NAP). Only expressed at high levels during aerobic growth. NapAB complex receives electrons from the membrane-anchored tetraheme protein NapC | C | MNRRDFIKNTAIASA | 55 | 90 | 26 | 254 | 64 | 489 | Core |
| Cj0776c | - | - | - | MNLKIFSIIVGILIAVIV | 55 | 90 | 26 | 255 | 64 | 490 | Core |
| Cj0775c | *val* S | amino acids such as threonine, to avoid such errors, it has a posttransfer editing activity that hydrolyzes mischarged Thr-tRNA(Val) in a tRNA-dependent manner | J | MYDKNLEKEYYQICEI | 55 | 90 | 26 | 255 | 64 | 490 | Core |
| Cj0737 | - | haemagglutination activity domain | U | MKKMSKHIVLSFAVS | 51 | 77 | 15 | 195 | 34 | 372 | Accessory |
| Cj0718 | *dna* E | DNA polymerase | L | MSQFTHLHLHTEYSLI | 55 | 90 | 26 | 255 | 64 | 490 | Core |
| Cj0717 | *spx* A | Belongs to the ArsC family | P | MKLYGIKNCNSVKKA | 55 | 90 | 26 | 255 | 64 | 490 | Core |
| Cj0710 | *rps* P | Belongs to the bacterial ribosomal protein bS16 family | J | MTVIRLTRMGRTKRP | 55 | 90 | 26 | 255 | 64 | 490 | Core |
| Cj0709 | ffh | Involved in targeting and insertion of nascent membrane proteins into the cytoplasmic membrane. Binds to the hydrophobic signal sequence of the ribosome-nascent chain (RNC) as it emerges from the ribosomes. The SRP-RNC complex is then targeted to the cytoplasmic membrane where it interacts with the SRP receptor FtsY. Interaction with FtsY leads to the transfer of the RNC complex to the Sec translocase for insertion into the membrane, the hydrolysis of GTP by both Ffh and FtsY, and the dissociation of the SRP-FtsY complex into the individual components | U | MFELVSESFKSAINKL | 55 | 90 | 26 | 255 | 64 | 490 | Core |
| Cj0703 | - | Protein of unknown function (DUF3972) | S | MQTYLELEEFCKLVHI | 55 | 90 | 26 | 255 | 64 | 490 | Core |
| Cj0700 | - | - | - | MTQEELDALMNGDV | 55 | 90 | 26 | 255 | 64 | 490 | Core |
| Cj0699c | *gln* A | glutamine synthetase | E | MGKFVNNIDDFFKFC | 55 | 90 | 26 | 255 | 64 | 490 | Core |

| | | | | | | | | | | | |
|---|---|---|---|---|---|---|---|---|---|---|---|
| Cj0684 | *pri* A | Involved in the restart of stalled replication forks. Recognizes and binds the arrested nascent DNA chain at stalled replication forks. It can open the DNA duplex, via its helicase activity, and promote assembly of the primosome and loading of the major replicative helicase DnaB onto DNA | L | MQRSDIRYYELAICGL | 55 | 90 | 26 | 255 | 64 | 490 | Core |
| Cj0680c | *uvr* B | damaged site, the DNA wraps around one UvrB monomer. DNA wrap is dependent on ATP binding by UvrB and probably causes local melting of the DNA helix, facilitating insertion of UvrB beta-hairpin between the DNA strands. Then UvrB probes one DNA strand for the presence of a lesion. If a lesion is found the UvrA subunits dissociate and the UvrB-DNA preincision complex is formed. This complex is subsequently bound by UvrC and the second UvrB is released. If no lesion is found, the DNA wraps around the other UvrB subunit that will check the other stand for damage | L | MLELTSEFKPSPDQQ | 55 | 90 | 26 | 255 | 64 | 490 | Core |
| Cj0628 | - | Autotransporter beta-domain | S | MTDSASLSGEMILSG | 0 | 21 | 0 | 13 | 0 | 34 | Accessory |
| Cj0506 | *ala* S | Catalyzes the attachment of alanine to tRNA(Ala) in a two-step reaction alanine is first activated by ATP to form Ala- AMP and then transferred to the acceptor end of tRNA(Ala). Also edits incorrectly charged Ser-tRNA(Ala) and Gly-tRNA(Ala) via its editing domain | J | MDIRKAYLDFFASKGI | 55 | 90 | 26 | 255 | 64 | 490 | Core |
| Cj0496 | - | Uncharacterised protein family (UPF0153) | S | MIFDKNFSYAFDENA | 55 | 90 | 26 | 255 | 64 | 490 | Core |
| Cj0493 | *fus* A | Catalyzes the GTP-dependent ribosomal translocation step during translation elongation. During this step, the ribosome changes from the pre-translocational (PRE) to the post- translocational (POST) state as the newly formed A-site-bound peptidyl-tRNA and P-site-bound deacylated tRNA move to the P and E sites, respectively. Catalyzes the coordinated movement of the two tRNA molecules, the mRNA and conformational changes in the ribosome | J | MSRSTPLKKVRNIGIA | 55 | 90 | 26 | 255 | 64 | 490 | Core |
| Cj0492 | *rps* G | One of the primary rRNA binding proteins, it binds directly to 16S rRNA where it nucleates assembly of the head domain of the 30S subunit. Is located at the subunit interface close to the decoding center, probably blocks exit of the E-site tRNA | J | MRRRKAPVREVLPDP | 55 | 90 | 26 | 255 | 64 | 490 | Core |
| Cj0491 | *rps* L | Interacts with and stabilizes bases of the 16S rRNA that are involved in tRNA selection in the A site and with the mRNA backbone. Located at the interface of the 30S and 50S subunits, it traverses the body of the 30S subunit contacting proteins on the other side and probably holding the rRNA structure together. The combined cluster of proteins S8, S12 and S17 appears to hold together the shoulder and platform of the 30S subunit | J | MPTINQLVRKERKKV | 55 | 90 | 26 | 255 | 64 | 490 | Core |
| Cj0490 | *ald* A | Belongs to the aldehyde dehydrogenase family | C | MTTYLNYIDGKFIPHN | 0 | 36 | 0 | 177 | 0 | 213 | Accessory |
| Cj0483 | *uxa* A | Altronate hydrolase | G | MKKIMGYRREDGKF( | 0 | 57 | 0 | 195 | 2 | 254 | Accessory |
| Cj0480c | - | Transcriptional regulator | K | MHQPTLRVLNILELLA | 0 | 57 | 0 | 194 | 0 | 251 | Accessory |
| Cj0479 | *rpo* C | DNA-dependent RNA polymerase catalyzes the transcription of DNA into RNA using the four ribonucleoside triphosphates as substrates | K | MSKFKVIEIKEDARPR | 55 | 90 | 26 | 255 | 64 | 490 | Core |
| Cj0478 | *rpo* B | DNA-dependent RNA polymerase catalyzes the transcription of DNA into RNA using the four ribonucleoside triphosphates as substrates | K | MLDNKLGNRLRVDFS | 55 | 90 | 26 | 255 | 64 | 490 | Core |
| Cj0477 | *rpl* L | Forms part of the ribosomal stalk which helps the ribosome interact with GTP-bound translation factors. Is thus essential for accurate translation | J | MAISKEDVLEYISNLSY | 55 | 90 | 26 | 255 | 64 | 490 | Core |
| Cj0475 | *rpl* A | Binds directly to 23S rRNA. The L1 stalk is quite mobile in the ribosome, and is involved in E site tRNA release | J | MAKIAKRLKELSQKID | 55 | 90 | 26 | 255 | 64 | 490 | Core |
| Cj0470 | *tuf* | This protein promotes the GTP-dependent binding of aminoacyl-tRNA to the A-site of ribosomes during protein biosynthesis | J | MAKEKFSRNKPHVNI | 55 | 90 | 26 | 255 | 64 | 490 | Core |
| Cj0464 | *rec* G | ATP-dependent DNA helicase | L | MKIKESDFEFFKKLKIF | 55 | 89 | 26 | 250 | 63 | 483 | Accessory |
| Cj0463 | *ymx* G | Peptidase, M16 | S | MQYLESRGIKIPFIFEK | 55 | 90 | 26 | 255 | 64 | 490 | Core |

| | | | | | | | | | | | |
|---|---|---|---|---|---|---|---|---|---|---|---|
| Cj0461c | *bac* E | Major facilitator Superfamily | EGP | MNYIELLKNNKNIRIL | 55 | 90 | 26 | 255 | 64 | 490 | Core |
| Cj0460 | *nus* A | Participates in both transcription termination and antitermination | K | MEKIADIIESIANEKNI | 55 | 90 | 26 | 255 | 64 | 490 | Core |
| Cj0459c | - | - | - | MELKLARTLINEKPKN | 55 | 90 | 26 | 255 | 64 | 490 | Core |
| Cj0458c | *mia* B | Catalyzes the methylthiolation of N6- (dimethylallyl)adenosine (i(6)A), leading to the formation of 2- methylthio-N6-(dimethylallyl)adenosine (ms(2)i(6)A) at position 37 in tRNAs that read codons beginning with uridine | J | MSAKKLFIQTLGCAM | 55 | 90 | 25 | 255 | 64 | 489 | Core |
| Cj0457c | MA20_05800 | protein conserved in bacteria | S | MGKSFKIHCLTYIIFIL | 55 | 90 | 26 | 255 | 64 | 490 | Core |
| Cj0453 | *thi* C | Catalyzes the synthesis of the hydroxymethylpyrimidine phosphate (HMP-P) moiety of thiamine from aminoimidazole ribotide (AIR) in a radical S-adenosyl-L-methionine (SAM)-dependent reaction | H | MKTQMNYAKEGIFTI | 54 | 90 | 26 | 252 | 64 | 486 | Core |
| Cj0452 | *dna* Q | dna polymerase iii | L | MSLQQIDQIISILNKQ | 53 | 90 | 26 | 255 | 64 | 488 | Core |
| Cj0451 | *rpe* | Belongs to the ribulose-phosphate 3-epimerase family | G | MYVAPSLLSANFLKLE | 55 | 90 | 26 | 255 | 64 | 490 | Core |
| Cj0444 | *cirA* _3 | receptor | P | MVALYGENEYFITDD | 35 | 66 | 0 | 174 | 6 | 281 | Accessory |
| Cj0434 | *gpm* I | Catalyzes the interconversion of 2-phosphoglycerate and 3-phosphoglycerate | G | MKQKCVLIITDGIGYN | 55 | 90 | 26 | 255 | 64 | 490 | Core |
| Cj0432c | *mur* D | Cell wall formation. Catalyzes the addition of glutamate to the nucleotide precursor UDP-N-acetylmuramoyl-L-alanine (UMA) | M | MKISLFGYGKTTRAIA | 55 | 90 | 26 | 255 | 64 | 490 | Core |
| Cj0431 | - | general secretion pathway protein | NU | MIKAFSLLEFVFIILILG | 55 | 90 | 26 | 255 | 64 | 490 | Core |
| Cj0431 | - | integral membrane protein | M | MEKIKNYKLIIILLSLDL | 55 | 90 | 26 | 255 | 64 | 490 | Core |
| Cj0429c | *yig* Z | hmm pf01205 | S | MQTIDQIFQTQIDIKK | 55 | 90 | 26 | 255 | 64 | 490 | Core |
| Cj0428 | - | - | - | MQVNYRTISSYEYDA | 55 | 90 | 26 | 255 | 64 | 490 | Core |
| Cj0426 | *ybi* T | abc transporter atp-binding protein | S | MVEVKNLTMRFANC | 55 | 90 | 26 | 255 | 64 | 490 | Core |
| Cj0422c | - | - | - | MTKKSKRDMAYELDI | 36 | 85 | 26 | 255 | 49 | 451 | Accessory |
| Cj0404 | - | Sporulation related domain | S | MENQKNEFDDIILEKS | 55 | 90 | 26 | 255 | 64 | 490 | Core |
| Cj0396c | *pgb* B | - | - | MKKIFLTLFCLIFLCAC | 55 | 90 | 26 | 255 | 63 | 489 | Core |
| Cj0393c | *mqo* | Malate quinone- oxidoreductase | C | MSQQEFDVLVIGAGI | 55 | 90 | 26 | 255 | 64 | 490 | Core |
| Cj0362 | - | membrane | S | MTEWINDYYFWIKW | 55 | 90 | 26 | 255 | 64 | 490 | Core |
| Cj0321 | *dxs* | Catalyzes the acyloin condensation reaction between C atoms 2 and 3 of pyruvate and glyceraldehyde 3-phosphate to yield 1-deoxy-D-xylulose-5-phosphate (DXP) | H | MSKKFAHTQEELEKLS | 55 | 90 | 26 | 254 | 64 | 489 | Core |
| Cj0318 | *fli* F | The M ring may be actively involved in energy transduction | N | MDFKNMLHQIGQLY | 55 | 90 | 26 | 255 | 64 | 490 | Core |
| Cj0292c | *pgt* P | Catalyzes the uptake of glycerol-3-phosphate into the cell with the simultaneous export of inorganic phosphate from the cell | G | MALGLILCALVNVLLG | 38 | 20 | 0 | 88 | 6 | 152 | Accessory |
| Cj0248 | - | Signal transduction protein | T | MIGDMNELLLKSVEV | 55 | 90 | 26 | 255 | 64 | 490 | Core |
| Cj0247c | - | FIST N domain | NT | MILFSEDMIENLCTNK | 54 | 90 | 26 | 240 | 55 | 465 | Accessory |
| Cj0196c | *pur* F | Catalyzes the formation of phosphoribosylamine from phosphoribosylpyrophosphate (PRPP) and glutamine | F | MCAVVGVINSKNAST | 55 | 90 | 26 | 255 | 64 | 490 | Core |
| Cj0192c | *clp* P | Cleaves peptides in various proteins in a process that requires ATP hydrolysis. Has a chymotrypsin-like activity. Plays a major role in the degradation of misfolded proteins | O | MFIPYVIEKSSRGERSY | 55 | 90 | 26 | 255 | 64 | 490 | Core |
| Cj0186c | *Ter* C | Membrane protein, TerC | P | MFEWIFSIDAWITLAT | 54 | 88 | 25 | 253 | 64 | 484 | Accessory |
| Cj0185c | *phn* A | Zn-ribbon-containing protein involved in phosphonate metabolism | P | MAKDANGIELNTGDS | 3 | 57 | 1 | 124 | 5 | 190 | Accessory |
| Cj0183 | *cor* C | COG1253 Hemolysins and related proteins containing CBS domains | S | MDPSQVLDLNQTSTA | 55 | 90 | 26 | 255 | 64 | 490 | Core |
| Cj0182 | *sbm* A | ABC transporter transmembrane region 2 | I | MFSSFFKSKKWALWA | 55 | 76 | 26 | 255 | 29 | 441 | Accessory |

| | | | | | | | | | | | |
|---|---|---|---|---|---|---|---|---|---|---|---|
| Cj0108 | *atp* C | Produces ATP from ADP in the presence of a proton gradient across the membrane | C | MNDLINFEIVTPLGVI | 55 | 90 | 26 | 255 | 64 | 490 | Core |
| Cj0105 | *atp* A | Produces ATP from ADP in the presence of a proton gradient across the membrane. The alpha chain is a regulatory subunit | C | MKFKADEISSIIKERIE | 55 | 90 | 26 | 255 | 64 | 490 | Core |
| Cj0100 | *par* A | involved in chromosome partitioning | D | MSEIITIANQKGGVG | 55 | 90 | 26 | 255 | 64 | 490 | Core |
| Cj0093 | - | curli production assembly transport component CsgG | M | MKIIKILFLGLFLSLSL | 55 | 88 | 26 | 214 | 42 | 425 | Accessory |
| Cj0089 | - | protein conserved in bacteria | S | MKIKVGLIFSGIACLF | 55 | 90 | 26 | 255 | 64 | 490 | Core |
| Cj0087 | *asp* A | Aspartate ammonia-lyase | E | MGTRKEHDFIGELEI | 55 | 90 | 26 | 255 | 64 | 490 | Core |
| Cj0086c | *ung* | Excises uracil residues from the DNA which can arise as a result of misincorporation of dUMP residues by DNA polymerase or due to deamination of cytosine | L | MEEITINIDKIKINDD | 55 | 90 | 26 | 255 | 64 | 490 | Core |
| Cj0082 | *cyd* B | cytochrome d ubiquinol oxidase, subunit II | C | MFFGLELEGLQIYW | 55 | 90 | 26 | 255 | 64 | 490 | Core |
| Cj0076c | *lld* P | L-lactate permease | C | MEQILTWQQIYDPF | 54 | 87 | 1 | 254 | 64 | 460 | Accessory |
| Cj0036 | - | protein conserved in bacteria | S | MQNLNQNEQIKCPS | 53 | 90 | 26 | 252 | 62 | 483 | Accessory |
| Cj1713 | *rlm* N | Specifically methylates position 2 of adenine 2503 in 23S rRNA and position 2 of adenine 37 in tRNAs | J | MKELVNILDFLPEELG | 55 | 90 | 26 | 255 | 64 | 490 | Core |
| Cj1703c | *rps* S | Protein S19 forms a complex with S13 that binds strongly to the 16S ribosomal RNA | J | MARSLKKGPFVDDH | 55 | 90 | 26 | 255 | 64 | 490 | Core |
| Cj1702c | *rpl* V | The globular domain of the protein is located near the polypeptide exit tunnel on the outside of the subunit, while an extended beta-hairpin is found that lines the wall of the exit tunnel in the center of the 70S ribosome | J | MSKALIKFIRLSPTKA | 55 | 90 | 26 | 255 | 64 | 490 | Core |
| Cj1701c | *rps* C | Binds the lower part of the 30S subunit head. Binds mRNA in the 70S ribosome, positioning it for translation | J | MGQKVNPIGLRLGIN | 55 | 90 | 26 | 255 | 64 | 490 | Core |
| Cj1690c | *rps* E | Located at the back of the 30S subunit body where it stabilizes the conformation of the head with respect to the body | J | MEKYNREEFEEVIVD | 55 | 90 | 26 | 255 | 64 | 490 | Core |
| Cj0628 | - | Autotransporter beta-domain | S | MGGGKDSSKSIISNF | 0 | 0 | 0 | 5 | 0 | 5 | Accessory |
| Cj1669c | *lig* | DNA ligase | L | MRFVFLICCACLVFA | 55 | 90 | 26 | 255 | 64 | 490 | Core |
| Cj1474c | *cts* D | Type II and III secretion system protein | NU | MIRLILINILFCHYLYA | 51 | 83 | 24 | 233 | 45 | 436 | Accessory |
| GRN82_06795 | - | Protein of unknown function (DUF2972) | S | MHKPNSAIERIKNHL | 0 | 3 | 0 | 20 | 0 | 23 | Accessory |

[a]Locus tags of genes are named by homlogous genens from the reference strain NCTC11168 (NCBI accession: AL111168.1) and CFSAN096296 (NCBI accession: CP047484)

[b]Clusters of Orthologous Groups

[c]Number of geneomes of a catrain group that carrie the gene (pig, cattle, chicken, generalists, others)

[d]Notes of the corresponding gene belongs to the accessory (A) or core genome (C ) content
